# Recovering signals of ghost archaic introgression in African populations

**DOI:** 10.1101/285734

**Authors:** Arun Durvasula, Sriram Sankararaman

## Abstract

While introgression from Neanderthals and Denisovans has been well-documented in modern humans outside Africa, the contribution of archaic hominins to the genetic variation of present-day Africans remains poorly understood. Using 405 whole-genome sequences from four sub-Saharan African populations, we provide complementary lines of evidence for archaic introgression into these populations. Our analyses of site frequency spectra indicate that these populations derive 2-19% of their genetic ancestry from an archaic population that diverged prior to the split of Neanderthals and modern humans. Using a method that can identify segments of archaic ancestry without the need for reference archaic genomes, we built genome-wide maps of archaic ancestry in the Yoruba and the Mende populations that recover about 482 and 502 megabases of archaic sequence, respectively. Analyses of these maps reveal segments of archaic ancestry at high frequency in these populations that represent potential targets of adaptive introgression. Our results reveal the substantial contribution of archaic ancestry in shaping the gene pool of present-day African populations.

**One sentence summary:** Multiple present-day African populations inherited genes from an unknown archaic population that diverged before modern humans and Neanderthals split.

## Main text

Admixture has been a dominant force in shaping patterns of genetic variation in human populations (*1*). Comparisons of genome sequences from archaic hominins to those from present-day humans have documented multiple interbreeding events including gene flow from Neanderthals into the ancestors of all non-Africans (*2*), from Denisovans into Oceanians (*3*) and eastern non-Africans (*4, 5*), as well as from early modern humans into the Neanderthals (*6*). However, the sparse fossil record and the difficulty in obtaining ancient DNA have made it challenging to dissect the contribution of archaic hominins to genetic diversity within Africa. While several studies have revealed contributions from deep lineages to the ancestry of present-day Africans (*7, 8, 9, 10, 11, 12*), the nature of these contributions remains poorly understood.

We leveraged whole genome sequence data from present-day African populations as well as archaic hominins to compute statistics that are sensitive to introgression in the history of African populations. Specifically, we tabulated the distribution of the frequencies of derived alleles (where a derived allele is determined relative to an inferred human ancestor) in African populations at single nucleotide polymorphisms (SNPs) for which a randomly sampled allele from an archaic individual was observed to also be derived. Theory predicts that this conditional site frequency spectrum (CSFS) is expected to be uniformly distributed when alleles are neutrally evolving under a demographic model in which the ancestor of modern and archaic humans, assumed to be at mutation-drift equilibrium, split with no subsequent gene flow between the two groups (*13, 14*). Importantly, this expectation is robust to assumptions about changes in population sizes in the history of modern human or archaic populations. Further, we show that this expectation holds even when there is population structure or gene flow in the history of the archaic population (SI Section S1).

We computed CSFS_YRI,N_: the CSFS in the Yoruba from Ibadan (YRI) while restricting to SNPs where a randomly sampled allele from the high-coverage Vindija Neanderthal (N) genome was observed to be derived (*15*). In contrast to the uniform spectrum expected from theory, we observe that the CSFS_YRI,N_ has a U-shape with an elevated proportion of SNPs with low and high frequency derived alleles relative to those at intermediate frequencies (Figure 1, Figure S4). The CSFS is nearly identical when we replace the Vindija Neanderthal genome with the high-coverage Denisova genome (*4*) (Figure 1, Figure S4). We observed a similar U-shaped CSFS in each of three additional African populations (Esan in Nigeria [ESN], Gambian in Western Divisions in the Gambia [GWD], and Mende in Sierra Leone [MSL]) included in the 1000 Genomes Phase 3 data set (Figure S4).

**Figure 1:**
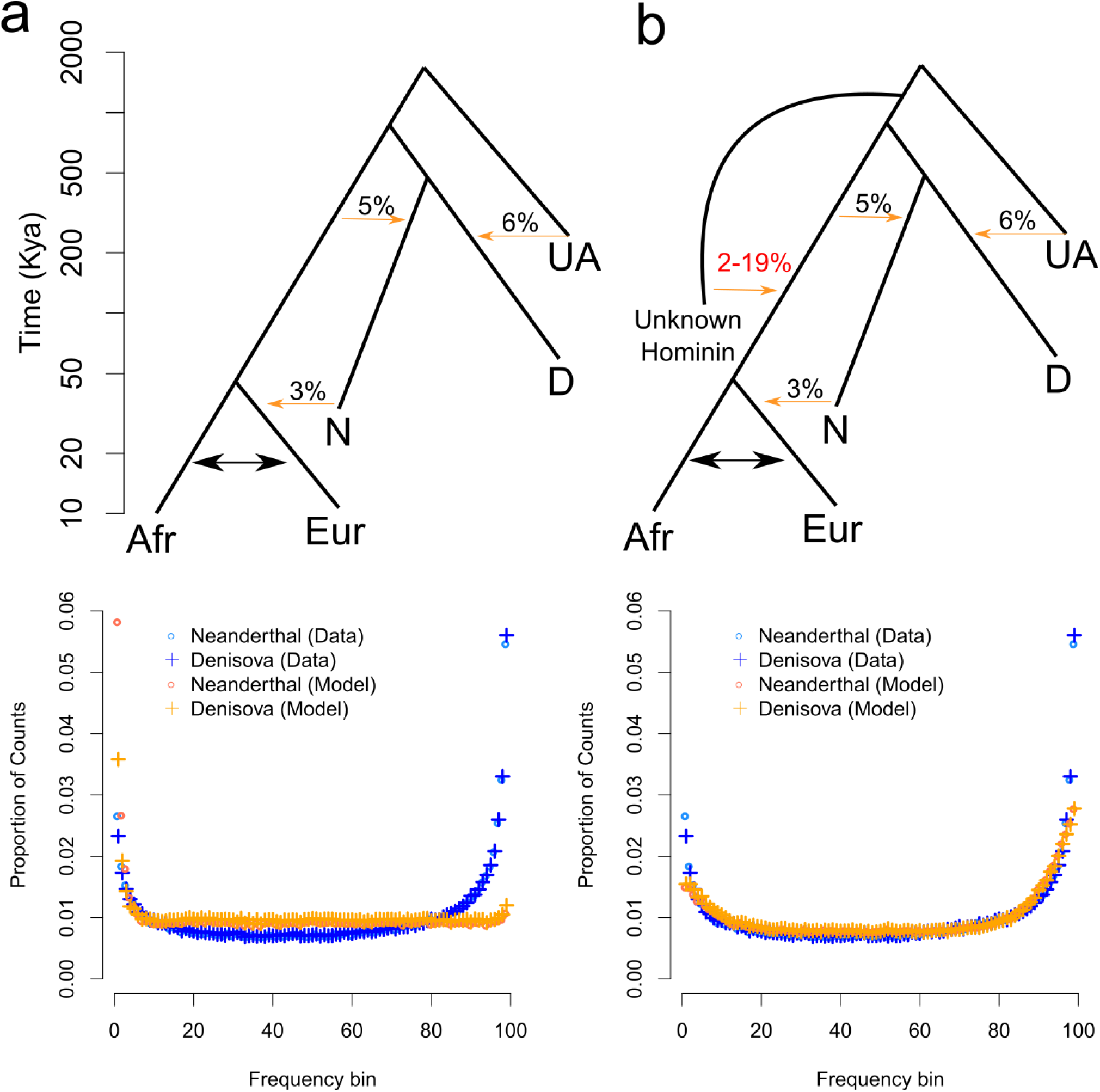
(a) Basic demographic model with conditional site frequency spectrum (CSFS) fit. Key: Afr = Africans, Eur = European, N = Neanderthal, D = Denisovan, UA = Unknown archaic (see (*18*)). Migration between Europe and Africa introduces an excess of low frequency variants, but does not capture the decrease in intermediate frequency variants and increase in high frequency variants. (b) Newly proposed model involving introgression into the modern human ancestor from an unknown hominin that separated from the human ancestor before the split of modern humans and the ancestors of Neanderthals and Denisovans. Below, we show the CSFS fit from the proposed model, which captures the U-shape observed in the data.

Mutational biases as well as errors in determining either the ancestral or the archaic allele could produce the observed CSFS. We confirmed that the shape of the CSFS_YRI,N_ was robust to the inclusion of only transition mutations, only transversion mutations, to the exclusion of hypermutable CpG sites (Figure S7) as well as when we computed the spectrum on the Yoruba genomes separately sequenced in the 1000 Genomes Phase 1 data (Figure S7). Errors in determining the ancestral allele could make low frequency ancestral alleles appear to be high frequency derived alleles and vice-versa and thus could potentially lead to a U-shaped CSFS. However, the shape of the CSFS remains qualitatively unchanged when we used either the chimpanzee genome or the consensus across the orangutan and chimpanzee genomes to determine the ancestral allele (Figure S9). We simulated both ancestral allele misidentification as well as errors in genotype calling in the high-coverage archaic genome. A fit to the data required both a 15% ancestral misidentification rate as well as 3% genotyping error rate in the archaic genome, substantially larger than previous estimates of these error rates (1% for ancestral misidentification rate in the EPO ancestral sequence (*16*) and 0.6% for the modern human contamination in the Vindija Neanderthal (*15*)) (SI Section S2.1; Figure S10). Taken together, these results indicate that the U-shaped CSFS observed in the African populations is not an artifact.

In order to determine whether realistic models of human history can explain the CSFS, we compared the CSFS estimated from coalescent simulations to the observed CSFS_YRI,N_ using a goodness-of-fit test (SI Sections S1.5 and S2). We augmented a model of the demographic history of present-day Africans (*17*) with a model of archaic populations inferred by Pruüfer *et al.* (*15*) (Figures 1, S1, S15). This model includes key interbreeding events between archaic and modern human populations such as the introgression from Neanderthals into non-Africans, from early modern humans into Neanderthals (*6*), as well as introgression into the Denisovans from an unknown archaic population (*18*). This model fails to fit the observed CSFS_YRI,N_ (*p*-value of a Kolmogorov-Smirnoff test on the residuals being normally distributed, KS *p* < 2 ×10^−16^). Extensions of this model to include realistic variation in mutation and recombination rates along the genome (Figure S11; KS *p* < 2 × 10^−16^; SI Section 2) as well as low levels of Neanderthal DNA introduced into African populations via migration between Europeans and Africans do not provide an adequate fit (Figure 1; KS *p* < 2 × 10^−16^; SI Section 2) nor does a model of gene flow between YRI and pygmy populations that has been proposed previously (Figure S12; KS *p* < 2 × 10^−16^; SI Section 2) (*19*). The expectation that the CSFS is uniformly distributed across allele frequencies relies on an assumption of mutation-drift equilibrium in the archaic-modern human ancestor. We confirmed that violations of this assumption (due to bottlenecks, expansions, and population structure in the ancestral population) were also unable to fit the data (KS *p* < 2 ×10^−16^ for all models; SI Section S3; Table S3; Figure S16).

Given that none of the current demographic models are able to fit the observed CSFS, we explored three models where present-day Africans trace part of their ancestry to (A) a population that split from the ancestors of present-day Africans after the split between archaic and modern humans, (B) to an archaic population that split from the ancestor of Neanderthals and Denisovans, or (C) an archaic population that diverged from the ancestor of modern and archaic humans before the archaic-modern human split (Figure S2; SI Section S4). A search for the most likely parameters for models A and B results in the introgressing archaic population splitting off from the modern human population at the same time as the modern human-Neanderthal split. Models A and B fail to fit the observed CSFS even at their most likely parameter estimates (KS *p* = 3.3 ×10^−15^ and *p* = 5.6 × 10^−6^ respectively, SI Section 4) because of insufficient genetic drift in the African population since the split from the archaic hominins (SI Section S5). Model C, on the other hand, is consistent with the data (KS *p* = 0.09) suggesting that part of the ancestry of present-day Africans must derive from a population that diverged prior to the split time of archaic and modern humans. In addition to the goodness-of-fit tests, we examined the likelihood of the best-fit parameters for each of the models and found that model C provides a significantly better fit than other models (model C having a higher composite log likelihood than the next best model Δ ℒ ℒ= ℒ ℒ _Nextbestmodel_ − ℒ ℒ _C_ = 26, 220 when we condition on the Vindija Neanderthal genome and Δ ℒ ℒ = −26, 562 when we condition on the Denisovan genome, Table S4, SI Section S1.4).

We used approximate Bayesian computation to refine the parameters of our most likely model (model C) from the CSFS (SI Section S6). Given the large number of parameters in this model, we fixed parameters that had previously been estimated (*15*) and jointly estimated the split time of the archaic population from the ancestral population, the time of introgression, the fraction of ancestry contributed by the introgressing population as well as the effective population size of the archaic population. We determined that the posterior mean for the split time is 625,000 years B.P. (95% HPD: 360,000-975,000) and the admixture time is 43,000 years B.P. (95% HPD:6,000-124,000), while the posterior mean for the admixture fraction is 0.11 (95% HPD: 0.045-0.19). Analyses of three other African populations (ESN, GWD, and MSL) yielded concordant estimates for these parameters (Figure 2, Table S7). Combining our results across the African populations, we estimate that the archaic population split from the ancestor of Neanderthals and modern humans 360 Ky 1.02 My B.P. and subsequently introgressed into the ancestors of present-day Africans 0 − 124 Ky B.P. contributing 2 − 19% of their ancestry. We caution that the true underlying demographic model is likely to be more complex. To explore aspects of this complexity, we examined the possibility that the archaic lineage diverged at the same time as the split time of modern humans and Neanderthals and found that this model can also produce a U-shaped CSFS with a likelihood that is relatively high, though lower than that of our best fit model (Δℒ ℒ = −9, 819 for the Neanderthal CSFS, Δℒ ℒ = −12, 247 for the Denisovan CSFS, KS *p ≤* 2.9 × 10^−6^). Our estimates of a large effective population size in the introgressing lineage (posterior mean of 25,000, 95% HPD: 23,000-27,000) could indicate additional structure. We find that the *N*_*e*_ of the introgressing lineage in YRI and MSL is larger than in the other African populations, possibly due to a differential contribution from a basal west African branch (*20*).

**Figure 2:**
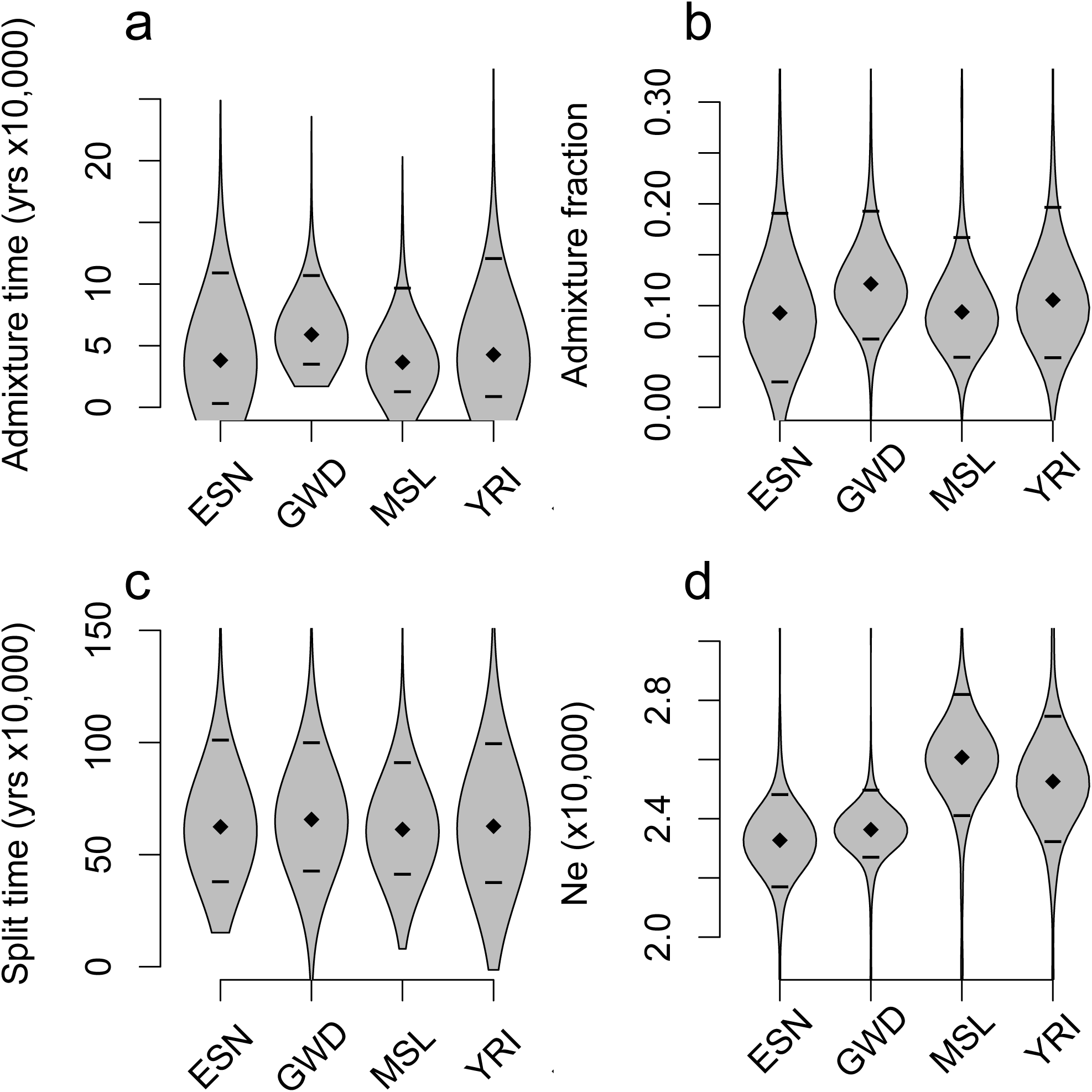
Approximate Bayesian computation estimates for the introgressing population across four African populations (Yoruba from Ibadan (YRI), Esan in Nigeria [ESN], Gambian in Western Divisions in the Gambia [GWD], and Mende in Sierra Leone [MSL]). Posterior means are denoted by diamonds and 95% credible intervals are denoted by lines. (a) the admixture time *t*_*a*_, (b) the admixture fraction *α* (c) the split time of the introgressing population *t*_*s*_, (d) the effective population size of the introgressing population *N*_*e*_. The parameter estimates are largely consistent across the African populations: we estimate split times of 360 Ky −1.02 My B.P., admixture times of 0-124 Ky –B.P., admixture fractions that range from 0.02-0.19, and effective population sizes that range from 22,000-28,000.

While we have chosen to represent the genetic contribution of this unknown population as a single discrete interbreeding event, a more realistic model could include low levels of gene flow in a structured population over an extended period of time. Previously proposed models of ancestral structure do not fit the data (KS *p* < 2 ×10^−16^ for the model described in Sankararaman et al. 2012 and KS *p* < 2 × 10^−16^ for the model proposed in Yang et al 2012, Figure S17), although the model proposed in Yang et al. does produce a slight U-shape. We explored models of population structure in Africa (*21*) where a lineage split from the ancestor of the modern human branch at times ranging from 100 Ky-550 Ky B.P. and continued to exchange genes with the modern human branch until the present. Models of continuous gene flow produce a U-shaped CSFS for low migration rates and deep splits but do not provide an adequate fit (KS *p ≤* 2. × 3 10^−5^, SI Section S7, Figures S13, S14).

To understand the fine-scale distribution of archaic ancestry along the genome, we used a recently developed statistical method (ArchIE) that combines multiple population genetic statistics to identify segments of diverged ancestry in fifty YRI and fifty MSL genomes without the need for an archaic reference genome (*22*) (SI Section S8). Briefly, the method uses summary statistics computed from present-day genome sequences as input to a logistic regression model to estimate the probability that a haploid segment of an individual genome (defined as a contiguous region of length 50 KB) is archaic. While the parameters of the model are estimated by simulating data under a model that closely matches the demographic history relating Neanderthals and non-Africans, we found that ArchIE has 68% power to detect archaic segments at a false discovery rate of about 7% under our best fit demographic model (model C) confirming that its inferences are robust and sensitive to archaic introgression in Africa.

On average, ≃6.6% and ≃ 7.0% of the genome sequences in YRI and MSL were labeled as putatively archaic in ancestry. We sought to test whether the divergent segments identified in YRI and MSL traced their primary ancestry to other African populations (*8, 10, 9*) or to known archaic hominins such as the Neanderthals or Denisovans. We computed the divergence of these segments to a genome sequence from each of four populations: two Central African pygmy populations (Biaka, Mbuti) and two archaic hominin populations (Neanderthal and Denisovan). We expect segments introgressed from any of these populations to be less diverged relative to non-archaic segments. On the contrary, the putatively archaic segments are more diverged, consistent with their source not being any of these populations (Figure 3c; SI Section S8.1). Merging the segments called as archaic across individual genomes, we obtained a total of 482 Megabases and 502 Mb of archaic genome sequence in the YRI and MSL respectively. We estimated the distribution of the time to the most recent common ancestor (TMRCA) between segments labeled archaic and those labeled non-archaic using the pairwise mode of MSMC (*23*) (Figure 3b; SI Section S8.2). and observed that the TMRCA is larger for the putatively archaic class of segments (a 1.69 and 1.23-fold increase in age for YRI and MSL, respectively), consistent with an archaic population being their source.

**Figure 3:**
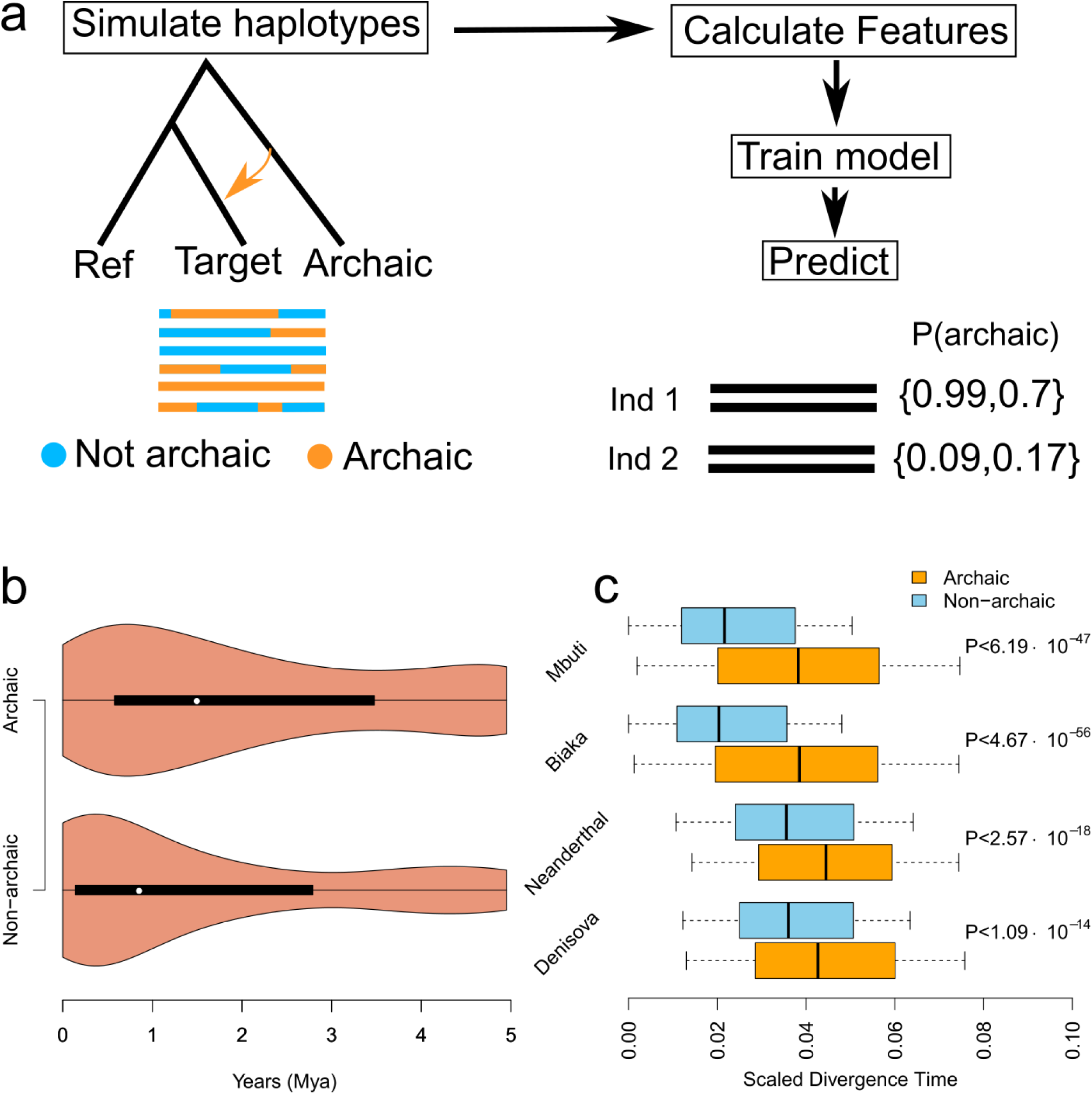
Archaic local ancestry inference performed with ArchIE. (a) Schematic of the method. We simulate data under a model of archaic introgression, calculate population genetic summary statistics, and train a model to predict the probability that a 50KB window in an individual comes from an archaic population. We apply this to data from the Yoruba and Mende populations from Africa. (b) The distribution of segment ages in Yoruba. Segments we denote archaic are 1.69x older than segments we denote non-archaic. (c) Scaled divergence estimates for Yoruba archaic segments versus non-archaic segments from the two modern human pygmy genomes (Mbuti and Biaka) and two archaic genomes (Neanderthal and Denisovan). *P* – values are computed via block jackknife. Archaic segments are more diverged from all four genomes than non-archaic segments.

We examined the frequencies of segments confidently labeled as archaic in ancestry along the genome to investigate whether natural selection could have shaped the distribution of archaic alleles (Figure S40). We found 33 loci with an archaic segment frequency ≥ 50% in the YRI (a cutoff chosen to be larger than the 99.9^*th*^ percentile of introgressed archaic allele frequencies based on a neutral simulations of archaic introgression with parameters related to the time of introgression and admixture fraction chosen conservatively to maximize the drift since introgression; SI Section S8.3; Figure S40) and 37 in the MSL. Some of these genes are at high frequency across both the YRI and MSL, including *NF1*, a tumor suppressor gene (83% in YRI, 85% in MSL), *MTFR2* a gene involved with mitochondrial aerobic respiration in the testis (67% in YRI, 78% in MSL),*HSD17B2*, a gene involved with hormone regulation (74% in YRI, 68% in MSL), *KCNIP4*, which is a gene involved with potassium channels (73% in YRI, 69% in MSL), and *TRPS1*, a gene associated with Trichorhinophalangeal Syndrome (71% in YRI, 75% in MSL; Table 1). Three of these genes have been found in previous scans for positive selection in the YRI: *NF1* (*24, 25*), *KCNIP4* (*26*), and *TRPS1* (*27*). On the other hand, we do not find elevated frequencies at *MUC7*, a gene previously found to harbor signatures of archaic introgression (*28*).

**Table 1:** Genes with a high frequency of archaic segments in Yoruba and Mende. Genes with a frequency above 50% are in bold. These are in the 99.9^*th*^ percentile of introgressed archaic allele frequencies based on neutral simulations.

| Chromosome | Gene name | Frequency (Yoruba) | Frequency (Mende) | Gene type |
| --- | --- | --- | --- | --- |
| chr12 | RP11-125N22.2 | 0.12 | **0.88** | pseudogene |
| chr17 | NF1 | **0.83** | **0.85** | protein coding |
| chr17 | KRT18P61 | **0.84** | 0.36 | pseudogene |
| chr1 | RP11-286M16.1 | **0.84** | **0.81** | lincRNA |
| chr6 | MTFR2 | **0.67** | **0.78** | protein coding |
| chr21 | MIR125B2 | **0.76** | **0.64** | miRNA |
| chr8 | TRPS1 | **0.71** | **0.75** | protein coding |
| chr16 | HSD17B2 | **0.74** | **0.68** | protein coding |
| chr4 | KCNIP4 | **0.73** | **0.69** | protein coding |

We have documented strong evidence for introgression in four present-day sub-Saharan African populations from an archaic lineage that likely diverged prior to the split of modern humans and the ancestors of Neanderthals and Denisovans. The recent time of introgression that we estimate (40, 000 years B.P. with wide credible intervals) suggests that archaic forms persisted in Africa till fairly recently (*29*). Alternately, the archaic population could have introgressed earlier into a modern human population which then subsequently interbred with the ancestors of the populations that we have analyzed here. The models that we have explored here are not mutually exclusive and it is likely that the history of African populations includes genetic contributions from multiple divergent populations as evidenced by the large effective population size associated with the introgressing archaic population. Nevertheless, our results suggest a complex history of interaction between modern and archaic hominins in Africa.

A number of previous studies have found evidence for deeply diverged lineages contributing genetic ancestry to the Pygmy (*9, 8*) and Yoruba (*7, 30*) populations. The signals of introgression across the African populations that we have analyzed raises questions regarding the identity of the archaic hominin and its interactions with modern human populations in Africa. Analysis of the CSFS in the Luhya from Webuye, Kenya (LWK) also reveals signals of archaic introgression although our interpretation is complicated by recent admixture in the LWK that involve populations related to western Africans and eastern African hunter-gatherers (*20*) (SI Section S9). A detailed understanding of archaic introgression as well as its role in adapting to diverse environmental conditions will require analysis of genomes from extant and ancient genomes across the geographic range of Africa.

## Acknowledgments

We thank K Lohmueller, N Patterson, M Lipson, M Schumer, P Moorjani, TV Kent, and members of the Sankararaman and Lohmueller labs for helpful comments and discussions. AD is supported by NSF Graduate Research Fellowship DGE-1650604 and SS is supported in part by NIH grants R00GM111744, R35GM125055, an Alfred P. Sloan Research Fellowship, and a gift from the Okawa Foundation.

