## Supplementary information for "Recovering signals of ghost archaic introgression in African populations"

**S10 ms command lines** **16**

**A The CSFS is uniform under structure in the archaic population** **18**

**B The CSFS is symmetric under Model A** **22**

### S1 The conditional site frequency spectrum (CSFS)

#### S1.1 Definition

We define the conditional site frequency spectrum (CSFS), $\text{CSFS}_{\text{pop}_1,\text{pop}_2}$, as the histogram of the counts of derived alleles in population $\text{pop}_1$ conditional on observing a derived allele in a related outgroup $\text{pop}_2$ [Chen et al., 2007]. We define $c_k$ as the number of SNPs at which the derived allele is present on $k$ chromosomes in a sample of $n$ total chromosomes in $\text{pop}_1$, while a single chromosome in the outgroup $\text{pop}_2$ carries a derived allele. $\text{CSFS}_{\text{pop}_1,\text{pop}_2}$ is the vector of counts $c_k$ for $k \in \{1..n-1\}$.

Chen et al. [2007] showed that if the ancestor of populations $\text{pop}_1$ and $\text{pop}_2$ is at mutation-drift equilibrium (*i.e.* the site frequency spectrum in the ancestor is $f(x) \propto \frac{1}{x}$ where $0 < x < 1$ is the derived allele frequency at a polymorphic SNP) and the two populations $\text{pop}_1$ and $\text{pop}_2$ split with no subsequent admixture, then the $\text{CSFS}_{\text{pop}_1,\text{pop}_2}$ is expected to be uniform, *i.e.*, $\text{CSFS}_{\text{pop}_1,\text{pop}_2}(k) = constant$. Importantly, this result does not depend on any additional aspects of the demographic history of either populations $\text{pop}_1$ and $\text{pop}_2$, except that they are randomly mating. We utilize the CSFS to study introgression in present-day Africans where we set $\text{pop}_1$ to present-day Africans and $\text{pop}_2$ to an archaic population, *i.e.*, Neanderthal or Denisovan.

One of the complications in applying the CSFS to learn about the history of present-day Africans arises from known departures from a simple model of isolation with no subsequent admixture. However, we consider the possibility of structure in the archaic population. This structure could have several forms that include the ancestral Neanderthal population being structured or it could involve gene flow from early modern humans into Neanderthals [Kuhlwilm et al., 2016] or as in the case of Denisovans, this could include gene flow from a highly-diverged archaic population [Prüfer et al., 2014]. We performed extensive simulations to show that structure in the archaic population continues also leads to a uniform CSFS (Section S2). Further, in the Mathematical Appendix A we show that the CSFS is uniform even if there is structure in the archaic population.

However, structure within population the African population ($\text{pop}_1$) since its split from the archaic population ($\text{pop}_2$), *e.g.*, due to admixture, is expected to produce deviations from the uniform CSFS.

#### S1.2 Data processing

For our primary analyses of the CSFS, we used the 1000 Genomes Phase 3 dataset [The 1000 Genomes Project Consortium, 2015] (release 20130502), the high-coverage Vindija Neanderthal genome [Prüfer et al., 2017], and the high-coverage Denisovan genome [Meyer et al., 2012]. We used the annotated ancestral alleles provided by the 1000 Genomes consortium and analyzed only autosomal SNPs. Archaic genotypes (Vindija and Denisovan) come from the pipeline described in Prüfer et al. [2017], which used snpAD for SNP calling (see S3 in [Prüfer et al., 2017]), and required a mapping quality $\geq 25$ and a mappability filter of 100. We did not apply an additional genotype quality filter for the data presented in Figure S4. However, we tested the sensitivity of the spectrum to the choice of genotype quality filters in the archaic when using a GQ filter of $\geq 30$ and $\geq 50$ and see very little difference in the shape of the spectrum (Figure S8).

In addition, we also computed the CSFS using the chimpanzee genome to polarize the ancestral alleles (Figure S9a) [The Chimpanzee Sequencing and Analysis Consortium et al., 2005]. We dropped sites in cases where the chimpanzee allele did not match either human allele. As a further check, we also repeated the analysis restricting only to sites where the chimpanzee and orangutan genomes have matching alleles [Locke et al., 2011]. These results are reported in Figure S9b. Finally, we repeated our analysis filtering out CpG hypermutable sites using the CpG annotations from Prüfer et al. [2014].

#### S1.3 CSFS from the 1000 Genomes data

We computed $\text{CSFS}_{\text{pop}_1,\text{pop}_2}$ where $\text{pop}_1$ is a modern human population and $\text{pop}_2$ is an archaic population. Specifically, we chose $\text{pop}_1$, in turn, to be the Yoruba from Nigeria (YRI), Mende from Sierra Leone (MSL), Esan in Nigeria (ESN), and Gambian from the Western Division in Gambia (GWD) while we chose $\text{pop}_2$ to be either the high-coverage Vindija Neanderthal or the high-coverage Denisovan genome (Figure S4).

We computed the CSFS from the 1000 Genomes phase 3 data [The 1000 Genomes Project Consortium, 2015] for each of the four African populations mentioned above (Figure S4) as well as for the CEPH Europeans from Utah (CEU) and Han Chinese from Beijing (CHB) (Figure S6).

For all populations we observe a U-shaped spectrum with an excess of derived alleles at low and high frequencies. In the African populations, we observe that the CSFS from conditioning on the Denisovan is nearly identical to the Vindija Neanderthal except at the lowest frequency bins, where there is an excess of counts for the Neanderthal CSFS. We interpret this difference as suggestive of low levels of Neanderthal-related ancestry in these populations consistent with previous studies [Prüfer et al., 2014]. In CEU and CHB, we also observe a U-shaped spectrum for both the Vindija Neanderthal and Denisovan, but with a more pronounced difference between the Neanderthal and Denisovan spectra, *i.e.*, an excess of counts in the low frequency derived sites when conditioned on the Vindija Neanderthal relative to the Denisovan. This difference is likely reflective of the Neanderthal introgression event experience by populations outside of Africa around 50,000 years ago [Sankararaman et al., 2012, Fu et al., 2015]. Section S9.2 explores the implication of observing a U-shaped CSFS in African as well as non-African populations.

To determine the robustness of the shape of the CSFS, we recomputed the CSFS in YRI using only transitions, transversions, and after removing CpG sites. We find very similar U-shaped CSFS across these mutation classes (Figure S7).

#### S1.4 Model comparison

We used coalescent simulations to assess whether a demographic model produces a CSFS that matches the empirical CSFS.

To assess the fit of a given demographic model $\mathcal{M}$ to the data, we compare the CSFS computed on the data simulated under $\mathcal{M}$ to that computed on the empirical data. We consider a model in which the empirical CSFS is obtained by sampling from the CSFS computed on the simulated data. For these fits, we model the proportion of SNPs that contain a given number $k$ of derived alleles rather than the number of SNPs. To assess the fit of the simulated CSFS under $\mathcal{M}$ ($\boldsymbol{S}_{\mathcal{M}}$) to the observed CSFS ($\boldsymbol{O}$), we use a multinomial composite likelihood:

$$\mathcal{LL}(\mathcal{M}) = P(\boldsymbol{O}|\boldsymbol{S}_{\mathcal{M}}) = \prod_{k=1}^{n-1} \left( \frac{S_k}{\sum_k S_k} \right)^{O_k}$$

Here $k$ indexes the derived allele count, $S_k$ denotes the number of SNPs with $k$ derived alleles observed in the simulated CSFS, while $O_k$ denotes the number of SNPs with $k$ derived alleles observed in the empirical CSFS. We caution that $\mathcal{LL}$ is a composite likelihood that ignores the dependence among SNPs so that comparisons of $\mathcal{LL}$ must be interpreted with caution.

#### S1.5 Goodness of fit

We defined a goodness of fit statistic that we use to assess whether the CSFS computed under a demographic model explains the major patterns of the empirical CSFS. The goodness of fit statistic is defined from the residuals obtained by trying to fit the simulated CSFS to the empirical CSFS. We assume that the counts of SNPs in each derived allele frequency bin of the empirical CSFS follows a binomial distribution with mean given by the proportion of SNPs that have the same derived allele frequency in the simulated CSFS. One complication is that the counts across bins of derived allele frequencies are not independent due to linkage disequilibrium. To account for this complication, we attempt to estimate the effective number of independent observations in the observed CSFS (rather than assume that each SNP is an independent observation). We define the residual for bin $k$ :

$$r_k = \sqrt{m_{eff}} \frac{o_k - s_k}{\sqrt{s_k(1-s_k)}}$$

Here $m_{eff}$ is the effective number of independent SNPs, $o_k$ represents the proportion of SNPs with derived allele count $k$ in the empirical CSFS, $s_k$ is the proportion of SNPs with derived allele count $k$ in the simulated CSFS, and $k$ indexes the count of derived allele. These residuals are expected to be approximately normally distributed when the number of observations is large (as is the case with the CSFS where each bin has $> 1000$ observations). $m_{eff}$ is a scaling factor to ensure that the residuals are standardized.

To calculate $m_{eff}$, we use two replicate whole genome simulations (3GB) under the same demographic model and set one as the observed data and one as the simulation. We divide the number of bins $k$ by the sum of the squared residuals:

$$m_{eff} = \frac{k}{\sum_{i=1}^{i=k} \left( \frac{o_k - s_k}{\sqrt{s_k(1-s_k)}} \right)^2}$$

A good fit will result in approximately normally distributed residuals, while poor fits will deviate significantly from a normal distribution. To obtain a formal test of fit, we use a Kolmogorov-Smirnoff test comparing the distribution of the residuals to a normal distribution. $p-$values that reject the null hypothesis suggest the model is a poor fit to the data. We use bins of allele counts ranging from 11 to 90, excluding the lowest and highest frequency bins as the counts from these bins are more likely to be affected by unmodeled genotyping errors, leading to false rejections of the null hypothesis. To assess the fit of a class of models (*e.g.* models A, B, and C), we report the $p-$value of the model with parameter estimates obtained via approximate Bayesian computation (Sections S4.1-S4.6).

Finally, we expanded the range of derived allele counts in our goodness of fit computation from $[11, 90]$ to $[6, 95]$ (Table S8). While none of the models fit adequately, model C has substantially higher p-values than the other models indicating that it continues to explain the CSFS better across this range of allele counts. The lack of fit across the expanded range of derived allele counts is likely due to unmodeled complexities in the underlying demographic history as well as error processes that affect the low and high frequency SNPs.

### S2 Current demographic models cannot explain the CSFS

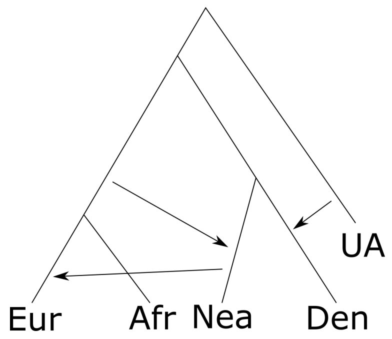

Figure S1: Demographic model from Prüfer et al. [2017]. See Section S2 for details.

We simulated genome sequence data according to the demographic history from Prüfer et al. [2017] (we refer to this model as the 'base model', Figure S1), which contains several major admixture events among modern human populations, Neanderthals, and Denisovans and was fit to explain key population genetic statistics that relate these populations. We use a mutation rate (per basepair) of $1.2 \times 10^{-8}$ and a recombination rate (per basepair) of $1.3 \times 10^{-8}$ and simulate 3,000 1 Mb regions for a total of 3GB of simulated sequence.

The base model includes introgression from an unknown archaic hominin into the Denisovan population (6%, 250Ky B.P.) [Prüfer et al., 2014], introgression from modern humans into Neanderthals (5%, 329Ky

B.P.) [Kuhlwilm et al., 2016] as well as an introgression event from Neanderthals into the ancestors of present-day non-Africans (3%, 50Ky B.P.). Population sizes are set to 10,000 individuals for all lineages except Neanderthal and Denisovan, which have population sizes of 1,000 individuals. The split times for pairs of populations come from Prüfer et al. [2017] and are as follows:

- Unknown archaic - modern human ancestor: 910Ky B.P.
- Archaic-Modern human: 550Ky B.P.
- Neanderthal-Denisovan: 415Ky B.P.
- Africa-Out of Africa: 62.5Ky B.P.

In addition to the above introgression events, we also included introgression from the Neanderthal population into the African lineage 43Ky B.P. which was shown by Prüfer et al. [2017] to lead to improved fits. We used introgression percentages of 1% and 0.015% (Figure S15ab). We see that 1% introgression fraction produces an increase in the counts of low frequency derived alleles in the CSFS and leads to a larger excess in Neanderthals relative to the Denisovan. This result is also seen in the 0.015% case, which is the best fitting parameter as observed by Prüfer et al. [2017]. However, we find that introgression from the ancestors of Neanderthals into the ancestors of Africans is insufficient to produce a U-shaped CSFS (Figure S15ab).

We also performed an additional simulation experiment where we modeled realistic variation in mutation and recombination rates along the genome (while fixing the demographic model to the base model). Mutation rates were calculated by estimating Watterson's $\theta$ [Watterson, 1975] from the number of segregating sites within 1 Mb windows across 50 randomly sampled YRI genomes from the 1000 Genomes Project Phase 3 release and calculating the mutation rate: $\mu = \theta/4N_eL$ where we set $N_e = 10,000$ and $L = 1$ Mb. Recombination rates were estimated from the combined, sex-averaged HapMap recombination map [International HapMap Consortium et al., 2007] (Figure S11).

Next, we simulated data from the model inferred by Gravel et al. [2011]. We restricted our analysis to the African population in this model which we augmented with the archaic populations described above with demographic parameters fixed to those inferred in Prüfer et al. [2017]. The CSFS under this model is also uniform, suggesting the model is insufficient to explain the data (Figure S15c).

In addition, we also explored a model with migration between the Out-of-Africa population and the African population over the last 20Ky B.P. at a rate of 0.0001 migrants per generation as has been suggested by a number of demographic models [Gravel et al., 2011, Harris and Nielsen, 2013, Petr et al., 2019]. We see that such a model does produce a slight increase in the number of derived alleles at a high frequency but is insufficient to explain the signal observed in the data (Figure S15d).

Finally, we simulated the demographic model inferred in Hsieh et al. [2016] relating the Yoruba and the Central African Pygmy (Baka and Biaka) populations. This model has gene flow between the three populations dating back to $\simeq$ 40Ky B.P. Importantly, it is plausible that the gene flow over this period of time could produce the U-shape we see in the data. However, simulations indicate that the parameters inferred from Hsieh et al. [2016] are insufficient to produce the U-shape of the spectrum we observe in the data (KS $p < 2 \times 10^{-16}$) (Figure S12).

#### S2.1 Ancestral misidentification and genotyping error cannot explain the CSFS

One possibility is that CSFS can be explained by a combination of demographic history and errors, in particular errors in identifying the ancestral allele (ancestral misidentification/mispolarization) and genotyping error in the archaic genome that we condition on (Neanderthal or Denisovan). To explore this possibility, we simulated data under the model from Prüfer et al 2017 and added ancestral misidentification ($e_1$) and genotyping error ($e_2$) over a grid of values $e_1, e_2 \in \{0\%, 3\%, 15\%, 30\%\}$. In Figure S10, we begin to observe a U-shaped CSFS with 3% ancestral misidentification and 0% genotyping error (Figure S10e). The best fit CSFS obtained when combining errors with the base model in Section S2 requires 15% ancestral misidentification and 3% genotyping error . In Paten *et al.* 2008, the authors estimate $<$ 1% error in the EPO alignment for the common ancestor of apes and monkeys and that the overall error rate for all ancestors is 3.2%, suggesting that the required ancestral misidentification rate is too large to explain the data [Paten et al., 2008]. In addition, Meyer *et al.* 2012 estimate the sequencing error in the high coverage Denisovan genome as 0.147% per read and a contamination rate of 0.224%, resulting in an overall error rate of 0.371% [Meyer et al., 2012]. In order to erroneously call a genotype at a coverage of 30x (the reported mean coverage for the Denisovan genome), 15 reads or more must support the alternate allele. Therefore, we estimate the genotyping error rate ($e_2$) using the binomial distribution as:

$$e_2 = \sum_{i=15}^{30} \binom{30}{i} 0.00371^i (1 - 0.00371)^{30-i} = 5.1 \times 10^{-29}$$

The above computation of $e_2$ relies on several simplifying assumptions such as independence of errors across reads. This is a reasonable assumption for sequencing errors but is less reasonable for contamination.

For the Vindija Neanderthal, Prüfer [2018] estimates genotyping discordance rates by subsampling the Vindija genome and comparing the frequency of genotype discordance. They subsample read depths $r \in \{1, 2, 3, 4, 5, 6, 7, 8, 9, 10, 12.5, 15, 17.5, 20, 25\}$, where $r$ is the average fold coverage, and obtain a genotype discordance rate for each subsample. We use this data to estimate the probability of genotyping error, $P(g)$ as follows.

$$P(g) = \sum_{i \in r} P(g|i)P(i) = 4.3 \times 10^{-7}$$

In addition to this estimate, Prüfer et al. [2017] report a nuclear contamination genotyping error rate of 0.6%. This represents the probability that the reported genotype comes from a modern human sample rather than the Neanderthal sample. This is an upper bound on the genotyping error rate (the probability that the called archaic genotype differs from the true archaic genotype). These numbers suggest the plausible error rates are lower than what would be required to produce the observed CSFS.

Finally, we note that the best fit model that includes ancestral misidentification and genotyping error is substantially worse than the best fitting model that we explore in Section S4 (model C; $\Delta\mathcal{LL} = -6356$).

### S3 The CSFS cannot be explained by departures from panmixia in the ancestor of archaics and modern humans

One of the assumptions for the CSFS to be uniformly distributed across allele frequencies is the requirement of mutation-drift equilibrium at the archaic-modern human ancestor. In the absence of detailed demographic models of the ancestral population, we explored whether departures from equilibrium in the archaic-modern human ancestor can produce the signals we see in the data using the base model from Prüfer et al. [2017] described in Section S2. We simulated a 5-fold expansion in the ancestor and see that the CSFS slightly increases in the low frequency bins relative to high frequency bins (Figure S16a). We simulated a 5-fold bottleneck in the ancestor and observed the opposite effect: a slight positive slope as the derived allele frequency increases (Figure S16b).

In addition, we turned our attention to models of population structure in the ancestor of archaic and modern humans. A model in which the ancestral population split 840 Ky B.P. into two sub-population followed by subsequent merging into one population 550Ky B.P. produces a uniform spectrum, as does a model with a split at 840Ky B.P. and continuous migration ($5 \times 10^{-5}$ migrants per generation) until 550Ky B.P. (Figure S16cd).

Finally, we explored previous models of ancestral structure from the literature. In particular, we simulated data under models proposed by Yang et al. [2012] and Sankararaman et al. [2012] (Figure S17). Neither model could account for the shape of the CSFS although the model proposed in Yang et al. [2012] does produce a slight U-shape. Overall, departures from mutation-drift equilibrium in the archaic-modern human ancestor are insufficient to produce the signals we see in the data.

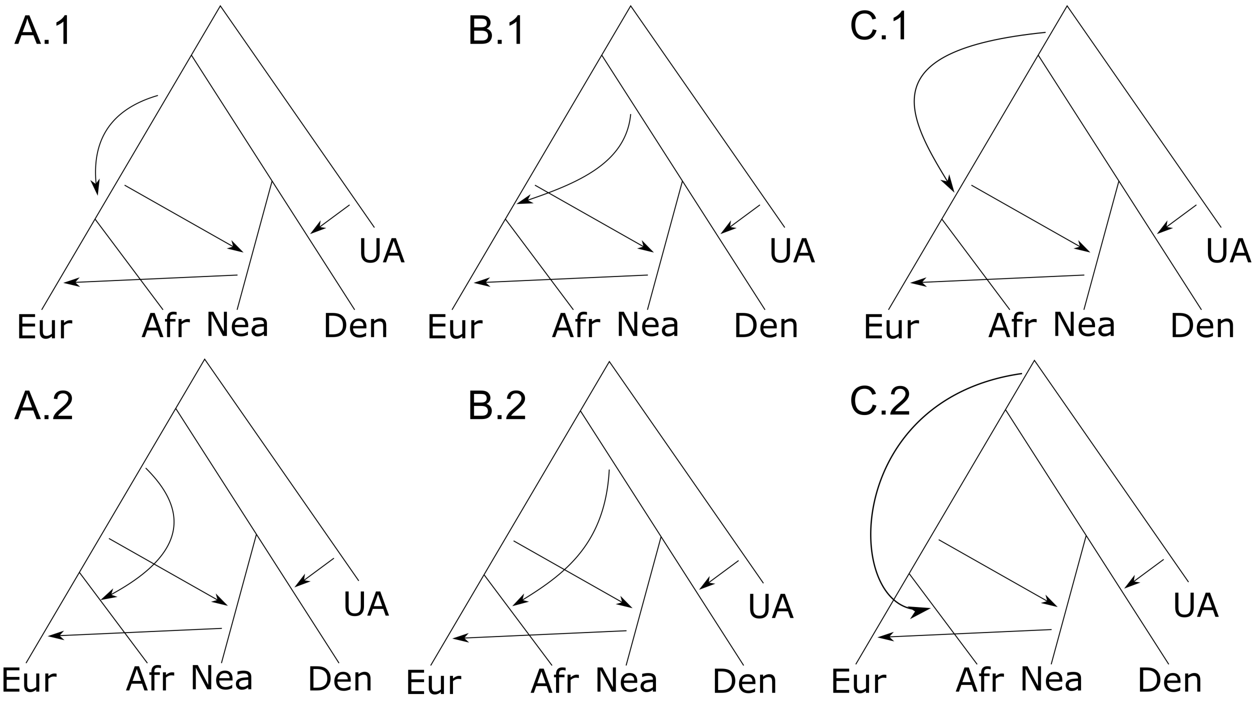

Figure S2: Demographic model topologies for introgression into the ancestors of present-day Africans

### S4 Exploration of models of introgression into the ancestors of present-day Africans

We simulated data assuming a model of introgression into the ancestors of present-day Africans on top of the base model simulated in Section S2. We consider three sources for the gene flow: A) the ancestors of modern humans after their split from the ancestors of archaic humans, B) the ancestors of archaic (Neanderthal/Denisovan) humans after their split from the ancestors of modern humans, C) a population that diverged from the ancestors of archaic and modern humans prior to the archaic-modern human split. For each of these models, we considered two variants depending on the time of introgression: 1. introgression occurred in the modern human ancestor prior to the out-of-Africa (OOA) event, 2. introgression occurred in the African population after the OOA event.

We use ms [Hudson, 2002] to simulate 3,000 unlinked 1 megabase (MB) regions (resulting in 3 gigabases (GB) of sequence). We provide representative ms command lines in Section S10. We used the following parameter values: ancestral effective population size $N_e = 10,000$, mutation rate $\mu = 1.2 \times 10^{-8}$/bp, region length $L = 1,000,000$ base pairs, recombination rate $r = 1.3 \times 10^{-8}$. When scaling generations to years, we use a generation time of 29 years per generation as in Fenner [2005]. Unless otherwise stated, all population sizes are the same as in the baseline model (See Section S2).

We also perform simulations for simple versions of the models presented below where we only simulate the African, Neanderthal, and Denisovan branches (removing the Non-African and unknown archaic branches) without the following gene flow events: from modern humans into the archaic branch, from the unknown archaic into the Denisovan branch, from the Neanderthal into the non-African branch. We present these simulations in Figures S19, S21, S23, S25, S27, and S29. We see that these models are nearly identical to the full models. We also report the fits of the models in Table S5. We describe how these fits are calculated in Sections S1.4 and S1.5.

In addition to the simulations with fixed parameter values described below, we also performed an exploration of the parameter space using approximate Bayesian computation (ABC). We used the proportional CSFS in YRI conditioned on the Vindija Neanderthal as a summary statistic and fit values for the split time of the introgressing lineage ($ts$), the effective population size of the introgressing lineage ($Ne$), the admixture time ($ta$), and the admixture fraction $\alpha$. We drew these parameters from uniform distributions (details of the parameter values are described in the respective sections), simulated 1GB of sequence, and computed the resulting CSFS. We repeated this procedure 10,000 times for each model and obtained posterior distributions using the R package 'abc' [Csilléry et al., 2012]. We used the 'neuralnet' method for non-linear regression (number of networks= 50, number of hidden units= 7) and set the tolerance to 0.1. Then, we simulated 3GB of data from the best fit values and calculated a goodness-of-fit $p$-value (as described in S1.5). We report times in terms of generations. Simulations of the best fitting models are shown in Figure S30.

#### S4.1 Model A.1 – Gene flow from modern human ancestor into the modern human ancestor before OOA

In model A.1, the ancestor of modern humans splits into two populations which later admix. The split occurs at 174Ky B.P. while the admixture occurs at 63.8Ky B.P. (Figure S2). We see that as the admixture fraction increases, there is an increase in allele counts at the low and high frequency bins of the CSFS, which is potentially consistent with the U-shaped spectrum we observe in the data. However, this model is unable to capture the curvature of the spectrum even at large introgression fractions (Figure S18). Further, we show in Mathematical Appendix B that the CSFS under this model is symmetric.

To further explore if specific parameters of model A.1 could explain the CSFS, we performed ABC where we drew parameters relevant to this demographic model from the following distributions:

$$\begin{aligned} ta &\sim Uniform(0.06, 0.19) \\ \alpha &\sim Uniform(0.0, 0.2) \\ ts &\sim Uniform(0.2, 0.47) \\ ne &\sim Uniform(0.1, 3) \end{aligned}$$

Based on the ABC, we found the posterior mean of the admixture time was 2,600 generations B.P. (95% HPD: 2,200-3,300), the admixture fraction was 0.05 (95% HPD: 0.03-0.08), the split time was 17,000 generations B.P. (95% HPD: 15,000-19,100), and the effective population size was 24,000 (95% HPD: 20,000-27,000). Simulations from the best fit values resulted in a KS $p < 2 \times 10^{-16}$.

#### S4.2 Model A.2 – Gene flow from modern human ancestor into African populations after OOA

Model A.2 differs from model A.1 in the time of introgression being more recent, *i.e.*, 50 Ky B.P. (Figure S2). Similar to model A.1, we see an increase in allele counts at the low and high frequency bins. However, this model is still unable to capture the curvature of the spectrum (Figure S20).

For parameter exploration, we drew parameters from the following distributions:

$$\begin{aligned} ta &\sim Uniform(0.01, 0.053) \\ \alpha &\sim Uniform(0.0, 0.2) \\ ts &\sim Uniform(0.1, 0.47) \\ ne &\sim Uniform(0.1, 3) \end{aligned}$$

We found the admixture time was 800 generations B.P. (95% HPD: 500-1200), the admixture fraction was 0.07 (95% HPD: 0.03-0.10), the split time was 17,000 generations B.P. (95% HPD: 14,000-19,000), and the effective population size was 20,000 (95% HPD: 12,500-26,000). Simulations from the best fit values resulted in a KS $p-$value of $3.3 \times 10^{-15}$.

#### S4.3 Model B.1 – Gene flow from archaic branch into the modern human ancestor

In model B.1, we simulate gene flow from the ancestors of the archaic branch containing Neanderthals and Denisovans (Figure S2). This lineage splits off from the ancestors of Neanderthals and Denisovans 464Ky B.P. and introgresses 200Ky B.P. into the modern human ancestor (Figure S22).

For parameter exploration, we drew parameters from the following distributions:

$$\begin{aligned} ta &\sim Uniform(0.06, 0.34) \\ \alpha &\sim Uniform(0.0, 0.2) \\ ts &\sim Uniform(0.35, 0.47) \\ ne &\sim Uniform(0.1, 3) \end{aligned}$$

We found the admixture time was 3,300 generations B.P. (95% HPD: 3,200-3,400), the admixture fraction was .08 (95% HPD: 0.07-0.08), the split time was 18,000 generations B.P. (95% HPD: 18,000-18,100), and the effective population size was 40,100 (95% HPD: 40,000-40,100). Simulations from the best fit values resulted in a KS $p-$value of $9.8 \times 10^{-8}$. The split time effectively reduces this model to a trifurcation.

#### S4.4 Model B.2 – Gene flow from archaic branch into African populations

Model B.2 is identical to model B.1 except that the time of introgression into the African population branch is fairly recent, *i.e.*, 50 kY B.P.(Figure S2, S24).

For parameter exploration, we drew parameters from the following distributions:

$$\begin{aligned} ta &\sim Uniform(0.01, 0.053) \\ \alpha &\sim Uniform(0.0, 0.2) \\ ts &\sim Uniform(0.35, 0.47) \\ ne &\sim Uniform(0.1, 3) \end{aligned}$$

We found the admixture time was 700 generations B.P. (95% HPD: 680-710), the admixture fraction was 0.11 (95% HPD: 0.11-0.11), the split time was 18,800 (95% HPD: 18,700-18,900), and the effective population size was 25,300 (95% HPD: 25,100-25,600). Simulations from the best fit values resulted in a KS $p-$value of $5.6 \times 10^{-6}$. The split time effectively reduces this model to a trifurcation.

#### S4.5 Model C.1 – Gene flow from a new unknown archaic branch into the modern human ancestor

In model C.1, we simulate gene flow from a branch population separates from the modern human and archaic common ancestor prior to their split from each other (696Ky B.P.) and then introgresses into the modern-human ancestor (63.8Ky B.P; Figure S2). We find that this model can produce the U-shaped spectrum we observe in the data, particularly at introgression fractions between 0.05-0.1 (Figure S26).

For parameter exploration, we drew parameters from the following distributions:

$$\begin{aligned} ta &\sim Uniform(0.06, 0.34) \\ \alpha &\sim Uniform(0.0, 0.2) \\ ts &\sim Uniform(0.48, 0.77) \\ ne &\sim Uniform(0.1, 3) \end{aligned}$$

We found the admixture time was 2,700 generations B.P. (95% HPD: 2,200-3,700), the admixture fraction was 0.05 (95% HPD: 0.03-0.07), the split time was 26,000 (95% HPD: 24,700-27,000), and the effective population size was 26,000 (95% HPD: 22,000-29,000). Simulations from the best fit values resulted in a KS $p-$value of 0.09.

#### S4.6 Model C.2 – Gene flow from a new unknown archaic branch into African populations

Model C.2 modifies model C.1 so that introgression into the African population is recent (50Ky B.P; Figure S2). We find that this model can also produce the U-shaped spectrum we observe in the data, again most notably at introgression fractions between 0.05-0.1 (Figure S28).

For parameter exploration, we drew parameters from the following distributions:

$$\begin{aligned} ta &\sim Uniform(0.01, 0.053) \\ \alpha &\sim Uniform(0.0, 0.2) \\ ts &\sim Uniform(0.48, 0.77) \\ ne &\sim Uniform(0.1, 3) \end{aligned}$$

We found the admixture time was 1,360 generations B.P. (95% HPD: 1,300-1,400), the admixture fraction was 0.06 (95% HPD: 0.05-0.06), the split time was 24,300 generations B.P. (95% HPD: 24,000-24,600), and the effective population size was 31,000 (95% HPD: 30,500-31,300). Simulations from the best fit values resulted in a KS $p-$value of $8.2 \times 10^{-6}$.

### S5 Parameter exploration of model A

#### S5.1 Approximate Bayesian computation with ancestral misidentification and genotyping error

We performed additional simulations to explore whether structure more recent than the archaic-modern human split can explain the CSFS.

First, since the CSFS is sensitive to the amount of drift on the introgressing lineage, we simulated model A.2 with several changes in the amount of drift. For all simulations, the introgressing lineage splits 6,000 generations ago, unless otherwise stated. We simulated the following changes to the introgressing lineage:

- $N_e = 100$ (Figure S31; $\Delta\mathcal{LL} = -27,072$)
- $N_e = 1$ (Figure S32; $\Delta\mathcal{LL} = -26,220$)
- $N_e = 1$ with subsequent migration between CEU-YRI (Figure S33; $\Delta\mathcal{LL} = -41,164$)
- $N_e = 0.1$ from a population that splits 200Ky B.P from modern humans (Figure S34; $\Delta\mathcal{LL} = -28,170$)
- a model where the introgressing population branches off the same time as the archaic population and has an $N_e$ of 100 (a trifurcation) (Figure S35; $\Delta\mathcal{LL} = -9,819$)

From these simulations, we find that increasing the amount of drift in the introgressing lineage while retaining a split time of 200Ky B.P. is insufficient to produce a U-shaped CSFS when the introgressing population branches off after the archaic-modern human split. On the other hand, we find that the when the introgressing population is at least as old as the archaic population, we observe a U-shaped CSFS at admixture fractions between 1-20% (Figure S35).

Additionally, we wanted to explore whether any combination of parameters in model A, including genotyping error in the archaic genome or error in the ancestral genome could produce the observed CSFS. We performed two sets of analyses using approximate Bayesian computation. First, we simulated from model A.1, drawing parameters for the split time of the introgressing population (ts – time is in units of $4{*}N_e$ generations where $N_e = 10000$), admixture fraction (alpha), admixture time (ta), effective population size of the introgressing population ($N_e$), ancestral misidentification rate (e1), and genotyping error in the archaic (e2) from the following priors:

$$\begin{aligned}
ta &\sim Uniform(0.06, 0.19)\\
alpha &\sim Uniform(0.0, 0.2)\\
ts &\sim Uniform(0.1, 0.35)\\
ne &\sim Uniform(0.00001, 1)\\
e1 &\sim Uniform(0.00003, 0.3)\\
e2 &\sim Uniform(0.00003, 0.3)
\end{aligned}$$

We constrained the admixture time $ta$ to 2,400-7,600 generations ago and the admixture fraction $alpha$ to 0-20%, which corresponds to a split prior to the Out-of-Africa migration and a low to moderate admixture fraction. We constrained the split time of the introgressing lineage to be at least 4,000 generations ago with a maximum of 14,000 generations ago, corresponding to a period prior to the Neanderthal-modern human split. We constrain the population size of the introgressing lineage to 0.1-10,000 diploid individuals, keeping the size of the population equal to or lower than the ancestral modern human population size. Finally, we allow for a wide interval of 0.003%-30% for each of the errors, encompassing a large range of possibilities including those mentioned in Section S2.1.

We drew parameters 50,000 times, simulated 100 MB and used the R package 'abc' to compute posterior distributions [Csilléry et al., 2012]. We used the 'neuralnet' method to perform non-linear regression setting 'numnet' = 20 and 'sizenet' = 7 and a tolerance of 0.005. We find that the best fitting models have an ancestral misidentification rate of 10.32-14.88% (95% HPD) and an archaic genotyping error rate of 1.95-6.84% (95% HPD; Figure S37).

We applied the same method to model A.2, drawing from the same priors except for the following:

$$\begin{aligned}
ta &\sim Uniform(0.01, 0.053)\\
ts &\sim Uniform(0.2, 0.35)
\end{aligned}$$

We constrained the admixture time to before the Out-of-Africa migration and the split time of the introgressing lineage to at least 8,000 generations ago. For this model, we find that the ancestral misidentification rate is 7.59-13.41% and the archaic genotyping error is 0.63-6.12% (Figure S38).

#### S5.2 Genetic drift between archaics and modern humans constrains model A

It is possible that mis-estimates in the split time of archaics and modern humans could allow model A to fit (that is, a model where the introgressing lineage splits after the archaic and modern human split). In particular, we wondered whether an older split time of archaics and modern humans could fit the data. We assessed this possibility by simulating CSFS from a demographic model that includes a trifurcation between modern humans, archaics, and the introgressing lineage with demographic parameters chosen to maximize drift in the modern human population since its split from the archaic population. Specifically, we chose an archaic-modern human split time of 765Ky B.P. (corresponding to the upper bound on the estimate of the archaic-modern human split time obtained by Prüfer et al. [2014] using a mutation rate of $0.5\times10^{-9}$/bp/year). We converted this estimate from years to generations using a lower bound on the generation time of 25 years per generation. These choices maximize the amount of drift after the archaic-modern human split. We simulated admixture fractions of 1%, 5%, 10%, and 20% (Figure S36). None of these parameters result in a CSFS that yields an adequate fit (KS $p < 1\times10^{-13}$), suggesting that demographic model A does not explain all the features of the CSFS.

Further, we also confirmed that the split time between archaic and modern humans is unlikely to be substantially larger than the values assumed in our simulations by comparing the drift on the modern human lineage produced by these models to estimates of drift in the modern human lineage since the split of modern and archaic humans . We estimated the drift on the modern human lineage since the split from the archaic humans using the property of the CSFS Chen et al. [2007]:

$$\mathrm{CSFS}_{\mathrm{YRI,N}} = \frac{\theta e^{-\tau}}{n+1}$$

Here $\theta = 4N_e\mu L$ (where $N_e$ is the effective population size of the ancestral population, $\mu$ is the mutation rate per basepair, and $L$ is the length of the sequence), $\tau = \int_0^T \frac{dz}{2N_e(z)}$ (where $T$ is the split time of the archaic and modern human populations in generations and $N_e(z)$ denotes the effective population size of the modern human population $z$ generations B.P.), and $n$ is the number of chromosomes in the sample. We wish to solve for $\tau$, the amount of drift. These theoretical results rely on the assumptions that yield a CSFS that is uniform. Since the empirical CSFS ($\mathrm{CSFS}_{\mathrm{YRI,N}}$) is not uniform, we use the mean counts of the CSFS between the allele frequency bins 20%-80%. We take the CSFS computed on 108 YRI individuals from the 1000 Genomes data and project it down to 50 individuals to obtain a mean count of 8,800. Using a mutation rate of $1.2 \times 10^{-8}$ mutations per basepair per generation, a sequence length of $3 \times 10^9$ (approximately the length of the human genome), and an ancestral population size of $10,000$, we obtain an estimate of $\tau \approx 0.48$, which corresponds to $\approx 9600$ generations (assuming an effective population size in the history of present-day African populations of $10,000$). Increasing the mutation rate to $2 \times 10^{-8}$ (an upper bound), we obtain an estimate of $\tau \approx 0.99$, corresponding to $\approx 20,000$ generations. Lowering the sequence length $L$ to more conservative values of $1.5 \times 10^9$ (approximately half the length of the human genome) results in a drift estimate of 0.3 ($\approx 6,000$ generations). Thus, based on the data, we conservatively estimate the amount of drift as $0.3 - 0.99$, or $6,000 - 20,000$ generations.

On the other hand, our simulations assume values of drift of 1.53, or $30,600$ generations. Therefore, conservative simulations in terms of the split time of archaics and modern humans produce much more drift than is supported by the data and still fail to fit the CSFS. We conclude that the introgressing lineage must split before the archaics split from modern humans.

### S6 Estimating parameters for the best fit model of archaic introgression

#### S6.1 Method overview

We used approximate Bayesian computation (ABC) [Csilléry et al., 2012] to estimate parameters. We drew parameters from uniform distributions for the split time, admixture time, effective population size, and admixture fraction and simulated 100 MB of data for each draw (Section S6.2). We repeated this procedure 75,000 times for the model and computed the CSFS for each parameter draw, converting the counts into proportions. We used the R package 'abc' to estimate parameters for each model [Csilléry et al., 2012]. We used the 'neuralnet' algorithm, which uses a non-linear regression method for fitting parameters. We note that because we are using regression that is not bound (i.e. the value is unconstrained) to estimate the parameters, it is possible to get estimates or confidence intervals that are below 0, which are nonsensical. We truncate these values to 0 when they occur. We set the number of networks to 50 and set the size of the hidden nodes to 7. We used a tolerance of 0.02.

We verified that this method is valid in our parameter space of interest by simulating data under model A.2 with a split time of 0.15, admixture time of 0.043 and a 15% admixture fraction (time is in units of $4{*}N_e$ generations where $N_e = 10,000$). We performed ABC on this simulated data and inferred posterior distributions for each parameter. The mean posterior values are as follows (95% highest posterior density [HPD] in parenthesis): admixture time: 0.047 (0.0436-0.501), admixture fraction: 0.1163 (0.0613-0.1739), and split time: 0.152 (0.1137-0.1874). These values are close to the simulated values, suggesting that the ABC provides valid estimates for this problem.

#### S6.2 Parameter estimation

We used ABC to fit 4 parameters from model C: the split time of the introgressing population (ts), the admixture fraction (alpha), the time of introgression (ta), and the effective population size of the introgressing population (ne). We sampled parameters from the following distributions:

$$\begin{aligned} ta &\sim Uniform(0.0, 0.25862069) \\ alpha &\sim Uniform((0.0, 1.0)) \\ ts &\sim Uniform(0.25862069, 1.72413793) \\ ne &\sim Uniform(0.00001, 3) \end{aligned}$$

The times are given in units of 4*$N_e$ generations where $N_e = 10,000$. We applied this method across the 4 African populations in the 1000 Genomes project. Because each population had a different number of samples, we projected the CSFS down to 50 diploid samples using the hypergeometric distribution. In brief, we compute the hypergeometric probability mass function for one variant at a particular frequency class, $i$. Then, we multiply the projected SFS for variant $i$ by the number of variants at that frequency. We perform this over all variant frequencies, $i \in \{1..n-1\}$, where $n$ is the original diploid sample size multiplied by 2.

We chose broad priors over the range of possible values for each of the parameters. Converting generations into years using a value of 29 years per generation, $ta$ ranges approximately from the present day to 300 Ky B.P and $ts$ ranges from 300 Ky B.P to 2 My B.P. $ne$ ranges from 1 diploid individual to 30,000 diploid individuals and $alpha$ includes all values from no contribution (0) to completely replacing the lineage (1).

##### S6.3 Uncertainty in date estimates

We perform our inference measuring time in coalescent units. Converting these units to years is not trivial because of the uncertainty in generation time. To incorporate variation and uncertainty in generation time, we convolve our posterior distributions on generation times with a uniform distribution of generation times between 25 and 33. In particular, we sample from the posterior distribution of admixture time and split time and multiply by $y$, where $y \sim Uniform(25, 33)$. We repeat this procedure 10,000 times and report summaries of this posterior distribution.

#### S7 Continuous migration versus a single pulse

We have modeled the introgression event as a single discrete admixture event in our simulations. It is plausible, however, that this event happened over an extended time scale with continuous migration. We wanted to determine whether a model of continuous migration could qualitatively affect our conclusions, in particular with respect to the split time of the introgressing lineage. In Henn et al. [2018], the authors distinguish between a single origin range expansion with regional persistence model, where a single population expands and receives gene flow from other populations over the past $\leq$ 200Ky B.P. and an African multiregional model where this split occurs further back in time, around 350 Ky B.P.

We simulated a lineage which split from the modern human lineage at various times ranging from 100 Ky B.P to 500Ky B.P with continuous migration with the modern human branch at rates $m \in \{2.5 \times 10^{-5}, 5 \times 10^{-5}, 1 \times 10^{-4}\, 2 \times 10^{-4}, 2 \times 10^{-3}, 2 \times 10^{-2}\}$ migrants per generation. We used a generation time of 29 years to convert these split times to generations. While this model can produce a U-shape with a split as recent as 175Ky B.P (Figure S13), we do not see a good fit to the data (KS $p \leq 2.3 \times 10^{-5}$; Figure S14).

#### S8 Local ancestry inference

We used ArchIE, a method we previously developed for archaic local ancestry inference that does not require access to an archaic reference genome [Durvasula and Sankararaman, 2018]. ArchIE is a machine learning method that uses data simulated under an assumed model and classifies 50KB windows as originating from an archaic population or not. ArchIE uses features that are informative of archaic introgression, including the number of private SNPs in a window, the S*-statistic [Plagnol and Wall, 2006], the minimum distance to an unadmixed reference population, and the distribution of pairwise differences between all segments in a sample.

To estimate the parameters of ArchIE, we simulated training data under a demography where an archaic population splits 12,000 generations before present and introgresses into a modern human population 2,000 generations B.P. with a 2% archaic admixture fraction. We use a fixed mutation rate of $1.25 \times 10^{-8}$ and a recombination rate of $1 \times 10^{-8}$. We simulate two modern human populations, the target, which receives archaic gene flow, and the reference, which does not receive archaic gene flow. The reference population split from the target 2,500 generations B.P. All population sizes were set to 10,000 and we simulated 50 diploid individuals in both the reference and target populations. We simulated 50 KB segments. We have shown through extensive simulations [Durvasula and Sankararaman, 2018] that ArchIE can accurately infer segments of deeply diverged ancestry even in settings where the true demographic model differs from the model used to estimate its parameters.

To further evaluate the accuracy of ArchIE in inferring diverged ancestry, we assessed its ability to recover introgressed lineages simulated our best fitting demographic model C (even though the parameters of ArchIE were estimated under a different demographic model described above). We simulated a lineage that split 625Ky B.P from modern humans and introgressed into the African branch 43Ky B.P with an admixture fraction of 11%. We included introgression from a Neanderthal branch (diverging 12,000 generations ago) into the reference population (representing CEU) 2,000 generations ago at a 0.02 admixture fraction. We simulated a total of 2.5 MB of sequence in 50KB segments.

We report power and false discovery rates (FDR) for both of these scenarios at the 20% false discovery rate threshold obtained under the model used for generating training data ($t_{arch} = P(archaic) = 0.862$). These quantities are defined as:

$$power(t) = \frac{TP(t)}{TP(t) + FN(t)}$$

$$FDR(t) = \frac{FP(t)}{FP(t) + TP(t)}$$

Here $t$ is the probability cutoff (0.862), $TP(t)$ is the number of true positives at threshold $t$(archaic segments that are assigned a probability of being archaic $\geq t$), $FN(t)$ is the number of false negatives at threshold $t$ (archaic segments we do not call archaic), $FP(t)$ is the number of false positives at threshold $t$ (non-archaic segments we call archaic), and $TN(t)$ is the number of true negatives at threshold $t$ (non-archaic segments we call non-archaic). Under our best fitting model, we have 68% power to detect archaic segments and a false discovery rate of 7%.

We applied ArchIE to 50 individuals from the YRI population and 50 from the MSL population using a sample of 50 CEU individuals as the reference population. We applied the 1000 Genomes strict mask and restricted to segments where $\geq$ 90% of the segment is callable. At the threshold $t_{arch}$, ArchIE calls $\simeq$6.6% of the genome in YRI and $\simeq$ 7.0% in the MSL as putatively archaic on average.

Our local ancestry inference provides us with the opportunity to dive deeper into previous analyses of ancestral structure and archaic introgression. In Figure S41 we show the frequency of archaic segments in YRI and MSL across the genome. We observe that the maps are highly correlated (Spearman's $\rho = 0.71$ $p < 2 \times 10^{-16}$). Additionally, in Plagnol and Wall [2006], the authors list 27 loci in the west African Yoruba that could be the result of ancient admixture. We looked at the frequency of archaic segments near these genes (±1MB from the start and end sites). We computed the maximum frequency of archaic segments near these genes and found that the average was 15% in YRI. This is consistent with those genes harboring archaic ancestry and confirms the results of Plagnol and Wall [2006].

#### S8.1 Scaled divergence estimates

We sought to determine whether the segments labeled archaic by ArchIE are likely to trace their ancestry to Central African Pygmy populations (Mbuti and Biaka) or to the Neanderthal and Denisovan genomes. To do this, we comparing the divergence of these segments to genomes from each of these populations. If segments labeled archaic are closer to one of these populations than those labeled non-archaic, it suggests that population may be the source.

We denote the genome we are comparing the calls from ArchIE to as the 'test' genome. To estimate divergence of a segment, we count the number of mutations specific to the archaic segment (*i.e.*, not shared with the test genome) and subtract the number of mutations shared between the archaic segment and the test genome. We divide this number by the number of segregating sites in the window to control for local mutation rate variation.

We test the null hypothesis that the difference in mean divergence between archaic and non-archaic segments is 0. We compute $p-$values for this null using a block jackknife [Kunsch, 1989]. Traversing the genome, we delete 10 MB blocks and recalculate the difference in mean divergence.

We find that for all comparisons, the mean divergence is significantly higher for archaic segments than non-archaic segments.

##### S8.2 Coalescent-based age estimates of introgressed segments

We used the posterior decoding from MSMCv1.1 [Schiffels and Durbin, 2014] to compute the age of putative archaic segments in the genome. When run on a pair of genomes, this method is called PSMC'. Intuitively, this method is a hidden Markov model that uses the observed heterozygosity within an individual to determine the time to most recent common ancestor in blocks along the genome. We selected one individual from each population (HG03079 for MSL and NA18486 for YRI) and computed the posterior distribution of coalescent times in 1000bp bins. The times from MSMC are scaled by 2N, where N is determined by the heterozygosity. We divide the observed $\hat{\theta}$ (Watterson's $\theta$, computed by MSMC. See methods section in Schiffels and Durbin [2014]) by the mutation rate $\mu = 1.25 \times 10^{-8}$. We multiply the coalescent times outputted by MSMC by this factor in order to scale it to generations. Then, we multiplied the generations by 29 years per generation to obtain coalescence estimates in years [Fenner, 2005]. We then intersected these times with the segment calls from ArchIE, restricting the analysis to confidently called archaic segments ( probability of being archaic $P(archaic) \geq t_{arch}$) and confidently called non-archaic segments ($P(archaic) \leq 1 - t_{arch}$). We restricted the analysis to windows where $> 90\%$ of the bases are callable as defined by the 1000 Genomes strict mask.

We find that in YRI, the median coalescent time for archaic segments is 1.5 Mya and 0.89 Mya for non-archaic segments, a 1.69-fold increase in age. In MSL, we find the median coalescent time is 1.1 Mya and 0.93 Mya for archaic and non-archaic segments respectively.

##### S8.3 Adaptive archaic introgression

We examined the maps of archaic introgression in YRI and MSL, computing the frequency of the archaic segments within 50 KB windows. We intersected our calls with gene annotations from Gencode v19 using bedtools [Quinlan and Hall, 2010] and included a gene if any basepair in a single 50KB segment overlapped with the gene [Frankish et al., 2019].

In order to evaluate whether the high frequency archaic segments could have risen to their present-day frequencies as a result of drift, we performed coalescent simulations using the parameters we estimated in S6.2. We simulated 100 replicates of 50KB with a 13% introgression pulse 1400 generations ago from a lineage that split from modern humans 23,000 generations ago. The introgressing lineage had an effective population size of 25,000. We chose the introgression pulse and introgression time to be on the upper bound of the 95% HPD estimates so as to maximize the drift in the population after introgression. Thus, we expect the distribution of archaic allele frequencies to be an upper bound on the neutral distribution. The distribution of allele frequencies for alleles originating in the introgressing lineage (Figure S40) gives us the null distribution of archaic allele frequencies expected under neutrality. Observing archaic alleles with frequencies above this threshold suggests that these alleles have risen to high frequencies due to selection.

#### S9 Extended Discussion

##### S9.1 Results from the Luhya

Examination of the CSFS in the Luhya from Kenya (LWK) also reveals a U-shaped spectrum, similar to the other African populations examined here (Figure S5). We performed the same ABC-based parameter estimation procedure and found that a linage splitting 624,000 years ago (95% HPD: 290,100-1,137,000) with an $N_e$ of 24,000 (95% HPD: 22,000-24,000) and introgressing 50,000 years ago (95% HPD: 11,000-113,00) is able to explain the shape of the spectrum. These parameters are similar to the other African populations.

However, we find that an introgression fraction of 0.18 (95 % HPD: 0.09-0.31) is also required, which is much higher than the other populations. Inferences in the LWK could be complicated by non-African ancestry in the Luhya, resulting in increased estimates of the admixture fraction[Wall et al., 2013]. However, gene flow from the Neanderthals mediated through non-African ancestry into the LWK may not be enough to quantitatively explain this increase. Another possibility is other unmodeled gene flow events outside of the ones we describe here.

##### S9.2 Pre- versus post- Out-of-Africa

We inspected the CSFS in the CEU conditioned on the Denisovan allele, since there is little Denisovan-related ancestry in Europeans [Mallick et al., 2016]. If the introgression event is shared in YRI and CEU, we expect to see a U-shaped CSFS in both populations (Figure S39abc). However, if the event is not shared and is specific to African populations only, the signal is expected to only appear in the African CSFS (Figure S39def). We observe that the CEU CSFS, like the CSFS in the African populations, is U-shaped with an increase in the counts of high frequency alleles. (Figures S4 and S6). Further, we also observe a U-shape in the CSFS computed in the Han Chinese (CHB) population (Figure S6). These results suggest that a component of the archaic ancestry that we detect in African populations is shared with non-African populations. We view these results as provisional and additional detailed modeling needs to be done to resolve the history and timing of archaic introgression.

#### S10 ms command lines

Here, we provide the command lines used in the various models.

**Model A.1**

```
ms 202 1 -t 480 -r 519.99948 1000000 -I 4 100 100 1 1 -en 0 3 0.1 -en 0 4 0.1
  -es 0.04310345 1 0.97 -ej 0.05387931 5 3 -ej 0.05387931 2 1 -es 0.055 1 ${admix}
  -ej 0.15 6 1 -es 0.2155172 4 0.94 -en 0.2155172 7 0.2 -es 0.2836207 3 0.95
  -ej 0.2836207 8 1 -ej 0.3577586 4 3 -ej 0.4741379 3 1 -ej 0.7844828 7 1
```

**Model A.2**

```
ms 202 1 -t 480 -r 519.99948 1000000 -I 4 100 100 1 1 -en 0 3 0.1 -en 0 4 0.1
  -es 0.04310345 1 0.97 -es 0.04310345 2 ${admix} -ej 0.05387931 5 3 -ej 0.05387931 2 1
  -ej 0.15 6 1 -es 0.2155172 4 0.94 -en 0.2155172 7 0.2 -es 0.2836207 3 0.95 -ej 0.2836207 8 1
  -ej 0.3577586 4 3 -ej 0.4741379 3 1 -ej 0.7844828 7 1
```

**Model B.1**

```
ms 202 1 -t 480 -r 519.99948 1000000 -I 4 100 100 1 1 -en 0 3 0.1 -en 0 4 0.1
  -es 0.04310345 1 0.97 -ej 0.05387931 5 3 -ej 0.05387931 2 1 -es 0.1725 1 ${admix}
  -es 0.2155172 4 0.94 -en 0.2155172 7 0.2 -es 0.2836207 3 0.95 -ej 0.2836207 8 1
  -ej 0.3577586 4 3 -ej 0.40 6 3 -ej 0.4741379 3 1 -ej 0.7844828 7 1
```

**Model B.2**

```
ms 202 1 -t 480 -r 519.99948 1000000 -I 4 100 100 1 1 -en 0 3 0.1 -en 0 4 0.1
  -es 0.04310345 1 0.97 -es 0.04310345 2 ${admix} -ej 0.05387931 5 3 -ej 0.05387931 2 1
  -es 0.2155172 4 0.94 -en 0.2155172 7 0.2 -es 0.2836207 3 0.95 -ej 0.2836207 8 1
  -ej 0.3577586 4 3 -ej 0.4 6 3 -ej 0.4741379 3 1 -ej 0.7844828 7 1
```

**Model C.1**

```
ms 202 1 -t 480 -r 519.99948 1000000 -I 4 100 100 1 1 -en 0 3 0.1 -en 0 4 0.1
  -es 0.04310345 1 0.97 -ej 0.05387931 5 3 -ej 0.05387931 2 1
  -es 0.055 1 ${admix} -es 0.2155172 4 0.94 -en 0.2155172 7 0.2
  -es 0.2836207 3 0.95 -ej 0.2836207 8 1 -ej 0.3577586 4 3
  -ej 0.4741379 3 1 -ej 0.6 6 1  -ej 0.7844828 7 1
```

**Model C.2**

```
ms 202 1 -t 480 -r 519.99948 1000000 -I 4 100 100 1 1 -en 0 3 0.1 -en 0 4 0.1
  -es 0.04310345 1 0.97 -es 0.04310345 2 ${admix} -ej 0.05387931 5 3 -ej 0.05387931 2 1
  -es 0.2155172 4 0.94 -en 0.2155172 7 0.2 -es 0.2836207 3 0.95 -ej 0.2836207 8 1
  -ej 0.3577586 4 3 -ej 0.4741379 3 1 -ej 0.6 6 1  -ej 0.7844828 7 1
```

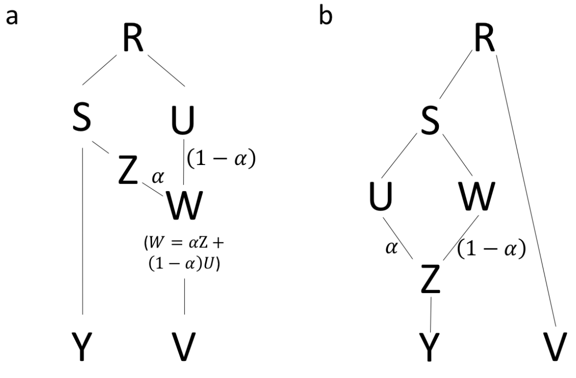

Figure S3: Demographic model topologies for mathematical results regarding a) the expectation of the CSFS when there is structure in the archaic population (Section A) and b) the symmetry of the CSFS under model A (Section B)

### A The CSFS is uniform under structure in the archaic population

We consider a simplified demographic model where the ancestors of the African and Neanderthal populations split with no subsequent gene flow. However, we consider the possibility of structure in the archaic population. This structure could have several forms that include the ancestral Neanderthal population being structured or it could involve gene flow from early modern humans into Neanderthals [Kuhlwilm et al., 2016] or as in the case of Denisovans, this could include gene flow from a highly-diverged archaic population [Prüfer et al., 2014].

To consider these possibilities, we consider the ancestral population ($R$) that splits into two populations ($S$ and $U$). At a later time, $S$ further splits into two populations: one of which evolves to become present-day Africans $Y$ and the other $Z$ admixes with $U$ to give rise to population $W$ (with admixture fraction $\alpha : 1-\alpha$) which then further evolves into the Neanderthal population $V$. See Figure S3a for a schematic of the topology. We use $R$ to represent the random variable denoting the frequency of the derived allele in population $R$ and $r$ to denote the realized value of this random variable (and analogously for each of the other populations). We measure the time elapsed $\tau$ from each of these populations to its parent population (where defined) in diffusion units, *i.e.*, $\tau = \frac{t}{2N}$ where $t$ is the elapsed time in generations and $N$ is the diploid effective population size that is assumed constant. When the population size $N(x)$ changes with time, we define $\tau = \int_0^t \frac{dx}{2N(x)}$. The drift leading to population $S$ from its parent population $R$ is denote $\tau_S$ and so on. Within each of the populations, we assume random mating. Finally, we also assume that populations $R$ and $S$ are at mutation-drift equilibrium.

We assume no recurrent mutation and neutral evolution of alleles. Let $X(\tau)$ denote the frequency of an allele at time $\tau$ where is time scaled in diffusion units. The SNP is assumed polymorphic at time $\tau = 0$, $0 < X(0) < 1$. The evolution of $X(\tau)$ can be described by a Wright-Fisher diffusion that describes the effects of genetic drift on the allele frequency. Let $P(y|x,\tau)$ denote the transition probability that the allele frequency is $y$ at time $\tau$ given that allele frequency at time 0 is $x$ (we will overload $P$ to refer to the probability density when $0 < y < 1$ and the probability when $y = 0$ or $y = 1$).

We have:

$$\begin{aligned} P(y|x,\tau) &= P(y, \mathbf{1}(y=0)|x,\tau) + P(y, \mathbf{1}(0<y<1)|x,\tau) + P(y, \mathbf{1}(y=1)|x,\tau) \\ &= \psi_0(x;\tau) + \phi(y,x;\tau) + \psi_1(x;\tau) \end{aligned} \quad (1)$$

The explicit forms of $\psi_0$, $\psi_1$, and $\phi$ have been solved by Kimura [1955].

We sample $n$ chromosomes in Africans and 1 chromosome from the Neanderthal. The CSFS $f_i$ is the number of SNPs at which $i$ chromosomes ($i \in \{1, \ldots, n-1\}$) in the African population carry the derived allele and the chromosome sampled from Neanderthal also carries the derived allele.

We show that:

$$f_i = constant \tag{2}$$

We have:

$$f_i = \binom{n}{i} \int y^i (1-y)^{n-i} f(y) dy \tag{3}$$

Here $f(y)$ denotes the population conditional frequency spectrum in Africans conditioned on observing a derived allele in Neanderthal. We note that the only SNPs that will contribute to $f_i$ are those that are polymorphic at $Y$ and contain the derived allele at $V$. This requires that these SNPs are also polymorphic at $S$. Further, it will be useful to partition the SNPs contributing to $f(y)$ to those that were polymorphic at $R$ (and also polymorphic at $S$) ($f_{old}$) and those mutations that arose in the ancestral population leading to $S$ from $R$ ($f_{new}$).

$$f(y) = f_{old}(y) + f_{new}(y) \tag{4}$$

The assumption of mutation drift equilbrium at $R$ and $S$ implies:

$$f_0(s) = f_{0,new}(s) + \int \phi(s, r; \tau_S) f_0(r) dr \tag{5}$$

Here $f_0$ is the frequency spectra at $R$ and $S$ respectively while $f_{0,new}$ denotes the frequency spectrum due to mutations that arose after $R$.

Let $P(r,s,u,z,w,y,v)$ denote the joint probability $P(R=r,S=s,U=u,Z=z,W=w,Y=y,V=v)$.

$$
\begin{aligned}
f_{old}(y) &= \int P(r,s,u,z,w,y,v)v drdsdudzdwdv \\
&= \int \Big\{ P(r)P(s|r,\tau_S)P(u|r,\tau_U)P(z|s,\tau_Z) \\
&\qquad P(v|w,\tau_V)P(y|s,\tau_Y)\mathbf{1}\left(w=\alpha z+(1-\alpha)u\right)v \Big\} drdsdudzdwdv \qquad (6) \\
&= \int \Big\{ f_0(r)\phi(s,r;\tau_S)P(u|r,\tau_U)P(z|s,\tau_Z) \\
&\qquad P(v|w,\tau_V)\phi(y,s;\tau_Y)\mathbf{1}\left(w=\alpha z+(1-\alpha)u\right)v \Big\} drdsdudzdwdv \qquad (7) \\
&= \int \Big\{ f_0(r)\phi(s,r;\tau_S)P(u|r,\tau_U)P(z|s,\tau_Z) \\
&\qquad \left(\int P(v|w,\tau_V)vdv\right)\phi(y,s;\tau_Y)\mathbf{1}\left(w=\alpha s+(1-\alpha)u\right) \Big\} drdsdudzdw \\
&= \int \Big\{ f_0(r)\phi(s,r;\tau_S)P(u|r,\tau_U)P(z|s,\tau_Z) \\
&\qquad \phi(y,s;\tau_Y)\left(\int \mathbf{1}\left(w=\alpha z+(1-\alpha)u\right)wdw\right) \Big\} drdsdudz \qquad (8) \\
&= \int \Big\{ f_0(r)\phi(s,r;\tau_S)P(u|r,\tau_U)P(z|s,\tau_Z)\phi(y,s;\tau_Y)\left(\alpha z+(1-\alpha)u\right) \Big\} drdsdudz \qquad (9) \\
&= \alpha \int \Big\{ f_0(r)\phi(s,r;\tau_S)\left(\int P(u|r,\tau_U)du\right)\left(\int P(z|s,\tau_Z)zdz\right)\phi(y,s;\tau_Y) \Big\} drds \\
&\quad +(1-\alpha)\int \Big\{ f_0(r)\phi(s,r;\tau_S)\left(\int P(u|r,\tau_U)udu\right)\left(\int P(z|s,\tau_Z)dz\right)\phi(y,s;\tau_Y) \Big\} drds \qquad (10) \\
&= \alpha \int \Big\{ f_0(r)\phi(s,r;\tau_S)s\phi(y,s;\tau_Y) \Big\} drds + (1-\alpha)\int \Big\{ f_0(r)\phi(s,r;\tau_S)r\phi(y,s;\tau_Y) \Big\} drds \qquad (11) \\
&= \alpha \int \Big\{ \left(\int f_0(r)\phi(s,r;\tau_S)dr\right) s\phi(y,s;\tau_Y) \Big\} ds + (1-\alpha)\int \Big\{ \left(\int f_0(r)\phi(s,r;\tau_S)rdr\right)\phi(y,s;\tau_Y) \Big\} ds \\
&= \alpha \int \left(\int f_0(r)\phi(s,r;\tau_S)dr\right) s\phi(y,s;\tau_Y)ds + (1-\alpha)\theta \int \left(\int \phi(s,r;\tau_S)dr\right)\phi(y,s;\tau_Y)ds \qquad (12) \\
&= \alpha \int \left(\int f_0(r)\phi(s,r;\tau_S)dr\right) s\phi(y,s;\tau_Y)ds + (1-\alpha)\theta\exp\left(-\tau_S\right)\int \phi(y,s;\tau_Y)ds \qquad (13) \\
&= \alpha \int \left(\int f_0(r)\phi(s,r;\tau_S)dr\right) s\phi(y,s;\tau_Y)ds + (1-\alpha)\theta\exp\left(-\tau_S\right)\exp\left(-\tau_Y\right) \qquad (14)
\end{aligned}
$$

Here Equation 6 follows from the conditional independence statements implied by our model. Equation 7 follows because only the polymorphic SNPs at $S$ and $Y$ contribute to $f(y)$. Equations 8 and 11 follow from the fact that the Wright-Fisher diffusion is a Martingale so that $\mathbb{E}\left[X(\tau)|X(0)\right]=X(0)$. Equation 12 follows from the fact that the frequency spectrum of derived alleles at $R$ under mutation-drift equilibrium is given by $f_0(r)=\frac{\theta}{r}$ [Griffiths, 2003]. Equations 13 and 14 follow from explicit integration using the analytical solution for $\phi(y,x;\tau)$ [Kimura, 1955, Chen et al., 2007].

$$
\begin{aligned}
f_{new}(y) &= \int P(s,z,w,y,v)v ds dz dw dv \\
&= \int \Big\{ P(s)P(z|s,\tau_Z)P(v|w,\tau_V)P(y|s,\tau_Y)\mathbf{1}\left(w=\alpha z\right)v \Big\} ds dz dw dv \qquad (15) \\
&= \int \Big\{ f_{0,new}(s)P(z|s,\tau_Z)P(v|w,\tau_V)\phi(y,s;\tau_Y)\mathbf{1}\left(w=\alpha z\right)v \Big\} ds dz dw dv \qquad (16) \\
&= \int \Big\{ f_{0,new}(s)P(z|s,\tau_Z)\left(\int P(v|w,\tau_V)v dv\right)\phi(y,s;\tau_Y)\mathbf{1}\left(w=\alpha z\right) \Big\} ds dz dw \\
&= \int \Big\{ f_{0,new}(s)P(z|s,\tau_Z)\phi(y,s;\tau_Y)\left(\int \mathbf{1}\left(w=\alpha z\right) w dw\right) \Big\} ds dz \qquad (17) \\
&= \alpha \int \Big\{ f_{0,new}(s)\left(\int P(z|s,\tau_Z) z dz\right)\phi(y,s;\tau_Y) \Big\} ds \qquad (18) \\
&= \alpha \int f_{0,new}(s) s \phi(y,s;\tau_Y) ds \qquad (19)
\end{aligned}
$$

Here Equation 15 follows from the conditional independence statements implied by our model. Equation 16 follows because only the polymorphic SNPs at $Y$ contribute to $f(y)$. Equations 17 and 19 follow from the Martingale property of Wright-Fisher diffusion.

From Equations 14 and 19:

$$
\begin{aligned}
f(y) &= f_{old}(y) + f_{new}(y) \\
&= \alpha \int \left( \int f_0(r)\phi(s,r;\tau_S)dr + f_{0,new}(s) \right) s\phi(y,s;\tau_Y)ds + (1-\alpha)\theta \exp\left(-(\tau_S+\tau_Y)\right) \\
&= \alpha \int f_0(s) s\phi(y,s;\tau_Y)ds + (1-\alpha)\theta \exp\left(-(\tau_S+\tau_Y)\right) \qquad (20) \\
&= \alpha\theta \int \phi(y,s;\tau_Y)ds + (1-\alpha)\theta \exp\left(-(\tau_S+\tau_Y)\right) \qquad (21) \\
&= \theta \Big( \alpha \exp\left(-\tau_Y\right) + (1-\alpha)\exp\left(-(\tau_S+\tau_Y)\right) \Big) \qquad (22)
\end{aligned}
$$

Equation 20 follows from Equation 5 while Equation 21 follows from the mutation-drift equilibrium and Equation 22 follows from previous results [Chen et al., 2007].

Substituting Equation 22 into Equation 3, we have:

$$
f_i = \frac{1}{n+1}\theta \Big( \alpha \exp\left(-\tau_Y\right) + (1-\alpha)\exp\left(-(\tau_S+\tau_Y)\right) \Big) \qquad (23)
$$

### B The CSFS is symmetric under Model A

We consider a simplified version of demographic model A where the ancestors of the African and Neanderthal populations split with no subsequent gene flow. However, in this model, the ancestral African population then splits into two sub-populations that later admix.

The ancestral population (labeled $R$) splits into the two populations that are the ancestors of Africans and Neanderthals (labeled $V$) respectively. The ancestral African population (labeled $S$) splits into two sub-populations. These two sub-populations (labeled $U$ and $W$) admix in the ratio $\alpha : 1-\alpha$ to form population $Z$ which then continues to evolve to give rise to the present-day African population (labeled $Y$). We sample a single chromosome from the Neanderthal population (labeled $V$). See Figure S3b for a schematic of the topology. We use $R$ to represent the random variable denoting the frequency of the derived allele in population $R$ and $r$ to denote the realized value of this random variable (and analogously for each of the other populations). We measure the time elapsed $\tau$ from each of these populations to its parent population (where defined) in diffusion units, *i.e.*, $\tau = \frac{t}{2N}$ where $t$ is the elapsed time in generations and $N$ is the diploid effective population size that is assumed constant. When the population size $N(x)$ changes with time, we define $\tau = \int_0^t \frac{dx}{2N(x)}$. The drift leading to population $S$ from its parent population $R$ is denote $\tau_S$ and so on. Within each of the populations, we assume random mating. Finally, we also assume that population $R$ is at mutation-drift equilibrium.

As in the previous section, we assume no recurrent mutation and neutral evolution of alleles. Let $X(\tau)$ denote the frequency of an allele at time $\tau$ where is time scaled in diffusion units. The evolution of $X(\tau)$ can be described by a Wright-Fisher diffusion that describes the effects of genetic drift on the allele frequency and $P(y|x,\tau)$ denotes the transition probability that the allele frequency is $y$ at time $\tau$ given that allele frequency at time 0 is $x$.

We sample $n$ chromosomes in Africans and 1 chromosome from the Neanderthal. The CSFS $f_i$ is the number of SNPs at which $i$ chromosomes ($i \in \{1,\ldots,n-1\}$) in the African population carry the derived allele and the chromosome sampled from Neanderthal also carries the derived allele. We show that the CSFS is symmetric under this model: $f_i = f_{n-i}, i \in \{1,\ldots,n-1\}$.

We then have:

$$f_i = \binom{n}{i} \int y^i (1-y)^{n-i} f(y) dy \quad (24)$$

$f(y)$ denotes the conditional frequency spectrum in Africans conditioned on observing a derived allele in Neanderthal. We note that the only SNPs that will contribute to $f_i$ are those that are polymorphic at $Y$ and contain the derived allele at $V$. This requires that these SNPs are also polymorphic at $R$, $S$, and $Z$.

Let $P(r,s,u,w,z,y,v)$ denote the joint probability $P(R=r, S=s, U=u, W=w, Z=z, Y=y, V=v)$.

$$\begin{aligned}
f(y) &= \int P(r,s,u,w,z,y,v) v drdsdudwdzdv \\
&= \int \left\{ P(r)P(v|r,\tau_V)P(s|r,\tau_S)P(u|s,\tau_U)P(w|s,\tau_W)P(y|z,\tau_Y)\mathbf{1}\left(z=\alpha u+(1-\alpha)w\right) \right\} vdrdsdudwdzdv \quad (25) \\
&= \int \left\{ f_0(r)P(v|r,\tau_V)\phi(s,r;\tau_S)P(u|s,\tau_U)P(w|s,\tau_W)\phi(y,z;\tau_Y)\mathbf{1}\left(z=\alpha u+(1-\alpha)w\right) \right\} vdrdsdudwdzdv \\
&= \int \left\{ f_0(r)\left(\int P(v|r,\tau_V)vdv\right)\phi(s,r;\tau_S)P(u|s,\tau_U)P(w|s,\tau_W)\phi(y,z;\tau_Y)\mathbf{1}\left(z=\alpha u+(1-\alpha)w\right) \right\} drdsdudwdz \\
&= \int \left\{ rf_0(r)\phi(s,r;\tau_S)P(u|s,\tau_U)P(w|s,\tau_W)\phi(y,z;\tau_Y)\mathbf{1}\left(z=\alpha u+(1-\alpha)w\right) \right\} drdsdudwdz \quad (26) \\
&= \theta \int \left\{ \phi(s,r,\tau_S)P(u|s,\tau_U)P(w|s,\tau_W)\phi(y,z;\tau_Y)\mathbf{1}\left(z=\alpha u+(1-\alpha)w\right) \right\} drdsdudwdz \quad (27) \\
&= \theta \int \left\{ \left(\int \phi(s,r,\tau_S)dr\right) P(u|s,\tau_U)P(w|s,\tau_W)\phi(y,z;\tau_Y)\mathbf{1}\left(z=\alpha u+(1-\alpha)w\right) \right\} dsdudwdz \quad (28) \\
&= \theta \exp\left(-\tau_S\right) \int \left\{ P(u|s,\tau_U)P(w|s,\tau_W)\phi(y,z;\tau_Y)\mathbf{1}\left(z=\alpha u+(1-\alpha)w\right) \right\} dsdudwdz \quad (29)
\end{aligned}$$

Here Equation 25 follows from the conditional independence statements implied by our model. Equation 26 follows from the fact that the Wright-Fisher diffusion is a Martingale. Equation 27 follows from the fact that the frequency spectrum of derived alleles at $R$ under mutation-drift equilibrium is given by $P(r) = \frac{\theta}{r}$ [Griffiths, 2003]. Equation 28 follows from explicit integration using the analytical solution for $P(s|r, \tau_S)$ [Kimura, 1955, Chen et al., 2007].

$$
\begin{aligned}
f(y) &= \theta \exp(-\tau_S) \int \Big\{ P(u|s, \tau_U) P(w|s, \tau_W) P(y|z, \tau_Y) \mathbf{1}(z = \alpha u + (1-\alpha)w) \Big\} ds du dw dz \\
&= \theta \exp(-\tau_S) \int \Big\{ P(1-u|1-s, \tau_U) P(1-w|1-s, \tau_W) \\
&\qquad P(1-y|1-z, \tau_Y) \mathbf{1}(1-z = \alpha(1-u) + (1-\alpha)(1-w)) \Big\} ds du dw dz \qquad (30) \\
&= \theta \exp(-\tau_S) \int P(u|s, \tau_U) P(w|s, \tau_W) P(1-y|z, \tau_Y) \mathbf{1}(z = \alpha u + (1-\alpha)w) ds du dw dz \qquad (31) \\
&= f(1-y) \qquad (32)
\end{aligned}
$$

Equation 30 uses the property of the transition probability that $P(y|x, \tau) = P(1-y|1-x, \tau)$ while Equation 31 follows by a change of variables.

Combining Equations 32 and 24, we have $f_i = f_{n-i}$ for all $i \in \{1, \ldots, n-1\}$.

Kay Prüfer, Fernando Racimo, Nick Patterson, Flora Jay, Sriram Sankararaman, Susanna Sawyer, Anja Heinze, Gabriel Renaud, Peter H. Sudmant, Cesare de Filippo, Heng Li, Swapan Mallick, Michael Dannemann, Qiaomei Fu, Martin Kircher, Martin Kuhlwilm, Michael Lachmann, Matthias Meyer, Matthias Ongyerth, Michael Siebauer, Christoph Theunert, Arti Tandon, Priya Moorjani, Joseph Pickrell, James C. Mullikin, Samuel H. Vohr, Richard E. Green, Ines Hellmann, Philip L. F. Johnson, Hélène Blanche, Howard Cann, Jacob O. Kitzman, Jay Shendure, Evan E. Eichler, Ed S. Lein, Trygve E. Bakken, Liubov V. Golovanova, Vladimir B. Doronichev, Michael V. Shunkov, Anatoli P. Derevianko, Bence Viola, Montgomery Slatkin, David Reich, Janet Kelso, and Svante Pääbo. The complete genome sequence of a Neanderthal from the Altai Mountains. *Nature*, 505(7481):43–49, January 2014. ISSN 0028-0836. doi: 10.1038/nature12886. URL http://www.nature.com/nature/journal/v505/n7481/abs/nature12886.html.

The 1000 Genomes Project Consortium. A global reference for human genetic variation. *Nature*, 526(7571): 68–74, October 2015. ISSN 0028-0836. doi: 10.1038/nature15393. URL https://www.nature.com/nature/journal/v526/n7571/full/nature15393.html.

Kay Prüfer, Cesare de Filippo, Steffi Grote, Fabrizio Mafessoni, Petra Korlević, Mateja Hajdinjak, Benjamin Vernot, Laurits Skov, Pinghsun Hsieh, Stéphane Peyrégne, David Reher, Charlotte Hopfe, Sarah Nagel, Tomislav Maricic, Qiaomei Fu, Christoph Theunert, Rebekah Rogers, Pontus Skoglund, Manjusha Chintalapati, Michael Dannemann, Bradley J. Nelson, Felix M. Key, Pavao Rudan, Željko Kućan, Ivan Gušić, Liubov V. Golovanova, Vladimir B. Doronichev, Nick Patterson, David Reich, Evan E. Eichler,

Montgomery Slatkin, Mikkel H. Schierup, Aida Andrés, Janet Kelso, Matthias Meyer, and Svante Pääbo. A high-coverage Neandertal genome from Vindija Cave in Croatia. *Science*, page eaao1887, October 2017. ISSN 0036-8075, 1095-9203. doi: 10.1126/science.aao1887. URL http://science.sciencemag.org/content/early/2017/10/04/science.aao1887.

Matthias Meyer, Martin Kircher, Marie-Theres Gansauge, Heng Li, Fernando Racimo, Swapan Mallick, Joshua G. Schraiber, Flora Jay, Kay Prüfer, Cesare de Filippo, Peter H. Sudmant, Can Alkan, Qiaomei Fu, Ron Do, Nadin Rohland, Arti Tandon, Michael Siebauer, Richard E. Green, Katarzyna Bryc, Adrian W. Briggs, Udo Stenzel, Jesse Dabney, Jay Shendure, Jacob Kitzman, Michael F. Hammer, Michael V. Shunkov, Anatoli P. Derevianko, Nick Patterson, Aida M. Andrés, Evan E. Eichler, Montgomery Slatkin, David Reich, Janet Kelso, and Svante Pääbo. A High-Coverage Genome Sequence from an Archaic Denisovan Individual. *Science*, 338(6104):222–226, October 2012. ISSN 0036-8075, 1095-9203. doi: 10.1126/science.1224344. URL http://science.sciencemag.org/content/338/6104/222.

The Chimpanzee Sequencing and Analysis Consortium, Robert H. Waterson, Eric S. Lander, and Richard K. Wilson. Initial sequence of the chimpanzee genome and comparison with the human genome. *Nature*, 437 (7055):69–87, September 2005. ISSN 1476-4687. doi: 10.1038/nature04072. URL https://www.nature.com/articles/nature04072.

Devin P. Locke, LaDeana W. Hillier, Wesley C. Warren, Kim C. Worley, Lynne V. Nazareth, Donna M. Muzny, Shiaw-Pyng Yang, Zhengyuan Wang, Asif T. Chinwalla, Pat Minx, Makedonka Mitreva, Lisa Cook, Kim D. Delehaunty, Catrina Fronick, Heather Schmidt, Lucinda A. Fulton, Robert S. Fulton, Joanne O. Nelson, Vincent Magrini, Craig Pohl, Tina A. Graves, Chris Markovic, Andy Cree, Huyen H. Dinh, Jennifer Hume, Christie L. Kovar, Gerald R. Fowler, Gerton Lunter, Stephen Meader, Andreas Heger, Chris P. Ponting, Tomas Marques-Bonet, Can Alkan, Lin Chen, Ze Cheng, Jeffrey M. Kidd, Evan E. Eichler, Simon White, Stephen Searle, Albert J. Vilella, Yuan Chen, Paul Flicek, Jian Ma, Brian Raney, Bernard Suh, Richard Burhans, Javier Herrero, David Haussler, Rui Faria, Olga Fernando, Fleur Darré, Domènec Farré, Elodie Gazave, Meritxell Oliva, Arcadi Navarro, Roberta Roberto, Oronzo Capozzi, Nicoletta Archidiacono, Giuliano Della Valle, Stefania Purgato, Mariano Rocchi, Miriam K. Konkel, Jerilyn A. Walker, Brygg Ullmer, Mark A. Batzer, Arian F. A. Smit, Robert Hubley, Claudio Casola, Daniel R. Schrider, Matthew W. Hahn, Victor Quesada, Xose S. Puente, Gonzalo R. Ordoñez, Carlos López-Otín, Tomas Vinar, Brona Brejova, Aakrosh Ratan, Robert S. Harris, Webb Miller, Carolin Kosiol, Heather A. Lawson, Vikas Taliwal, André L. Martins, Adam Siepel, Arindam RoyChoudhury, Xin Ma, Jeremiah Degenhardt, Carlos D. Bustamante, Ryan N. Gutenkunst, Thomas Mailund, Julien Y. Dutheil, Asger Hobolth, Mikkel H. Schierup, Oliver A. Ryder, Yuko Yoshinaga, Pieter J. de Jong, George M. Weinstock, Jeffrey Rogers, Elaine R. Mardis, Richard A. Gibbs, and Richard K. Wilson. Comparative and demographic analysis of orang-utan genomes. *Nature*, 469(7331):529–533, January 2011. ISSN 1476-4687. doi: 10.1038/nature09687. URL https://www.nature.com/articles/nature09687.

Sriram Sankararaman, Nick Patterson, Heng Li, Svante Pääbo, and David Reich. The Date of Interbreeding between Neandertals and Modern Humans. *PLOS Genetics*, 8(10):e1002947, October 2012. ISSN 1553-7404. doi: 10.1371/journal.pgen.1002947. URL http://journals.plos.org/plosgenetics/article?id=10.1371/journal.pgen.1002947.

Qiaomei Fu, Mateja Hajdinjak, Oana Teodora Moldovan, Silviu Constantin, Swapan Mallick, Pontus Skoglund, Nick Patterson, Nadin Rohland, Iosif Lazaridis, Birgit Nickel, Bence Viola, Kay Prüfer, Matthias Meyer, Janet Kelso, David Reich, and Svante Pääbo. An early modern human from Romania with a recent Neanderthal ancestor. *Nature*, 524(7564):216–219, August 2015. ISSN 1476-4687. doi: 10.1038/nature14558. URL https://www.nature.com/articles/nature14558.

G. A. Watterson. On the number of segregating sites in genetical models without recombination. *Theoretical Population Biology*, 7(2):256–276, April 1975. ISSN 0040-5809. doi: 10.1016/0040-5809(75)90020-9. URL http://www.sciencedirect.com/science/article/pii/0040580975900209.

International HapMap Consortium, Kelly A. Frazer, Dennis G. Ballinger, David R. Cox, David A. Hinds, Laura L. Stuve, Richard A. Gibbs, John W. Belmont, Andrew Boudreau, Paul Hardenbol, Suzanne M. Leal, Shiran Pasternak, David A. Wheeler, Thomas D. Willis, Fuli Yu, Huanming Yang, Changqing Zeng,

Yang Gao, Haoran Hu, Weitao Hu, Chaohua Li, Wei Lin, Siqi Liu, Hao Pan, Xiaoli Tang, Jian Wang, Wei Wang, Jun Yu, Bo Zhang, Qingrun Zhang, Hongbin Zhao, Hui Zhao, Jun Zhou, Stacey B. Gabriel, Rachel Barry, Brendan Blumenstiel, Amy Camargo, Matthew Defelice, Maura Faggart, Mary Goyette, Supriya Gupta, Jamie Moore, Huy Nguyen, Robert C. Onofrio, Melissa Parkin, Jessica Roy, Erich Stahl, Ellen Winchester, Liuda Ziaugra, David Altshuler, Yan Shen, Zhijian Yao, Wei Huang, Xun Chu, Yungang He, Li Jin, Yangfan Liu, Yayun Shen, Weiwei Sun, Haifeng Wang, Yi Wang, Ying Wang, Xiaoyan Xiong, Liang Xu, Mary M. Y. Waye, Stephen K. W. Tsui, Hong Xue, J. Tze-Fei Wong, Luana M. Galver, Jian-Bing Fan, Kevin Gunderson, Sarah S. Murray, Arnold R. Oliphant, Mark S. Chee, Alexandre Montpetit, Fanny Chagnon, Vincent Ferretti, Martin Leboeuf, Jean-François Olivier, Michael S. Phillips, Stéphanie Roumy, Clémentine Sallée, Andrei Verner, Thomas J. Hudson, Pui-Yan Kwok, Dongmei Cai, Daniel C. Koboldt, Raymond D. Miller, Ludmila Pawlikowska, Patricia Taillon-Miller, Ming Xiao, Lap-Chee Tsui, William Mak, You Qiang Song, Paul K. H. Tam, Yusuke Nakamura, Takahisa Kawaguchi, Takuya Kitamoto, Takashi Morizono, Atsushi Nagashima, Yozo Ohnishi, Akihiro Sekine, Toshihiro Tanaka, Tatsuhiko Tsunoda, Panos Deloukas, Christine P. Bird, Marcos Delgado, Emmanouil T. Dermitzakis, Rhian Gwilliam, Sarah Hunt, Jonathan Morrison, Don Powell, Barbara E. Stranger, Pamela Whittaker, David R. Bentley, Mark J. Daly, Paul I. W. de Bakker, Jeff Barrett, Yves R. Chretien, Julian Maller, Steve McCarroll, Nick Patterson, Itsik Pe'er, Alkes Price, Shaun Purcell, Daniel J. Richter, Pardis Sabeti, Richa Saxena, Stephen F. Schaffner, Pak C. Sham, Patrick Varilly, David Altshuler, Lincoln D. Stein, Lalitha Krishnan, Albert Vernon Smith, Marcela K. Tello-Ruiz, Gudmundur A. Thorisson, Aravinda Chakravarti, Peter E. Chen, David J. Cutler, Carl S. Kashuk, Shin Lin, Gonçalo R. Abecasis, Weihua Guan, Yun Li, Heather M. Munro, Zhaohui Steve Qin, Daryl J. Thomas, Gilean McVean, Adam Auton, Leonardo Bottolo, Niall Cardin, Susana Eyheramendy, Colin Freeman, Jonathan Marchini, Simon Myers, Chris Spencer, Matthew Stephens, Peter Donnelly, Lon R. Cardon, Geraldine Clarke, David M. Evans, Andrew P. Morris, Bruce S. Weir, Tatsuhiko Tsunoda, James C. Mullikin, Stephen T. Sherry, Michael Feolo, Andrew Skol, Houcan Zhang, Changqing Zeng, Hui Zhao, Ichiro Matsuda, Yoshimitsu Fukushima, Darryl R. Macer, Eiko Suda, Charles N. Rotimi, Clement A. Adebamowo, Ike Ajayi, Toyin Aniagwu, Patricia A. Marshall, Chibuzor Nkwodimmah, Charmaine D. M. Royal, Mark F. Leppert, Missy Dixon, Andy Peiffer, Renzong Qiu, Alastair Kent, Kazuto Kato, Norio Niikawa, Isaac F. Adewole, Bartha M. Knoppers, Morris W. Foster, Ellen Wright Clayton, Jessica Watkin, Richard A. Gibbs, John W. Belmont, Donna Muzny, Lynne Nazareth, Erica Sodergren, George M. Weinstock, David A. Wheeler, Imtaz Yakub, Stacey B. Gabriel, Robert C. Onofrio, Daniel J. Richter, Liuda Ziaugra, Bruce W. Birren, Mark J. Daly, David Altshuler, Richard K. Wilson, Lucinda L. Fulton, Jane Rogers, John Burton, Nigel P. Carter, Christopher M. Clee, Mark Griffiths, Matthew C. Jones, Kirsten McLay, Robert W. Plumb, Mark T. Ross, Sarah K. Sims, David L. Willey, Zhu Chen, Hua Han, Le Kang, Martin Godbout, John C. Wallenburg, Paul L'Archevêque, Guy Bellemare, Koji Saeki, Hongguang Wang, Daochang An, Hongbo Fu, Qing Li, Zhen Wang, Renwu Wang, Arthur L. Holden, Lisa D. Brooks, Jean E. McEwen, Mark S. Guyer, Vivian Ota Wang, Jane L. Peterson, Michael Shi, Jack Spiegel, Lawrence M. Sung, Lynn F. Zacharia, Francis S. Collins, Karen Kennedy, Ruth Jamieson, and John Stewart. A second generation human haplotype map of over 3.1 million SNPs. *Nature*, 449(7164):851–861, October 2007. ISSN 1476-4687. doi: 10.1038/nature06258.

Simon Gravel, Brenna M. Henn, Ryan N. Gutenkunst, Amit R. Indap, Gabor T. Marth, Andrew G. Clark, Fuli Yu, Richard A. Gibbs, The 1000 Genomes Project, and Carlos D. Bustamante. Demographic history and rare allele sharing among human populations. *Proceedings of the National Academy of Sciences*, 108 (29):11983–11988, July 2011. ISSN 0027-8424, 1091-6490. doi: 10.1073/pnas.1019276108. URL `http://www.pnas.org/content/108/29/11983`.

Kelley Harris and Rasmus Nielsen. Inferring Demographic History from a Spectrum of Shared Haplotype Lengths. *PLOS Genetics*, 9(6):e1003521, June 2013. ISSN 1553-7404. doi: 10.1371/journal.pgen.1003521. URL `http://journals.plos.org/plosgenetics/article?id=10.1371/journal.pgen.1003521`.

Martin Petr, Svante Pääbo, Janet Kelso, and Benjamin Vernot. Limits of long-term selection against Neandertal introgression. *Proceedings of the National Academy of Sciences*, page 201814338, January 2019. ISSN 0027-8424, 1091-6490. doi: 10.1073/pnas.1814338116. URL `https://www.pnas.org/content/early/2019/01/14/1814338116`.

PingHsun Hsieh, Krishna R. Veeramah, Joseph Lachance, Sarah A. Tishkoff, Jeffrey D. Wall, Michael F.

Hammer, and Ryan N. Gutenkunst. Whole-genome sequence analyses of Western Central African Pygmy hunter-gatherers reveal a complex demographic history and identify candidate genes under positive natural selection. *Genome Research*, February 2016. ISSN 1088-9051, 1549-5469. doi: 10.1101/gr.192971.115. URL http://genome.cshlp.org/content/early/2016/02/08/gr.192971.115.

Benedict Paten, Javier Herrero, Stephen Fitzgerald, Kathryn Beal, Paul Flicek, Ian Holmes, and Ewan Birney. Genome-wide nucleotide-level mammalian ancestor reconstruction. *Genome Research*, 18(11): 1829–1843, November 2008. ISSN 1088-9051, 1549-5469. doi: 10.1101/gr.076521.108. URL http://genome.cshlp.org/content/18/11/1829.

Kay Prüfer. snpAD: an ancient DNA genotype caller. *Bioinformatics*, 34(24):4165–4171, December 2018. ISSN 1367-4803. doi: 10.1093/bioinformatics/bty507. URL https://academic.oup.com/bioinformatics/article/34/24/4165/5042170.

Melinda A. Yang, Anna-Sapfo Malaspinas, Eric Y. Durand, and Montgomery Slatkin. Ancient structure in Africa unlikely to explain Neanderthal and non-African genetic similarity. *Molecular Biology and Evolution*, 29(10):2987–2995, October 2012. ISSN 1537-1719. doi: 10.1093/molbev/mss117.

Richard R. Hudson. Generating samples under a Wright–Fisher neutral model of genetic variation. *Bioinformatics*, 18(2):337–338, February 2002. ISSN 1367-4803. doi: 10.1093/bioinformatics/18.2.337. URL https://academic.oup.com/bioinformatics/article/18/2/337/225783.

Jack N. Fenner. Cross-cultural estimation of the human generation interval for use in genetics-based population divergence studies. *American Journal of Physical Anthropology*, 128(2):415–423, 2005. ISSN 1096-8644. doi: 10.1002/ajpa.20188. URL https://onlinelibrary.wiley.com/doi/abs/10.1002/ajpa.20188.

Katalin Csilléry, Olivier François, and Michael G. B. Blum. abc: an R package for approximate Bayesian computation (ABC). *Methods in Ecology and Evolution*, 3(3):475–479, June 2012. ISSN 2041-210X. doi: 10.1111/j.2041-210X.2011.00179.x. URL https://besjournals.onlinelibrary.wiley.com/doi/abs/10.1111/j.2041-210X.2011.00179.x.

Brenna M Henn, Teresa E Steele, and Timothy D Weaver. Clarifying distinct models of modern human origins in Africa. *Current Opinion in Genetics & Development*, 53:148–156, December 2018. ISSN 0959-437X. doi: 10.1016/j.gde.2018.10.003. URL http://www.sciencedirect.com/science/article/pii/S0959437X1730182X.

Arun Durvasula and Sriram Sankararaman. Recovering signals of ghost archaic admixture in the genomes of present-day Africans. *bioRxiv*, page 285734, March 2018. doi: 10.1101/285734. URL https://www.biorxiv.org/content/early/2018/03/21/285734.

Vincent Plagnol and Jeffrey D. Wall. Possible Ancestral Structure in Human Populations. *PLOS Genetics*, 2(7):e105, July 2006. ISSN 1553-7404. doi: 10.1371/journal.pgen.0020105. URL http://journals.plos.org/plosgenetics/article?id=10.1371/journal.pgen.0020105.

Hans R. Kunsch. The Jackknife and the Bootstrap for General Stationary Observations. *The Annals of Statistics*, 17(3):1217–1241, September 1989. ISSN 0090-5364, 2168-8966. doi: 10.1214/aos/1176347265. URL https://projecteuclid.org/euclid.aos/1176347265.

Stephan Schiffels and Richard Durbin. Inferring human population size and separation history from multiple genome sequences. *Nature Genetics*, 46(8):919–925, August 2014. ISSN 1546-1718. doi: 10.1038/ng.3015. URL https://www.nature.com/articles/ng.3015.

Aaron R. Quinlan and Ira M. Hall. BEDTools: a flexible suite of utilities for comparing genomic features. *Bioinformatics*, 26(6):841–842, March 2010. ISSN 1367-4803. doi: 10.1093/bioinformatics/btq033. URL https://academic.oup.com/bioinformatics/article/26/6/841/244688.

Adam Frankish, Mark Diekhans, Anne-Maud Ferreira, Rory Johnson, Irwin Jungreis, Jane Loveland, Jonathan M. Mudge, Cristina Sisu, James Wright, Joel Armstrong, If Barnes, Andrew Berry, Alexandra Bignell, Silvia Carbonell Sala, Jacqueline Chrast, Fiona Cunningham, Tomás Di Domenico, Sarah Donaldson, Ian T. Fiddes, Carlos García Girón, Jose Manuel Gonzalez, Tiago Grego, Matthew Hardy, Thibaut Hourlier, Toby Hunt, Osagie G. Izuogu, Julien Lagarde, Fergal J. Martin, Laura Martínez, Shamika Mohanan, Paul Muir, Fabio C. P. Navarro, Anne Parker, Baikang Pei, Fernando Pozo, Magali Ruffier, Bianca M. Schmitt, Eloise Stapleton, Marie-Marthe Suner, Irina Sycheva, Barbara Uszczynska-Ratajczak, Jinuri Xu, Andrew Yates, Daniel Zerbino, Yan Zhang, Bronwen Aken, Jyoti S. Choudhary, Mark Gerstein, Roderic Guigó, Tim J. P. Hubbard, Manolis Kellis, Benedict Paten, Alexandre Reymond, Michael L. Tress, and Paul Flicek. GENCODE reference annotation for the human and mouse genomes. *Nucleic Acids Research*, 47(D1):D766–D773, January 2019. ISSN 1362-4962. doi: 10.1093/nar/gky955.

Jeffrey D. Wall, Melinda A. Yang, Flora Jay, Sung K. Kim, Eric Y. Durand, Laurie S. Stevison, Christopher Gignoux, August Woerner, Michael F. Hammer, and Montgomery Slatkin. Higher Levels of Neanderthal Ancestry in East Asians than in Europeans. *Genetics*, 194(1):199–209, May 2013. ISSN 0016-6731, 1943-2631. doi: 10.1534/genetics.112.148213. URL `http://www.genetics.org/content/194/1/199`.

Swapan Mallick, Heng Li, Mark Lipson, Iain Mathieson, Melissa Gymrek, Fernando Racimo, Mengyao Zhao, Niru Chennagiri, Susanne Nordenfelt, Arti Tandon, Pontus Skoglund, Iosif Lazaridis, Sriram Sankararaman, Qiaomei Fu, Nadin Rohland, Gabriel Renaud, Yaniv Erlich, Thomas Willems, Carla Gallo, Jeffrey P. Spence, Yun S. Song, Giovanni Poletti, Francois Balloux, George van Driem, Peter de Knijff, Irene Gallego Romero, Aashish R. Jha, Doron M. Behar, Claudio M. Bravi, Cristian Capelli, Tor Hervig, Andres Moreno-Estrada, Olga L. Posukh, Elena Balanovska, Oleg Balanovsky, Sena Karachanak-Yankova, Hovhannes Sahakyan, Draga Toncheva, Levon Yepiskoposyan, Chris Tyler-Smith, Yali Xue, M. Syafiq Abdullah, Andres Ruiz-Linares, Cynthia M. Beall, Anna Di Rienzo, Choongwon Jeong, Elena B. Starikovskaya, Ene Metspalu, Jüri Parik, Richard Villems, Brenna M. Henn, Ugur Hodoglugil, Robert Mahley, Antti Sajantila, George Stamatoyannopoulos, Joseph T. S. Wee, Rita Khusainova, Elza Khusnutdinova, Sergey Litvinov, George Ayodo, David Comas, Michael F. Hammer, Toomas Kivisild, William Klitz, Cheryl A. Winkler, Damian Labuda, Michael Bamshad, Lynn B. Jorde, Sarah A. Tishkoff, W. Scott Watkins, Mait Metspalu, Stanislav Dryomov, Rem Sukernik, Lalji Singh, Kumarasamy Thangaraj, Svante Pääbo, Janet Kelso, Nick Patterson, and David Reich. The Simons Genome Diversity Project: 300 genomes from 142 diverse populations. *Nature*, 538(7624):201, October 2016. ISSN 1476-4687. doi: 10.1038/nature18964. URL `https://www.nature.com/articles/nature18964`.

Motoo Kimura. Solution of a Process of Random Genetic Drift with a Continuous Model. *Proceedings of the National Academy of Sciences*, 41(3):144–150, March 1955. ISSN 0027-8424, 1091-6490. doi: 10.1073/pnas.41.3.144. URL `https://www.pnas.org/content/41/3/144`.

R. C. Griffiths. The frequency spectrum of a mutation, and its age, in a general diffusion model. *Theoretical Population Biology*, 64(2):241–251, September 2003. ISSN 0040-5809. doi: 10.1016/S0040-5809(03)00075-3. URL `http://www.sciencedirect.com/science/article/pii/S0040580903000753`.

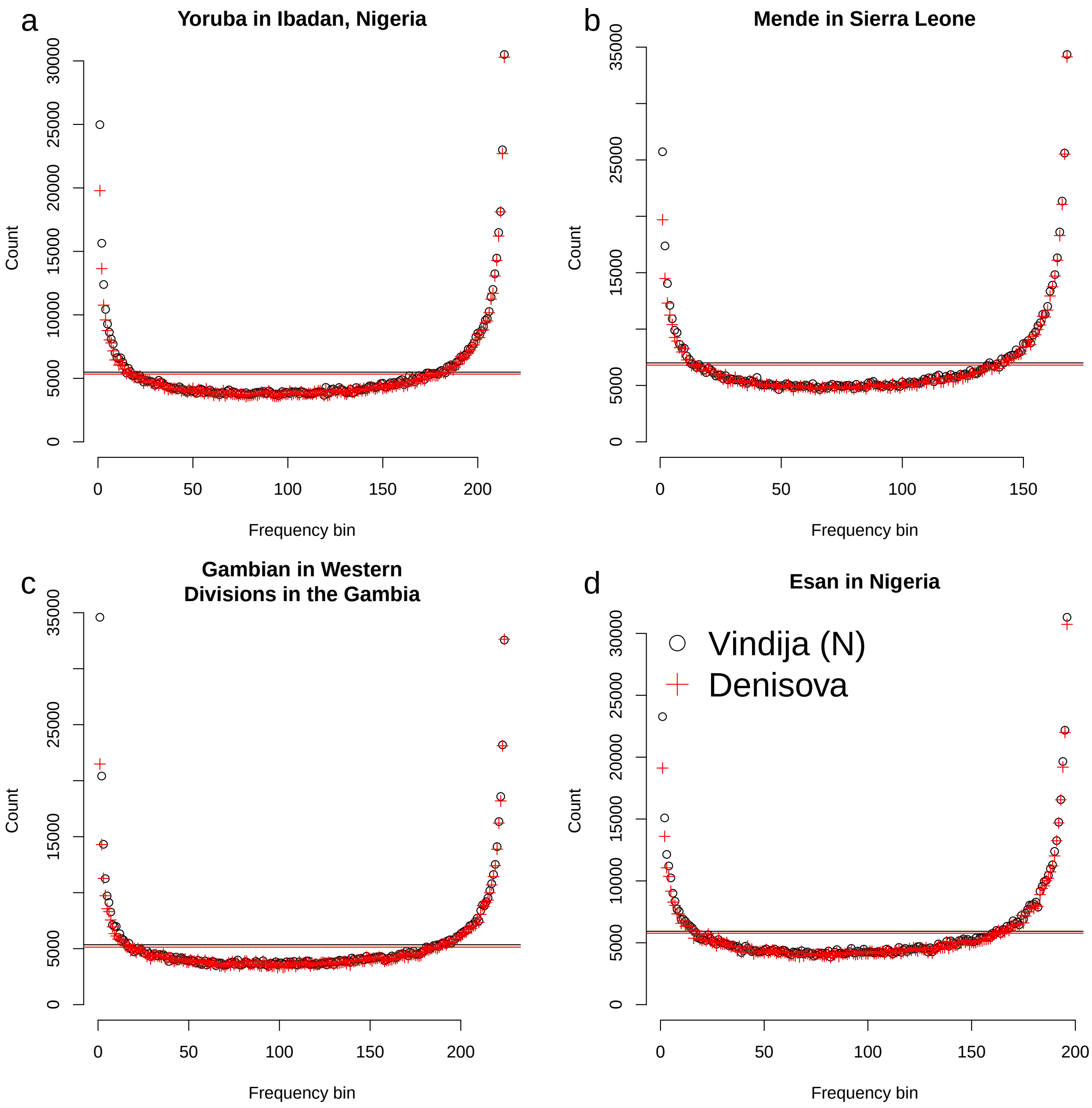

Figure S4: CSFS from 1000G Phase 3 data across all African populations included in the dataset. Black circles denote the CSFS conditioned on the Vindija Neanderthal [Prüfer et al., 2017] and red pluses denote the CSFS conditioned on the high coverage Denisovan [Meyer et al., 2012].

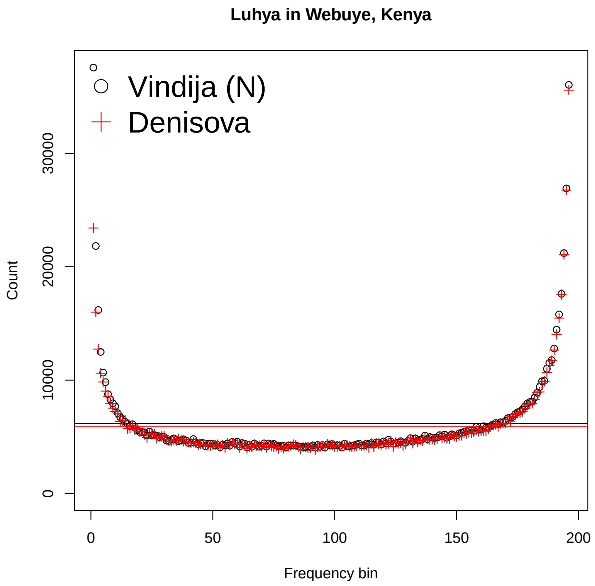

Figure S5: CSFS from 1000G Phase 3 data the Luhya population. Black circles denote the CSFS conditioned on the Vindija Neanderthal [Prüfer et al., 2017] and red pluses denote the CSFS conditioned on the high coverage Denisovan [Meyer et al., 2012].

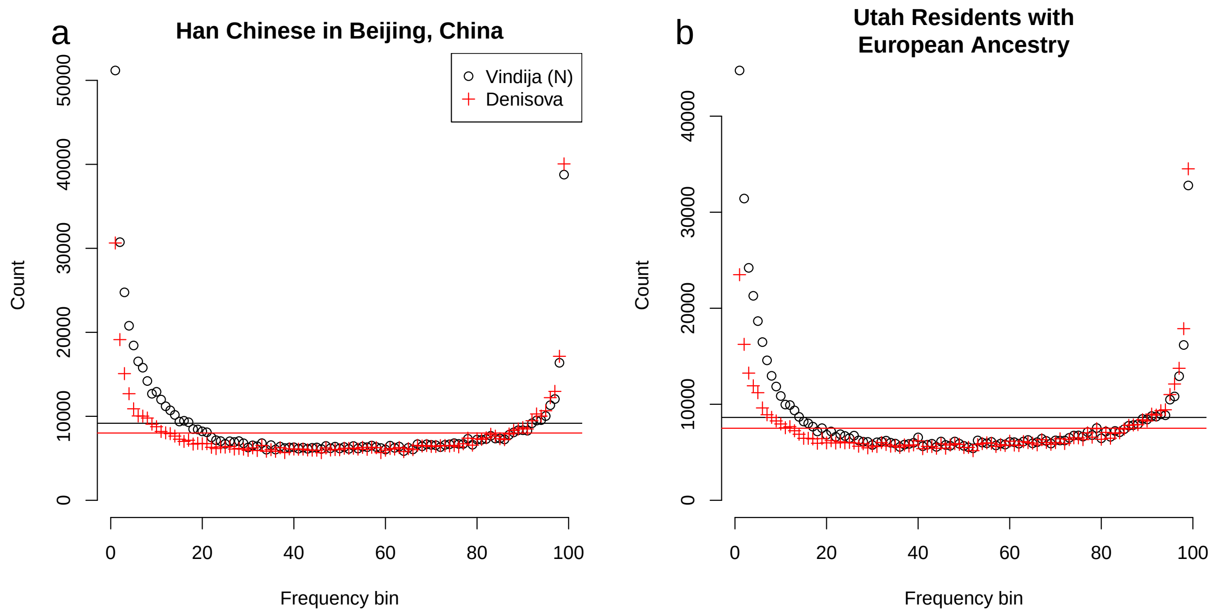

Figure S6: Conditional site frequency spectra (CSFS) from 1000 Genomes Phase 3 data. We show the number of sites in each frequency class in a sample of 50 individuals from each population, conditioned on observing a derived mutation in the Vindija Neanderthal (black points) and the Denisova (red plus). The lines denote the mean counts, which is the expectation under panmixia. (a) Han Chinese in Beijing (CHB) (b)European (CEU)

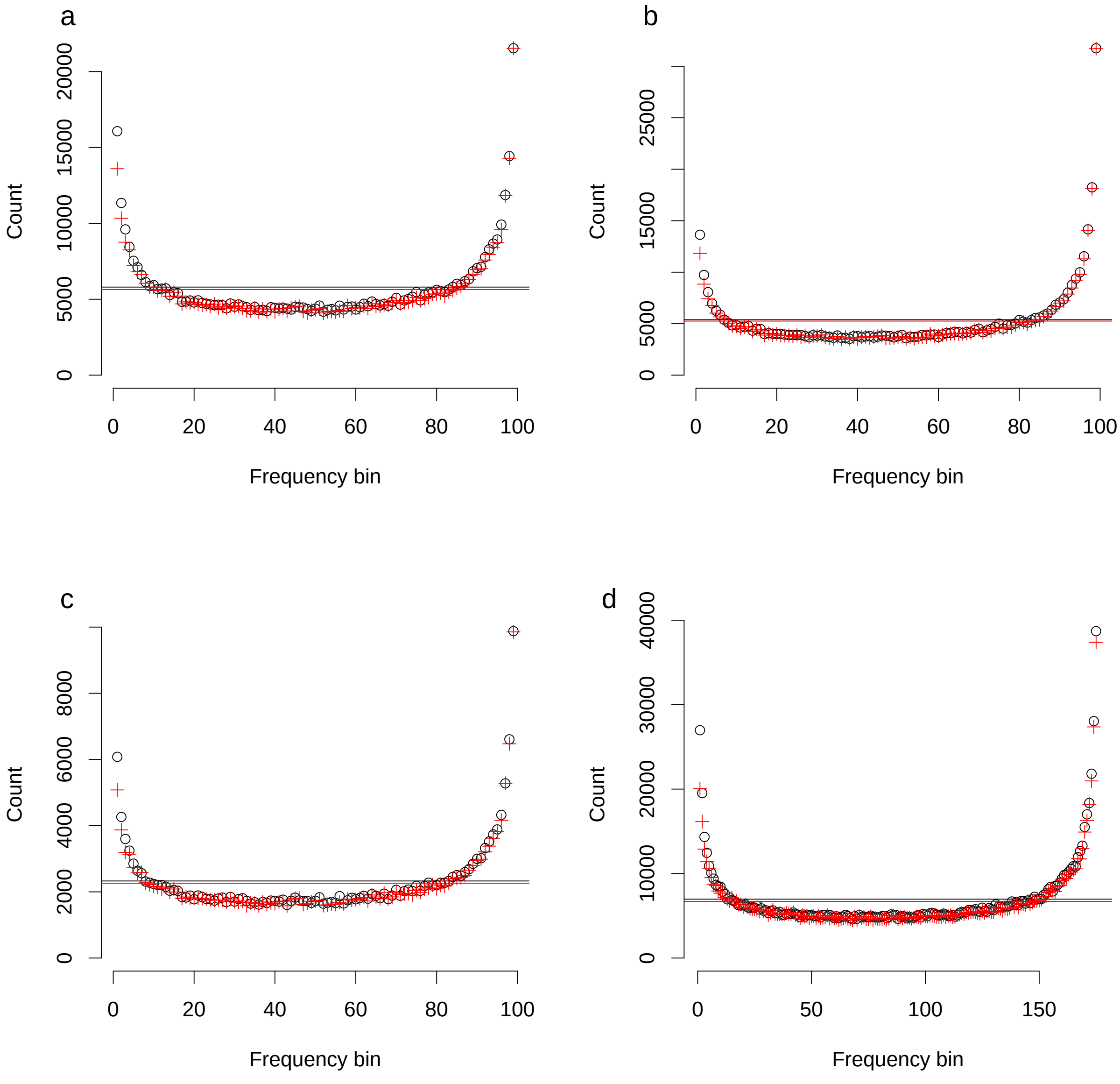

Figure S7: CSFS calculated for YRI with (a) CpG sites removed (b) transitions only (c) transversions only (d) Phase 1 1000 Genomes data (88 diploid individuals). Black circles denote the CSFS conditioned on the Vindija Neanderthal [Prüfer et al., 2017] and red pluses denote the CSFS conditioned on the high coverage Denisovan [Meyer et al., 2012].

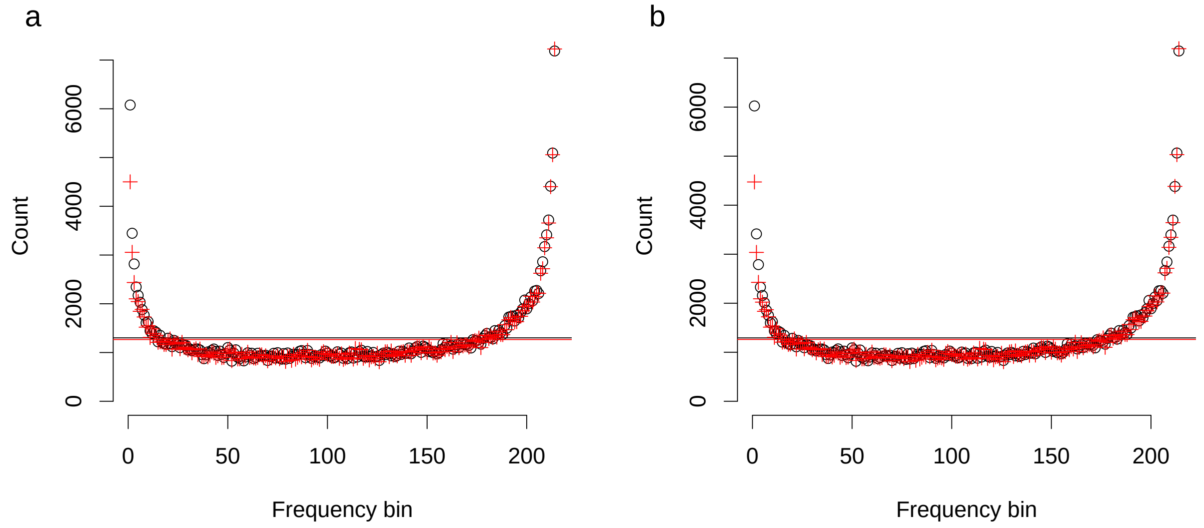

Figure S8: The CSFS in YRI restricted to sites where the archaic (Neanderthal or Denisovan) has a genotype quality (GQ) (a) $\geq 30$ (b) $\geq 50$. Black circles denote the CSFS conditioned on the Vindija Neanderthal [Prüfer et al., 2017] and red pluses denote the CSFS conditioned on the high coverage Denisovan [Meyer et al., 2012].

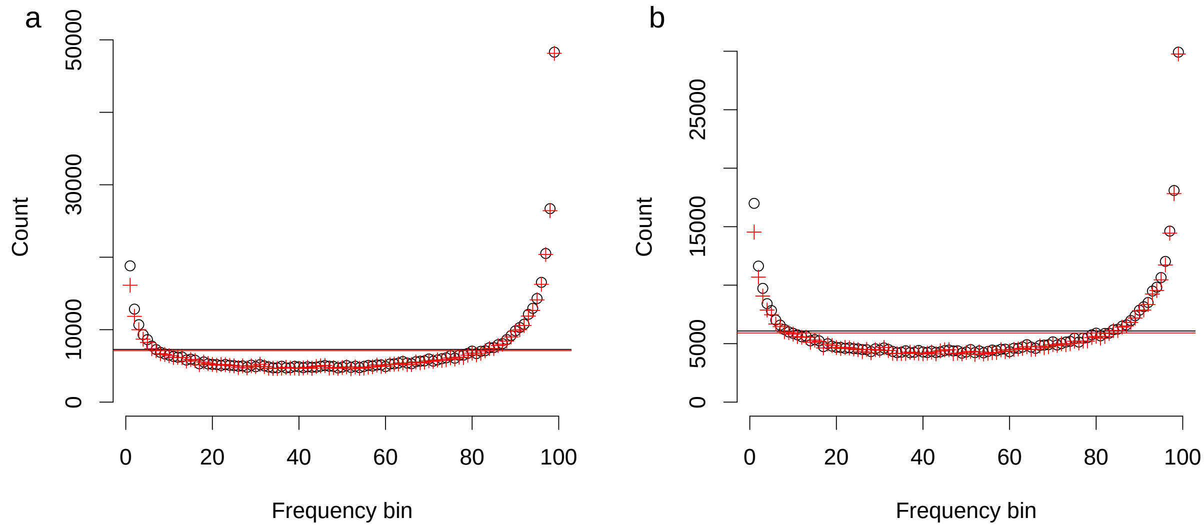

Figure S9: (a) The CSFS in YRI using the chimpanzee genome to polarize the ancestral allele. (b) The CSFS in YRI restricting to sites where the chimpanzee and orangutan genomes have the same allele, which is used as the ancestral allele. These results are nearly identical to results using the EPO alignment to polarize the ancestral allele although the total number of sites is reduced. Black circles denote the CSFS conditioned on the Vindija Neanderthal [Prüfer et al., 2017] and red pluses denote the CSFS conditioned on the high coverage Denisovan [Meyer et al., 2012].

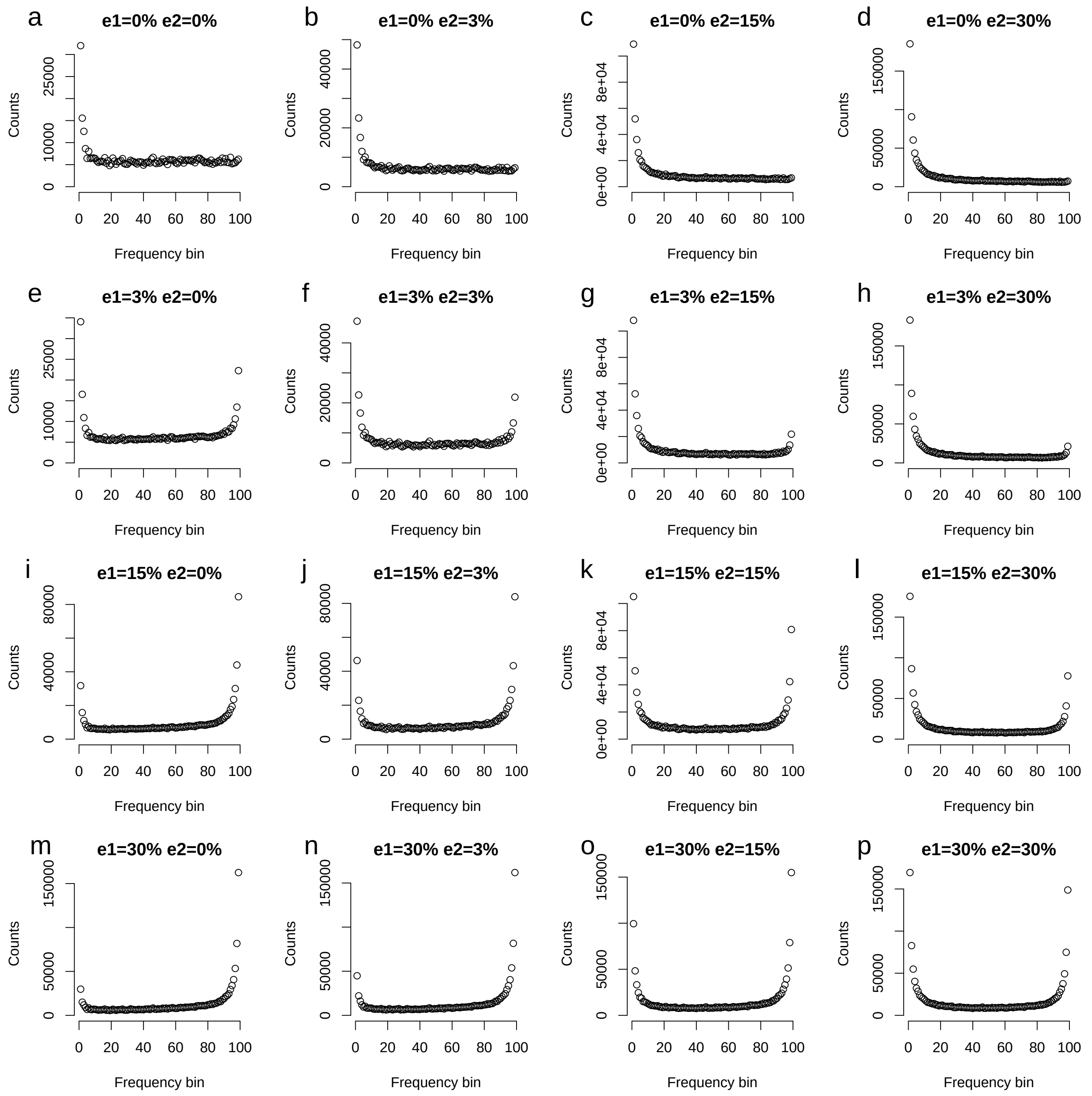

Figure S10: Simulations of the baseline model (Section S2) with both ancestral misidentification (e1) and genotyping error in the archaic (e2). Model likelihoods are presented in table S2

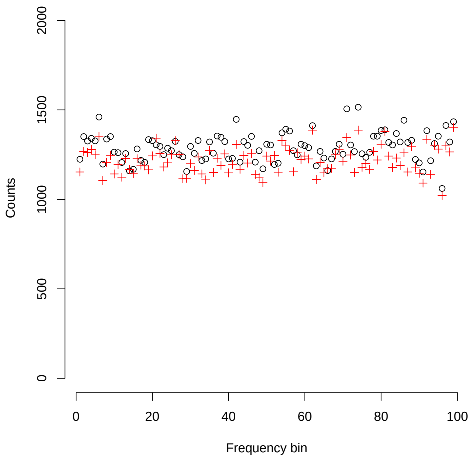

Figure S11: Mutation rate and recombination rate variation: We simulated data from the base model (Section S2) in 1 MB chunks and varied the mutation rate and recombination rate according to mutation rates estimated using Watterson's $\theta$ (See Section S2) and recombination rates from International HapMap Consortium et al. [2007]. Black circles denote the CSFS conditioned on the Vindija Neanderthal [Prüfer et al., 2017] and red pluses denote the CSFS conditioned on the high coverage Denisovan [Meyer et al., 2012].

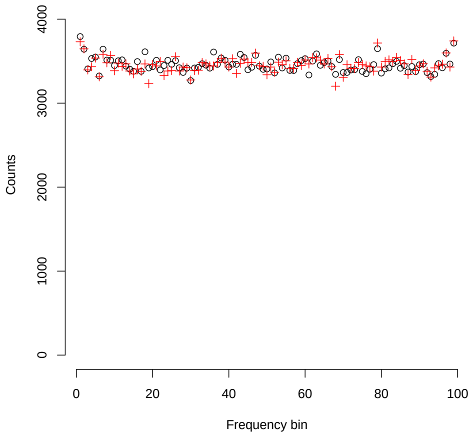

Figure S12: Simulations of the demographic model inferred from Hsieh et al. [2016] relating the Yoruba, Baka, and Biaka populations. This model has gene flow between the three populations dating back to $\simeq$ 40Ky B.P. This gene flow is insufficient to produce the U-shape of the spectrum we observe in the data. Black circles denote the CSFS conditioned on the Vindija Neanderthal [Prüfer et al., 2017] and red pluses denote the CSFS conditioned on the high coverage Denisovan [Meyer et al., 2012].

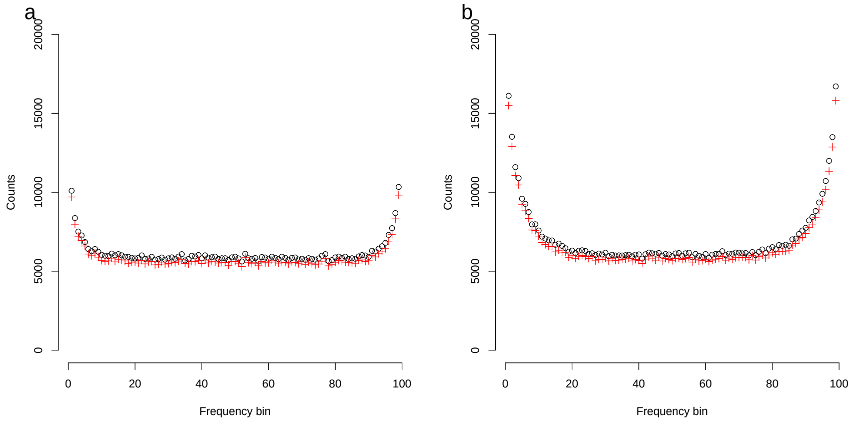

Figure S13: A model with continuous migration between lineages in Africa over the last (a) 6,000 generations (≃175Ky B.P.). This model is similar to the Single Origin Range expansion with Regional Persistence model from Henn et al. [2018]. (b) 12,000 generations (≃350Ky B.P.) This model is similar to the African Multiregionalism model in Henn et al. [2018]. Black circles denote the CSFS conditioned on the Vindija Neanderthal [Prüfer et al., 2017] and red pluses denote the CSFS conditioned on the high coverage Denisovan [Meyer et al., 2012].

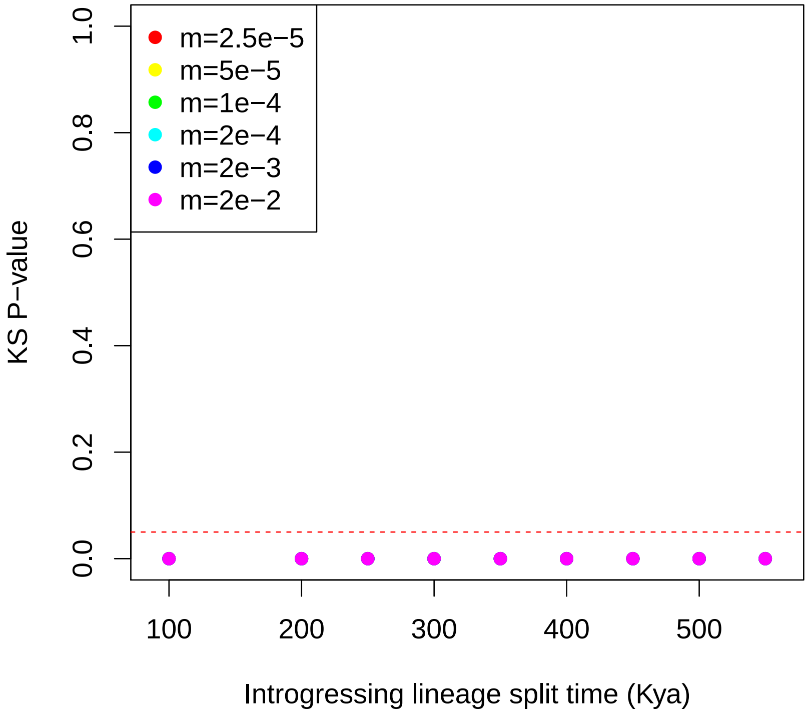

Figure S14: Models with continuous migration ($m$ in units of migrants per generation) since the introgressing lineages lineage splits. We compute a $p-$value assessing the model fit to the data (See Section S1.5). The red line denotes a 0.05 cutoff. A non-significant $p-$value indicates that we cannot reject the null hypothesis that the residuals are normally distributed, indicating a good fit of the model to the data. We see that a model with continuous migration does not fit well.

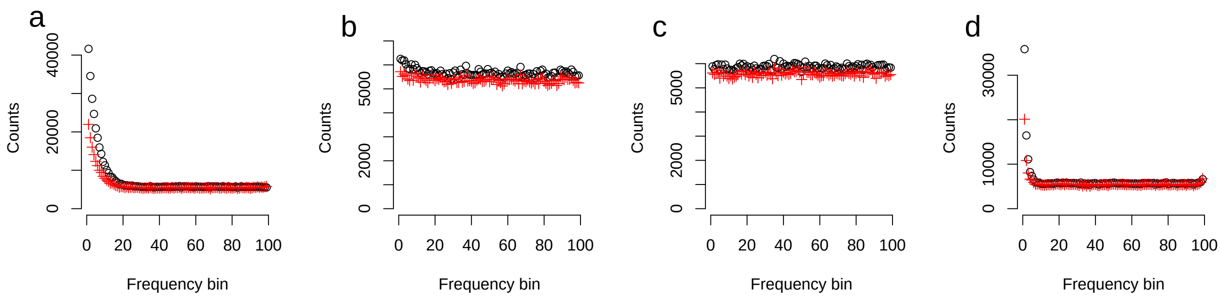

Figure S15: Current demographic models from the literature cannot explain the shape of the CSFS observed in Figure S4. (a) The model from Prüfer et al. [2017], with 1% admixture from the Neanderthal branch into Yoruba 43Ky B.P. (b) the same model, but with a 0.015% admixture pulse, which is the best fitting model in Prüfer et al. [2017] (S9b). (c) The model from Gravel et al. [2011] with archaic lineages and admixture events included (d) The model from Petr et al. [2019], which introduces Neanderthal DNA into Yoruba via migration between Europeans and Yorubans over the last 20Ky B.P. Black circles denote the CSFS conditioned on the Vindija Neanderthal [Prüfer et al., 2017] and red pluses denote the CSFS conditioned on the high coverage Denisovan [Meyer et al., 2012].

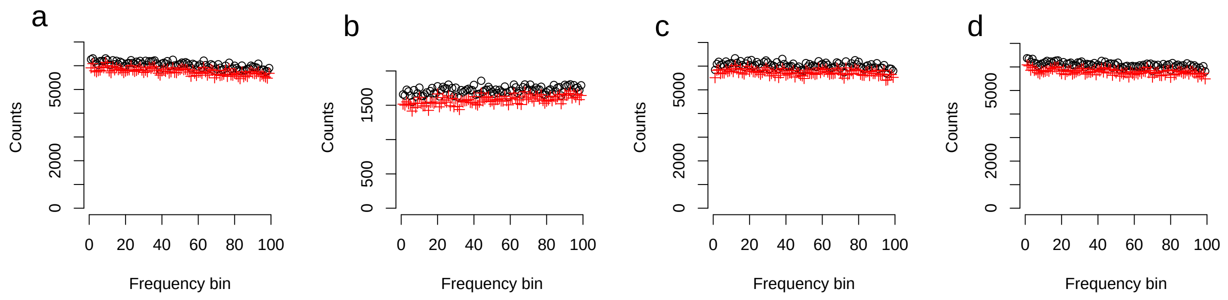

Figure S16: Models involving structure in the ancestor of modern humans and archaics cannot explain the observed CSFS. (a) $5N_a$ population expansion in the ancestor of modern humans and archaics. (b) a $0.2N_a$ bottleneck in the ancestor. (c) Population subdivision (840Ky B.P.) and subsequent merging in the ancestor at 550Ky B.P. (d) Population subdivision (840Ky B.P.) and continuous migration in the ancestor until 550Ky B.P. Black circles denote the CSFS conditioned on the Vindija Neanderthal [Prüfer et al., 2017] and red pluses denote the CSFS conditioned on the high coverage Denisovan [Meyer et al., 2012].

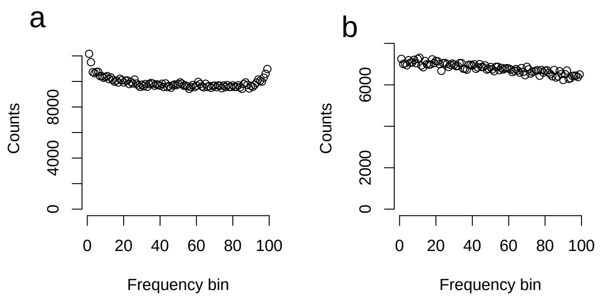

Figure S17: Models involving ancestral structure from the literature cannot explain the observed CSFS. (a) the 'Ancient structure model' (Figure 1B) from Yang et al. [2012], with continuous gene flow between Neanderthals and modern humans. (b) the 'Ancient structure model' from Sankararaman et al. [2012] with continuous gene flow between Neanderthals and modern humans.

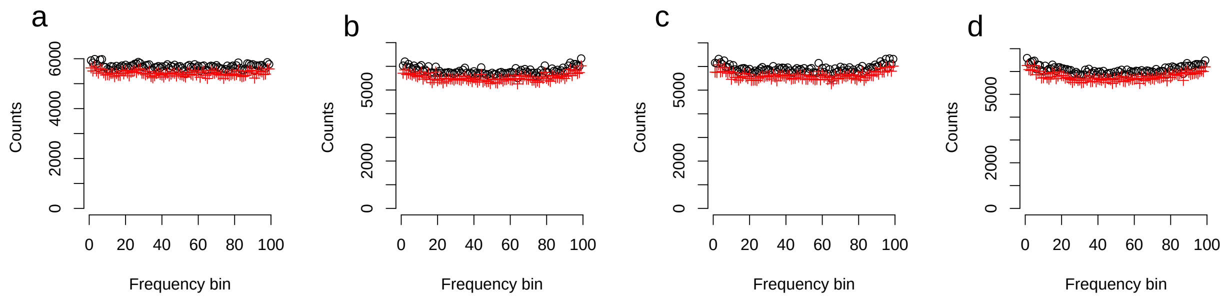

Figure S18: Model A.1: Gene flow from the modern human ancestor branch back into the modern human ancestor prior to the out of Africa event. See section S4.1 for a detailed description. Panels show different amounts of gene flow into the modern human ancestor (a) 1% (b) 5% (c) 10% (d) 20%. Black circles denote the CSFS conditioned on the Vindija Neanderthal [Prüfer et al., 2017] and red pluses denote the CSFS conditioned on the high coverage Denisovan [Meyer et al., 2012].

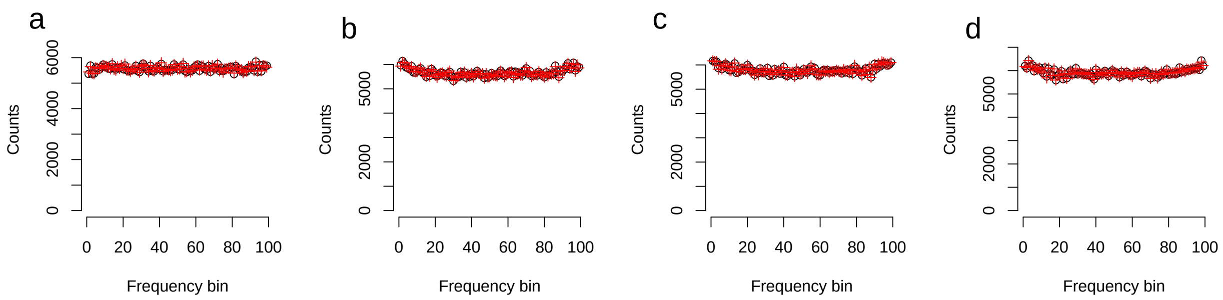

Figure S19: Model sA.1: We simulate the model in Figure S18, but only include the African, Neanderthal, and Denisovan branches (a) 1% (b) 5% (c) 10% (d) 20%. Black circles denote the CSFS conditioned on the Vindija Neanderthal [Prüfer et al., 2017] and red pluses denote the CSFS conditioned on the high coverage Denisovan [Meyer et al., 2012].

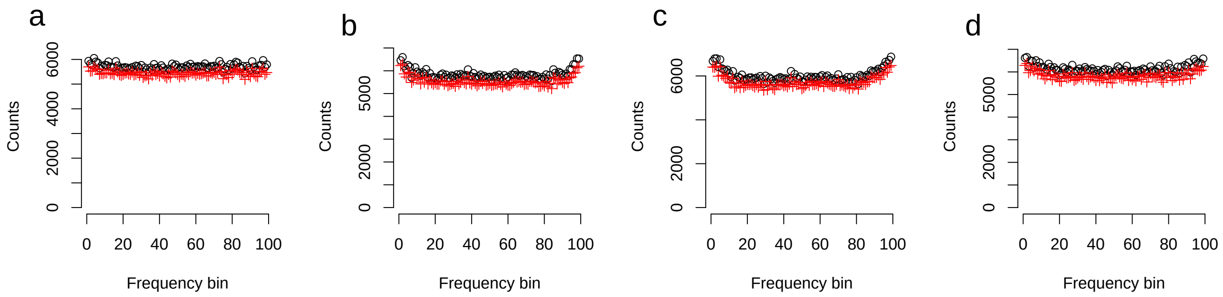

Figure S20: Model A.2: Gene flow from the modern human ancestor branch into the African branch after the out of Africa event. See section S4.2 for a detailed description. Panels show different amounts of gene flow into the African branch (a) 1% (b) 5% (c) 10% (d) 20%. Black circles denote the CSFS conditioned on the Vindija Neanderthal [Prüfer et al., 2017] and red pluses denote the CSFS conditioned on the high coverage Denisovan [Meyer et al., 2012].

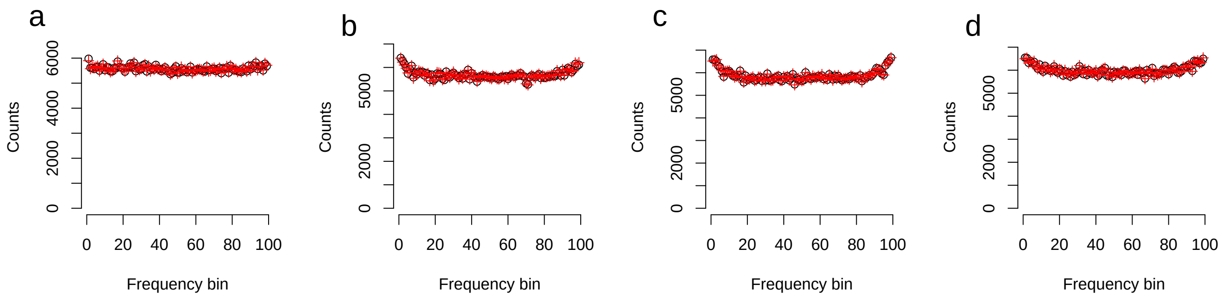

Figure S21: Model sA.2: We simulate the model in Figure S20, but only include the African, Neanderthal, and Denisovan branches (a) 1% (b) 5% (c) 10% (d) 20%. Black circles denote the CSFS conditioned on the Vindija Neanderthal [Prüfer et al., 2017] and red pluses denote the CSFS conditioned on the high coverage Denisovan [Meyer et al., 2012].

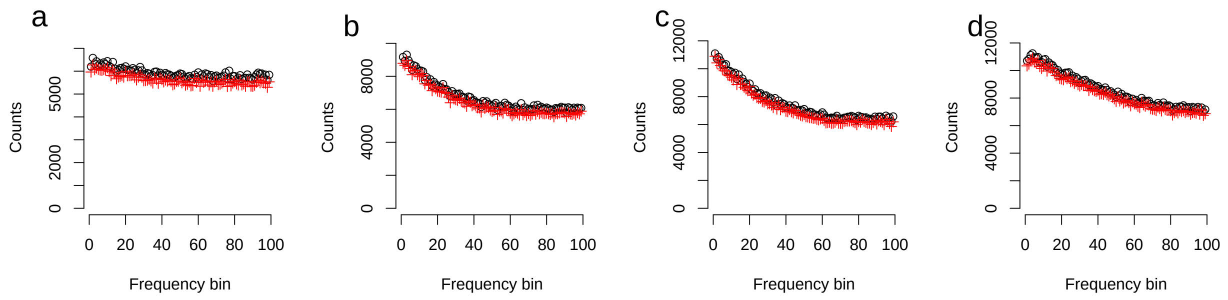

Figure S22: Model B.1: Gene flow from the archaic branch into the modern human ancestor prior to the out of Africa event. See section S4.3 for a detailed description. Panels show different amounts of gene flow into the modern human ancestor (a) 1% (b) 5% (c) 10% (d) 20%. Black circles denote the CSFS conditioned on the Vindija Neanderthal [Prüfer et al., 2017] and red pluses denote the CSFS conditioned on the high coverage Denisovan [Meyer et al., 2012].

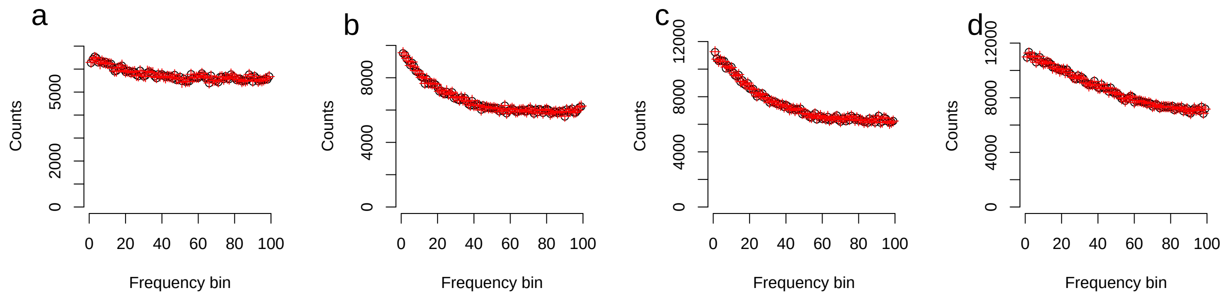

Figure S23: Model sB.1: We simulate the model in Figure S22, but only include the African, Neanderthal, and Denisovan branches (a) 1% (b) 5% (c) 10% (d) 20%. Black circles denote the CSFS conditioned on the Vindija Neanderthal [Prüfer et al., 2017] and red pluses denote the CSFS conditioned on the high coverage Denisovan [Meyer et al., 2012].

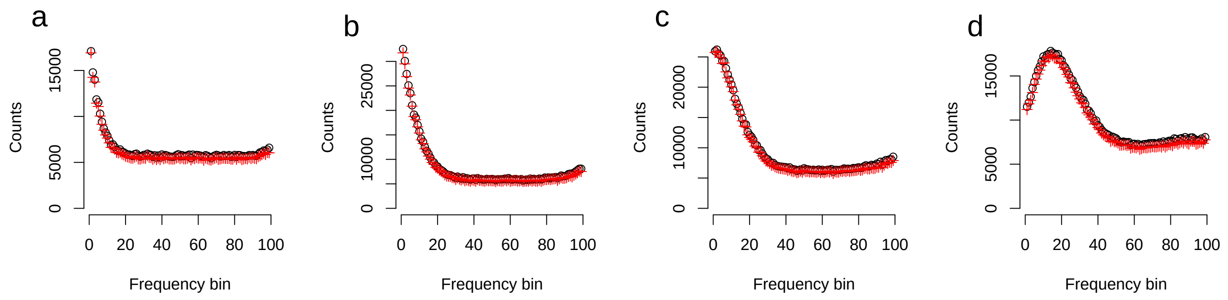

Figure S24: Model B.2: Gene flow from the archaic branch into the African branch after the out of Africa event. See section S4.4 for a detailed description. Panels show different amounts of gene flow into the African branch (a) 1% (b) 5% (c) 10% (d) 20%. Black circles denote the CSFS conditioned on the Vindija Neanderthal [Prüfer et al., 2017] and red pluses denote the CSFS conditioned on the high coverage Denisovan [Meyer et al., 2012].

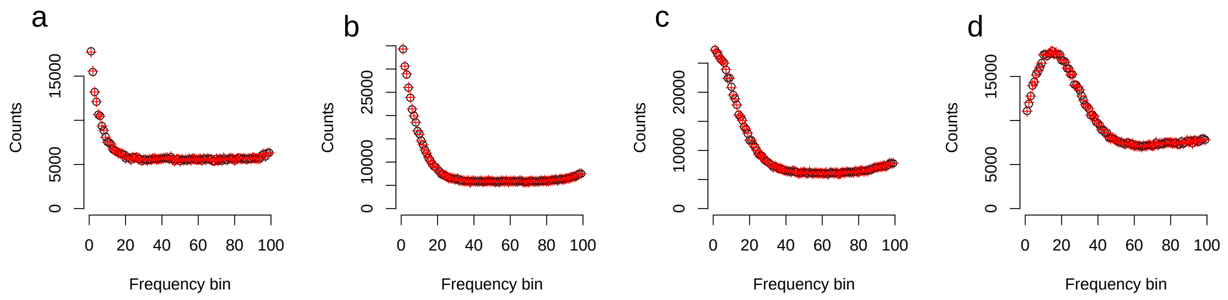

Figure S25: Model sB.2: We simulate the model in Figure S24, but only include the African, Neanderthal, and Denisovan branches (a) 1% (b) 5% (c) 10% (d) 20%. Black circles denote the CSFS conditioned on the Vindija Neanderthal [Prüfer et al., 2017] and red pluses denote the CSFS conditioned on the high coverage Denisovan [Meyer et al., 2012].

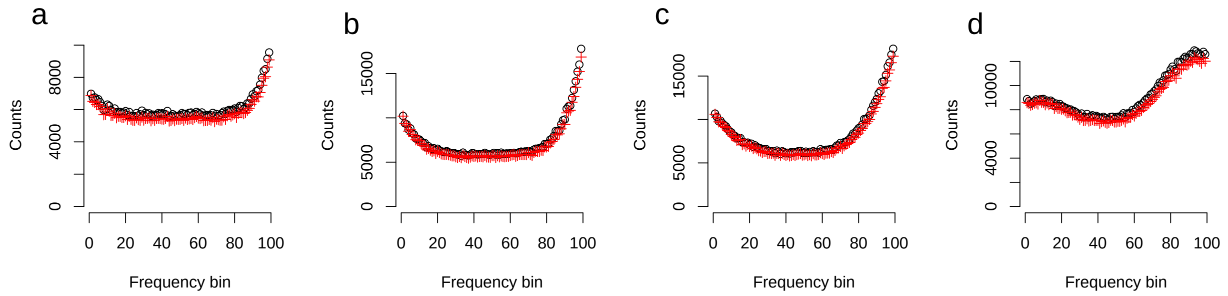

Figure S26: Model C.1: Gene flow from an unknown archaic branch into the modern human ancestor prior to the out of Africa event. See section S4.5 for a detailed description. Panels show different amounts of gene flow into the modern human ancestor (a) 1% (b) 5% (c) 10% (d) 20%. Black circles denote the CSFS conditioned on the Vindija Neanderthal [Prüfer et al., 2017] and red pluses denote the CSFS conditioned on the high coverage Denisovan [Meyer et al., 2012].

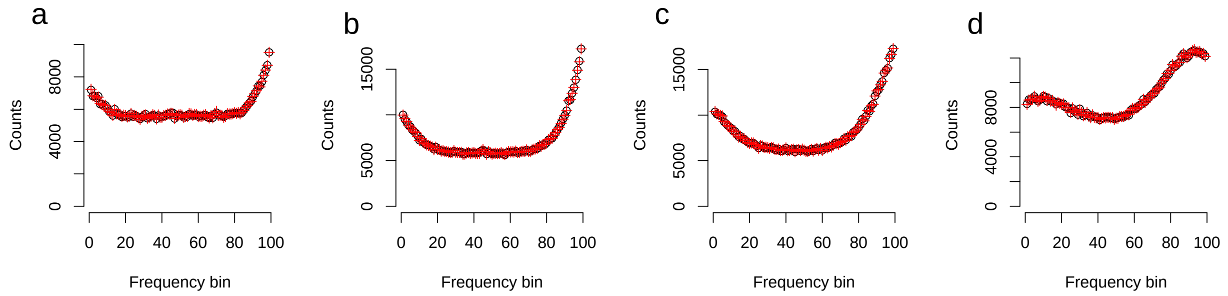

Figure S27: Model sC.1: We simulate the model in Figure S26, but only include the African, Neanderthal, and Denisovan branches (a) 1% (b) 5% (c) 10% (d) 20%. Black circles denote the CSFS conditioned on the Vindija Neanderthal [Prüfer et al., 2017] and red pluses denote the CSFS conditioned on the high coverage Denisovan [Meyer et al., 2012].

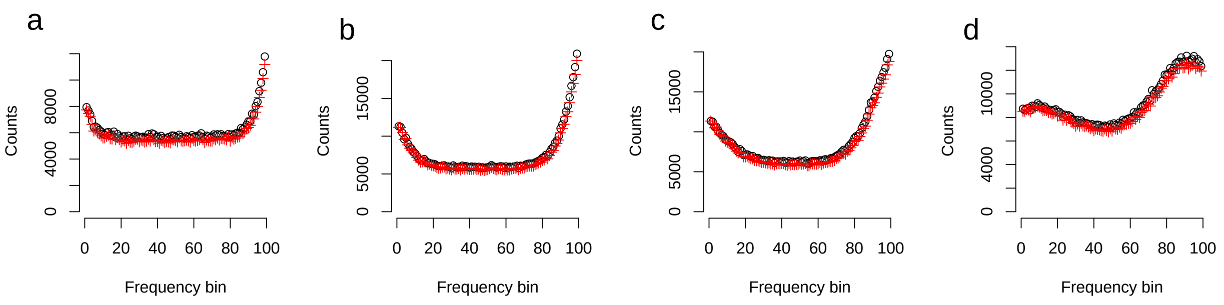

Figure S28: Model C.2: Gene flow from an unknown archaic branch into the African branch after the out of Africa event. See section S4.6 for a detailed description. Panels show different amounts of gene flow into the African branch (a) 1% (b) 5% (c) 10% (d) 20%. Black circles denote the CSFS conditioned on the Vindija Neanderthal [Prüfer et al., 2017] and red pluses denote the CSFS conditioned on the high coverage Denisovan [Meyer et al., 2012].

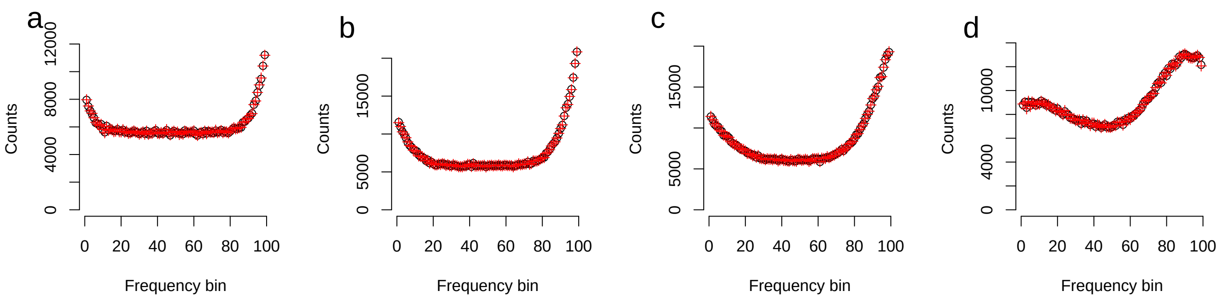

Figure S29: Model sC.2: We simulate the model in Figure S28, but only include the African, Neanderthal, and Denisovan branches (a) 1% (b) 5% (c) 10% (d) 20%. Black circles denote the CSFS conditioned on the Vindija Neanderthal [Prüfer et al., 2017] and red pluses denote the CSFS conditioned on the high coverage Denisovan [Meyer et al., 2012].

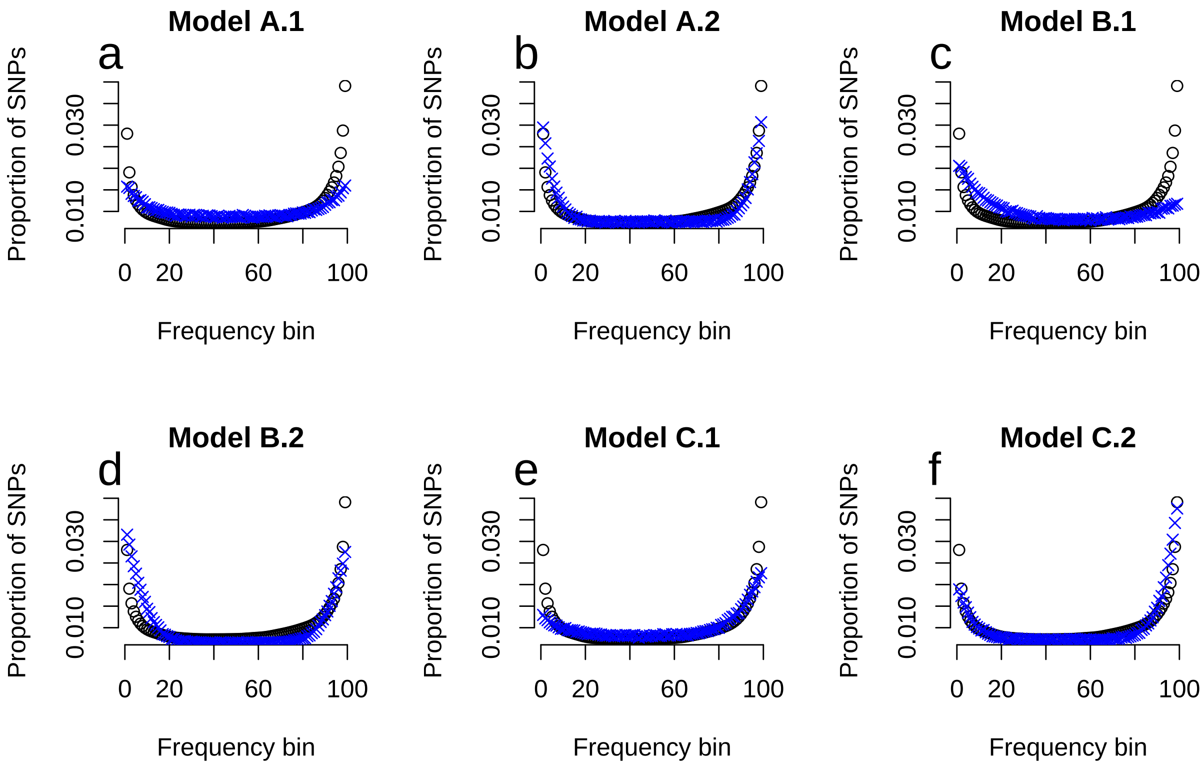

Figure S30: Simulations of the best fitting parameters for models A, B, C (Section S4). Blue crosses denote the simulated spectra and black circles denote the observed $CSFS_{YRI,N}$.

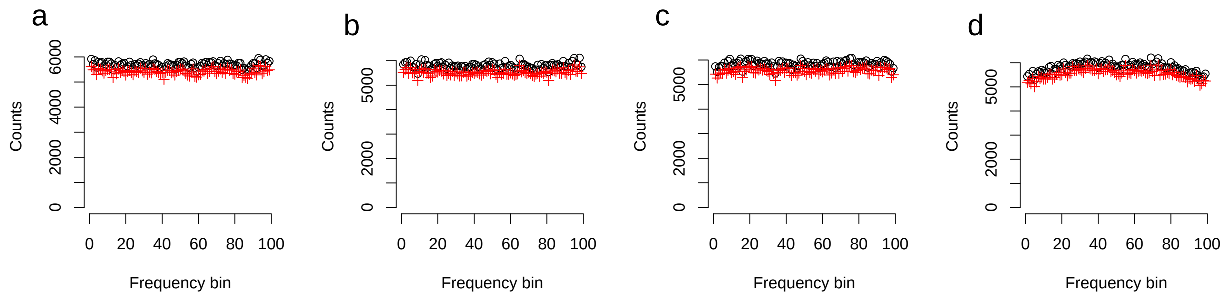

Figure S31: Model A.2 with a population size of $0.01N_a$ in the introgressing population. The panels represent different levels of gene flow (a) 1% (b) 5% (c) 10% (d) 20%. Black circles denote the CSFS conditioned on the Vindija Neanderthal [Prüfer et al., 2017] and red pluses denote the CSFS conditioned on the high coverage Denisovan [Meyer et al., 2012].

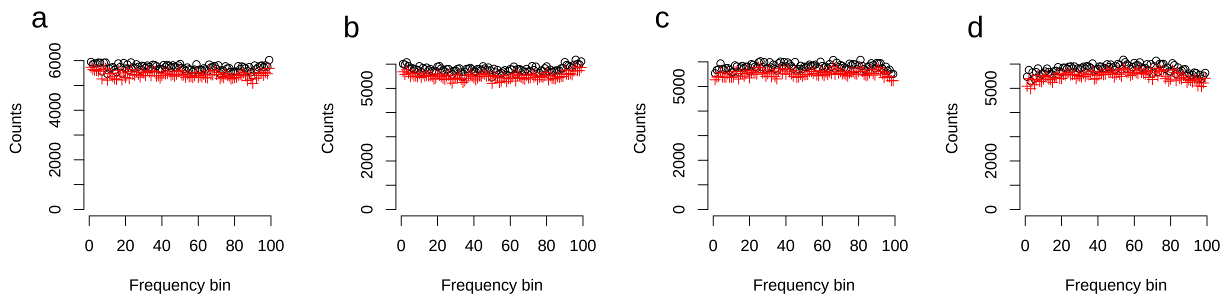

Figure S32: Model A.2 with a population size of $1 \times 10^{-4} N_a$ in the introgressing population. The panels represent different levels of gene flow (a) 1% (b) 5% (c) 10% (d) 20%. Black circles denote the CSFS conditioned on the Vindija Neanderthal [Prüfer et al., 2017] and red pluses denote the CSFS conditioned on the high coverage Denisovan [Meyer et al., 2012].

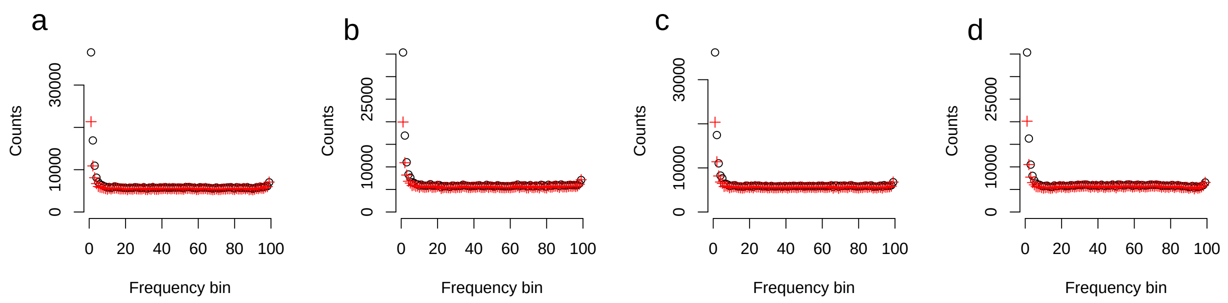

Figure S33: Model A.2 with a population size of $1 \times 10^{-4} N_a$ in the introgressing population and migration between CEU and YRI over the last 20Ky B.P. The panels represent different levels of gene flow (a) 1% (b) 5% (c) 10% (d) 20%. Black circles denote the CSFS conditioned on the Vindija Neanderthal [Prüfer et al., 2017] and red pluses denote the CSFS conditioned on the high coverage Denisovan [Meyer et al., 2012].

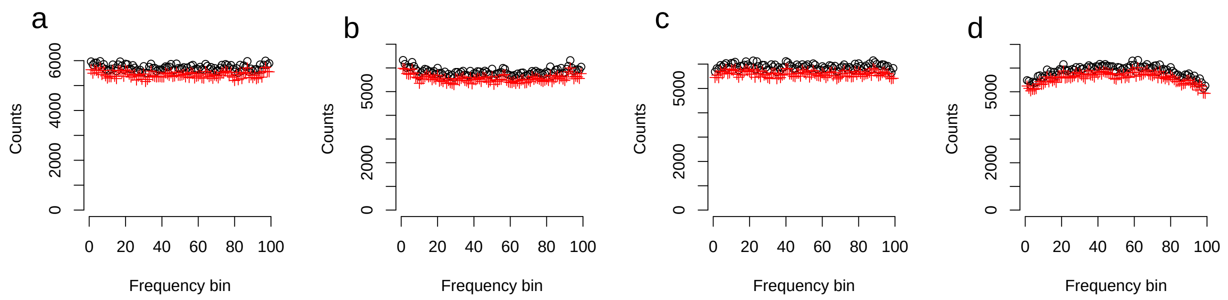

Figure S34: Model A.2 with a population size of $1 \times 10^{-5} N_a$ in the introgressing population, which branches off 200Ky B.P. The panels represent different levels of gene flow (a) 1% (b) 5% (c) 10% (d) 20%. Black circles denote the CSFS conditioned on the Vindija Neanderthal [Prüfer et al., 2017] and red pluses denote the CSFS conditioned on the high coverage Denisovan [Meyer et al., 2012].

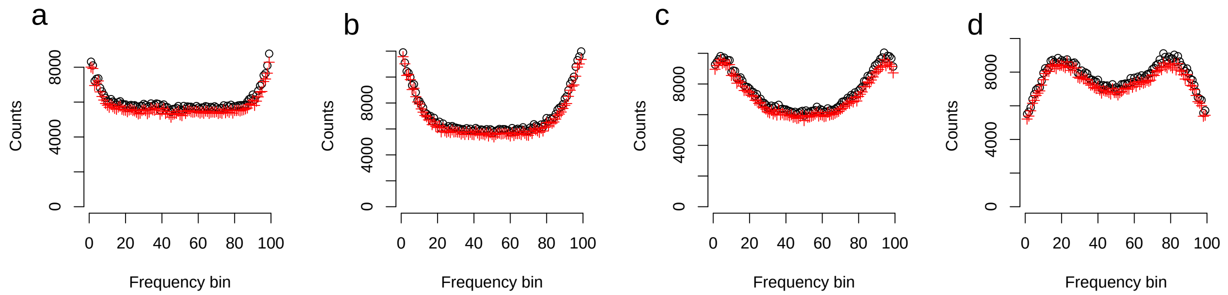

Figure S35: Model A.2 where the introgressing population splits at the same time as the archaic population (550Ky B.P.) with a population size of $0.01N_a$. The panels represent different levels of gene flow (a) 1% (b) 5% (c) 10% (d) 20%. Black circles denote the CSFS conditioned on the Vindija Neanderthal [Prüfer et al., 2017] and red pluses denote the CSFS conditioned on the high coverage Denisovan [Meyer et al., 2012].

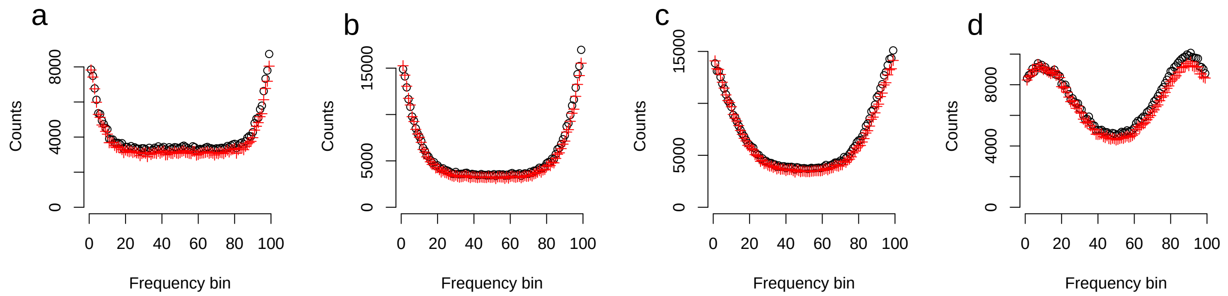

Figure S36: Model A.2 where the introgressing population splits at the same time as the archaic population, 765Ky B.P. This simulation shows that uncertainty in the split time between archaics and modern humans cannot explain the CSFS. The panels represent different levels of gene flow (a) 1% (b) 5% (c) 10% (d) 20%. Black circles denote the CSFS conditioned on the Vindija Neanderthal [Prüfer et al., 2017] and red pluses denote the CSFS conditioned on the high coverage Denisovan [Meyer et al., 2012].

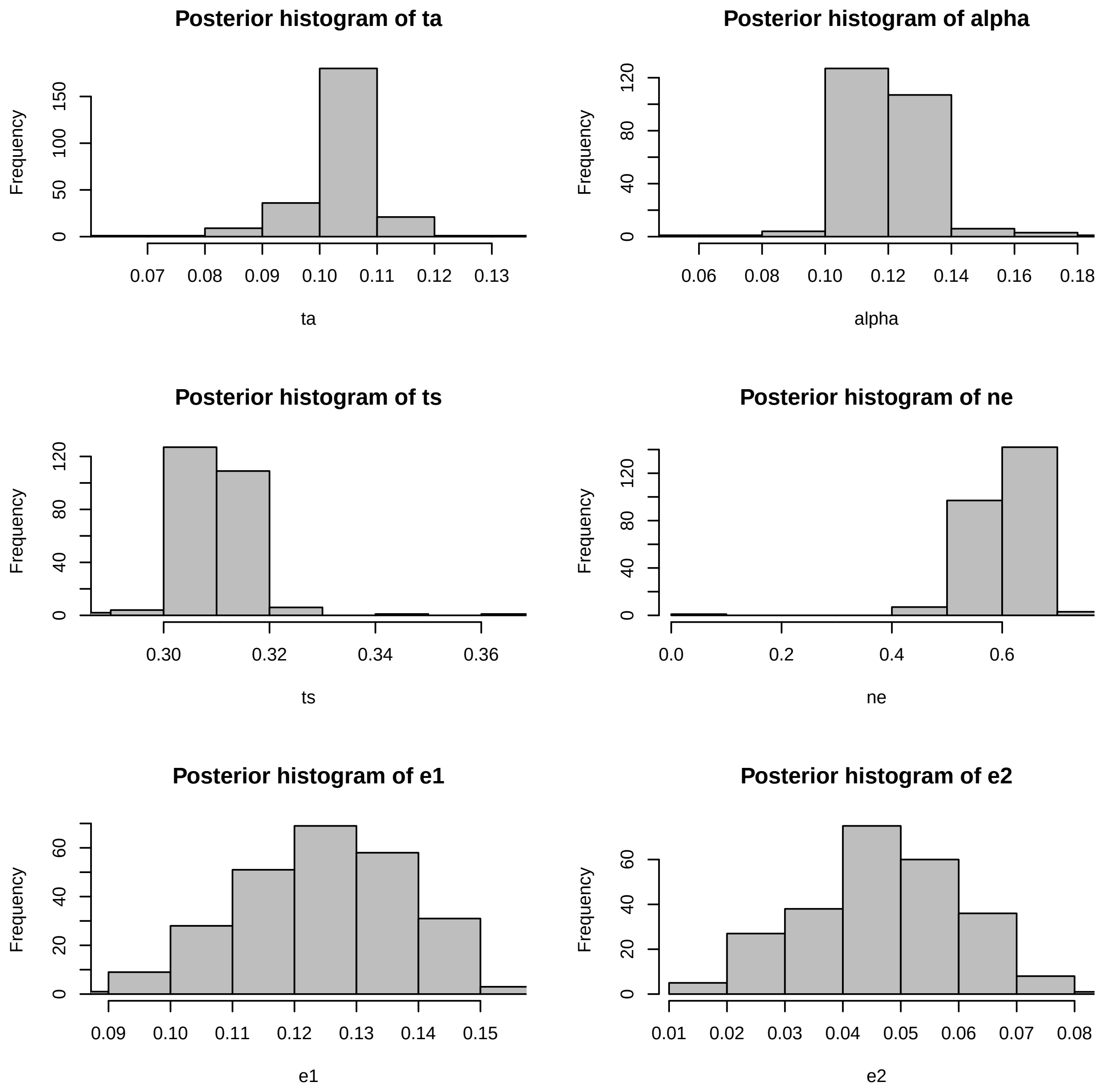

Figure S37: Parameter estimates using ABC for Model A.1 including ancestral misidentification (e1) and genotyping error in the archaic (e2). These estimates are made on a per-base pair rate. 'ta' is the time of admixture, scaled by $4N_a$, where $N_a$ is the ancestral population size (10,000). 'alpha' is the admixture fraction. 'ts' is the split time scaled by $4N_a$ and 'ne' is the effective population size of the introgressing lineage.

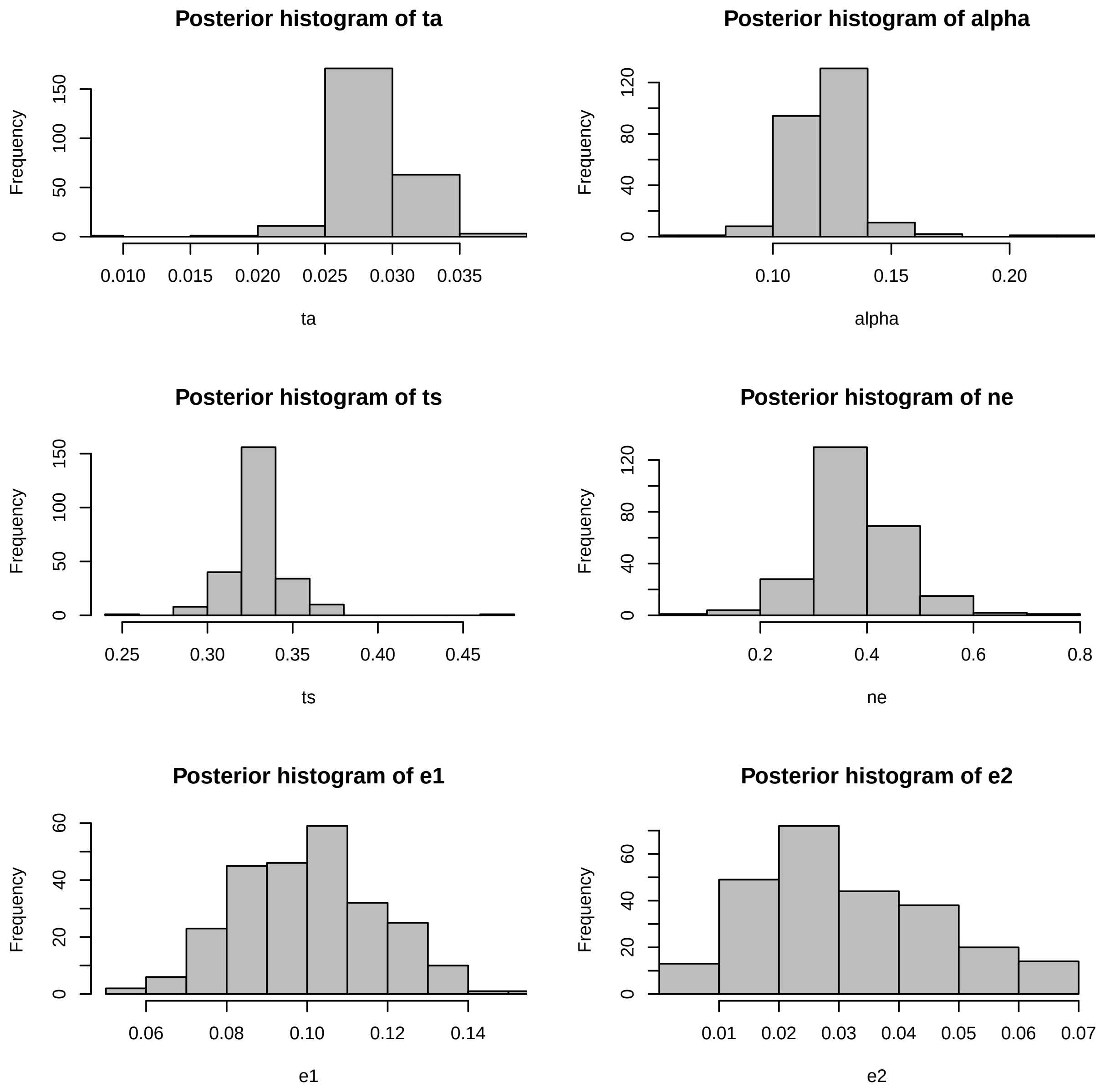

Figure S38: Parameter estimates using ABC for Model A.2 including ancestral misidentification (e1) and genotyping error in the archaic (e2). These estimates are made on a per-base pair rate. ‘ta’ is the time of admixture, scaled by $4N_a$, where $N_a$ is the ancestral population size (10,000). ‘alpha’ is the admixture fraction. ‘ts’ is the split time scaled by $4N_a$ and ‘ne’ is the effective population size of the introgressing lineage.

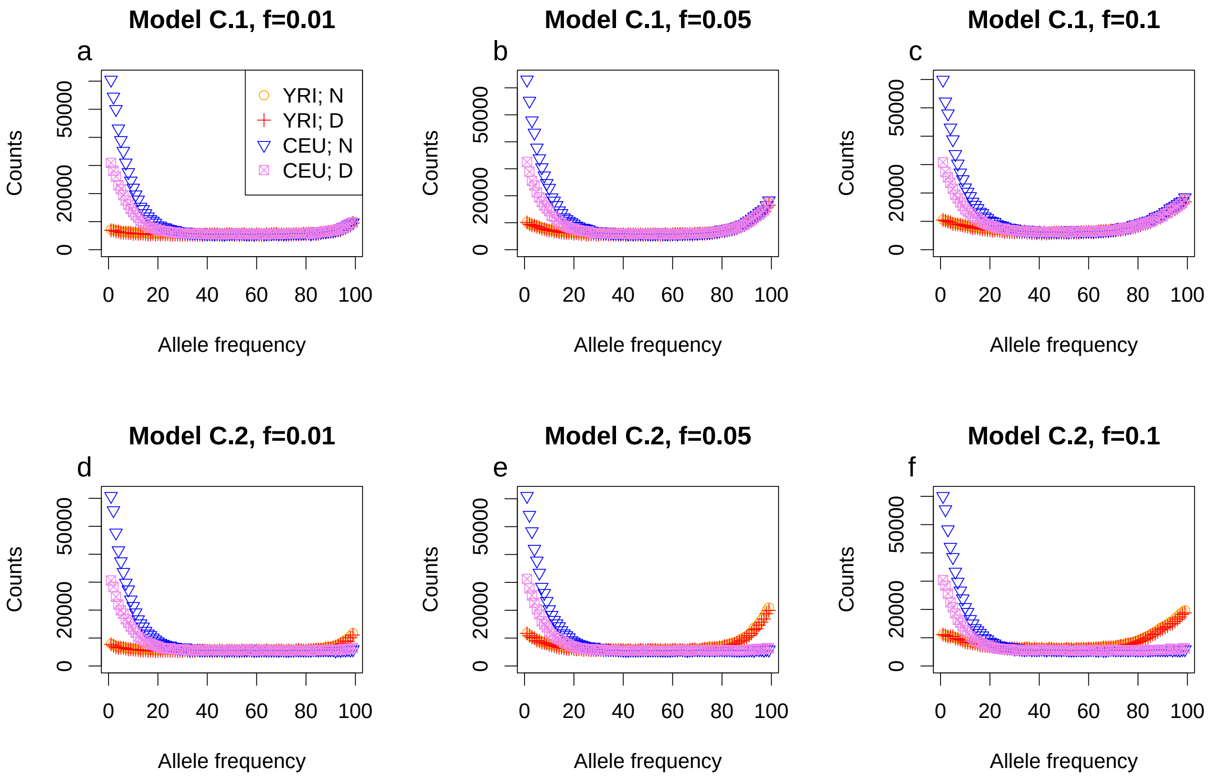

Figure S39: Marginalized joint CSFS of YRI and CEU from simulations. We contrast introgression into the modern human branch before the Out-of-Africa split (C.1 – top row) against introgression into the African branch specifically (C.2 – bottom row). We see that model C.1 produces an increase in high frequency derived alleles for both CEU and YRI, while C.2 only produces that signal in YRI.

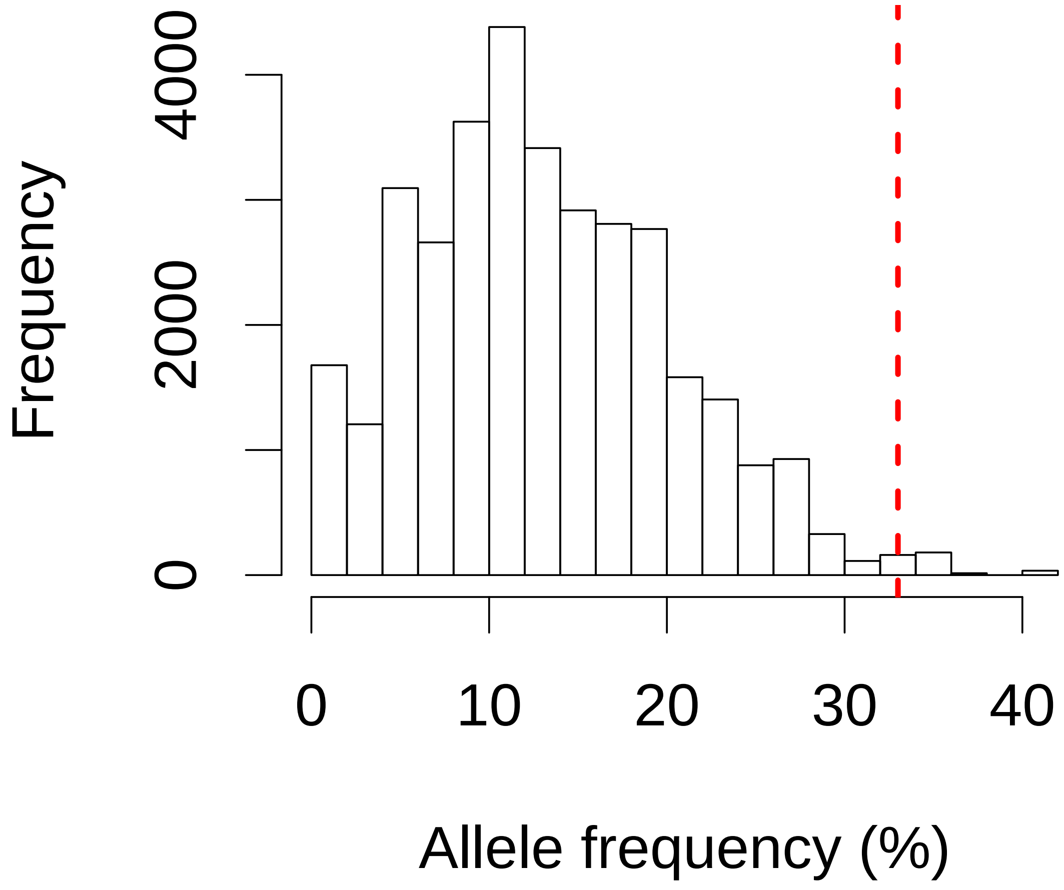

Figure S40: Distribution of allele frequencies for neutral archaic SNPs from model C with 13% introgression and an introgression time of 42Ky B.P. The red line denotes the 99% quantile, which corresponds to an allele frequency of 33%.

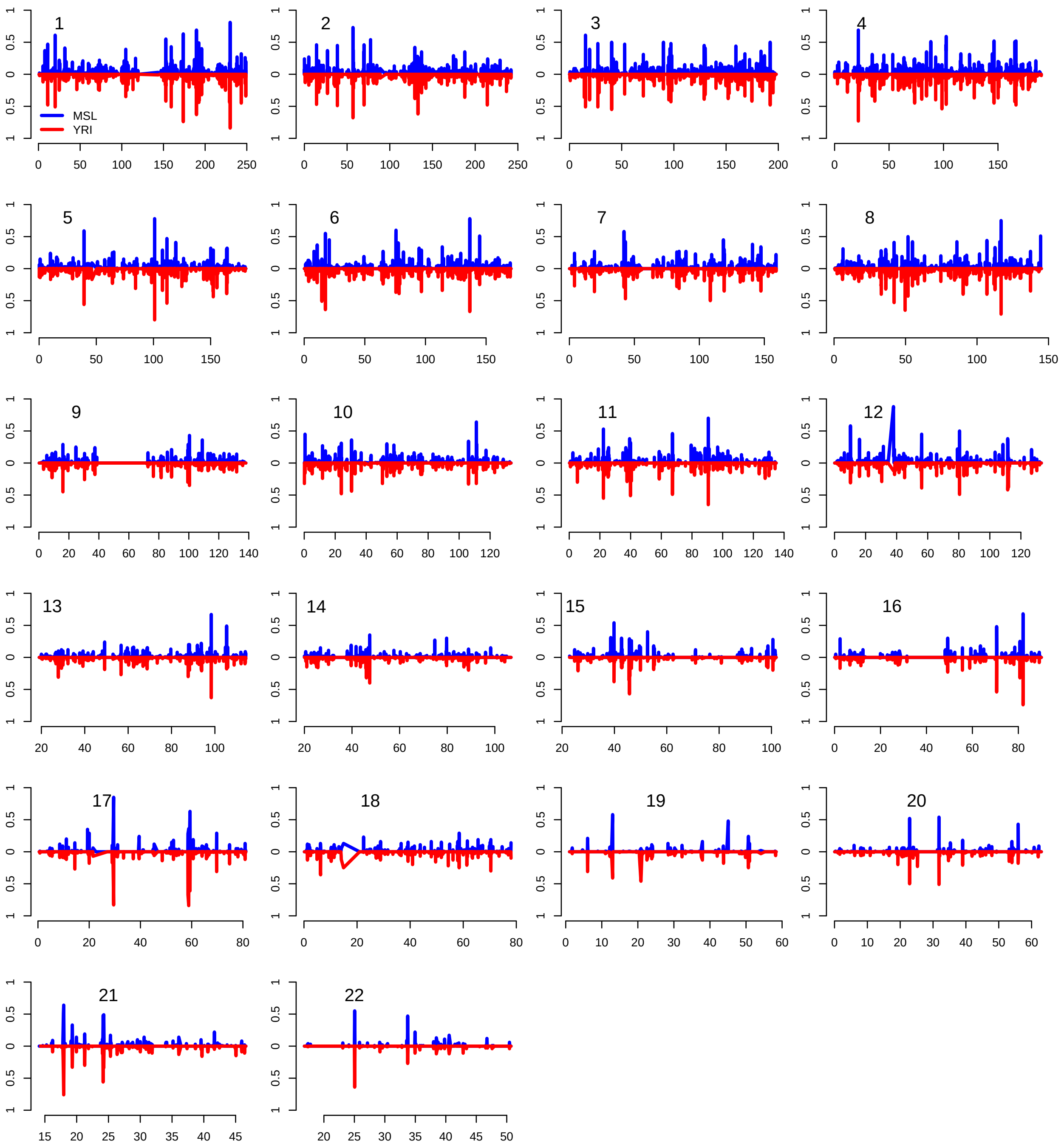

Figure S41: Archaic segment frequency map for MSL and YRI. Spearman's $\rho$ between the frequencies of the two maps is 0.71 ($p < 2 \times 10^{-16}$).

| Model | Description |
|---|---|
| A.1 | Gene flow into the modern human ancestor from a lineage that splits from the modern human ancestor 6K generations ago |
| A.2 | Gene flow into the African branch from a lineage that splits from the modern human ancestor 6K generations ago |
| A.2 contraction | Model A.2 with an effective population size of 100 in the introgressing branch |
| A.2 contraction 2 | Model A.2 with an effective population size of 1 in the introgressing branch |
| A.2 contraction 2 CEU-YRI migration | Model A.2 with an effective population size of 100 in the introgressing branch and migration between YRI and CEU over the last 20Kya |
| A.2 contraction 3 | Model A.2 with an effective population size of 0.1 in the introgressing branch |
| A.2 trifurcation | Model A.2 with the introgressing lineage splitting off the same time as the archaic branch ( 550Kya) and an effective population size of 100 |
| B.1 | Gene flow into the modern human ancestor from a lineage that splits from the archaic branch (ancestor of Denisovan and Neanderthal) 16K generations ago |
| B.2 | Gene flow into the African branch from a lineage that splits from the archaic branch (ancestor of Denisovan and Neanderthal) 16K generations ago |
| Baseline with errors | A model including the major demographic events (see Figure 1) with ancestral misidentification error and archaic sequencing error |
| C.1 | Gene flow into the modern human ancestor from a lineage that splits from the modern human ancestor 24K generations ago |
| C.2 | Gene flow into the African branch from a lineage that splits from the modern human ancestor 24K generations ago |
| Continuous Migration | Continuous bidirectional migration between the African branch and a lineage that splits 174Kya over the past 174Kya |
| Continuous Migration 2 | Continuous bidirectional migration between the African branch and a lineage that splits 350Kya over the past 350Kya |
| sA.1 | A simplified version of model A.1 simulating on the African Neanderthal and Denisovan branches |
| sA.2 | A simplified version of model A.2 simulating on the African Neanderthal and Denisovan branches |
| sB.1 | A simplified version of model B.1 simulating on the African Neanderthal and Denisovan branches |
| sB.2 | A simplified version of model B.2 simulating on the African Neanderthal and Denisovan branches |
| sC.1 | A simplified version of model C.1 simulating on the African Neanderthal and Denisovan branches |
| sC.2 | A simplified version of model C.2 simulating on the African Neanderthal and Denisovan branches |
| SIM004 | The model from Pruefer et al 2017; including gene flow from the super archaic into the Denisovan; modern humans into Neanderthals; Neanderthal into Europeans; and a small contribution from the Neanderthal directly to the African branch |
| SIM005 | The model from Pruefer et al 2017; with migration between CEU and YRI over the last 20;000 years |
| SIM006 | The model from Pruefer et al 2017; with a 5x population expansion in the modern human-archaic ancestor |
| SIM007 | The model from Pruefer et al 2017; with a 5x population bottleneck in the modern human-archaic ancestor |
| SIM008 | The model from Pruefer et al 2017; with a population that branches off from and rejoins the modern human-archaic ancestor |
| SIM009 | The model from Pruefer et al 2017; with a population that branches off from the modern human-archaic ancestor and includes bidirectional migration between the lineages |
| SIM010 | The model from Pruefer et al 2017; with a population that branches off from and rejoins the modern human-archaic ancestor with an effective population size of 100 |
| SIM012 | The ancestral structure model from Yang et al 2012 |
| SIM013 | The ancestral structure model from Sankararaman et al 2012 |

Table S1: Description of the models examined in this work.

| **Model** | **Ancestral misidentification rate (e1)** | **Genotype error in archaic (e2)** | **$\Delta\mathcal{LL}$ (N)** |
|---|---|---|---|
| Baseline model with errors | 0.03 | 0.03 | -40266 |
| Baseline model with errors | 0.03 | 0.15 | -147596 |
| Baseline model with errors | 0.03 | 0.3 | -269099 |
| Baseline model with errors | 0.03 | 0 | -21487 |
| Baseline model with errors | 0.15 | 0.03 | -6356 |
| Baseline model with errors | 0.15 | 0.15 | -57467 |
| Baseline model with errors | 0.15 | 0.3 | -141569 |
| Baseline model with errors | 0.15 | 0 | -7897 |
| Baseline model with errors | 0.3 | 0.03 | -39044 |
| Baseline model with errors | 0.3 | 0.15 | -42293 |
| Baseline model with errors | 0.3 | 0.3 | -92908 |
| Baseline model with errors | 0.3 | 0 | -54731 |
| Baseline model with errors | 0 | 0.03 | -94699 |
| Baseline model with errors | 0 | 0.15 | -215818 |
| Baseline model with errors | 0 | 0.3 | -345684 |
| Baseline model with errors | 0 | 0 | -68887 |

Table S2: We simulated data from the Prüfer *et al.* 2017 model and added in ancestral misidentification error and genotyping error in the archaic. $\Delta\mathcal{LL}$ values come from comparing to model C. **N** refers to the fit of the simulated model compared to $CSFS_{YRI,N}$, the site frequency spectrum in Yoruba conditional on observing a derived allele in the Vindija Neanderthal. **D** refers to $CSFS_{YRI,D}$, the same spectrum conditional on observing a derived allele in the Denisovan.

| **Model** | **Admixture frac** | **$\Delta\mathcal{LL}$ (N)** | **$\Delta\mathcal{LL}$ (D)** |
|---|---|---|---|
| SIM004 | 0.00001 | -76327 | -73841 |
| SIM004 | 0.00015 | -73248 | -72149 |
| SIM004 | 0.001 | -62883 | -66355 |
| SIM004 | 0.01 | -140008 | -77851 |
| SIM004 | 0.02 | -232317 | -111044 |
| SIM005 | 0 | -68973 | -59745 |
| SIM006 | 0 | -79970 | -77013 |
| SIM007 | 0 | -76740 | -71665 |
| SIM008 | 0 | -80141 | -76350 |
| SIM009 | 0 | -77188 | -74001 |
| SIM010 | 0 | -80590 | -77755 |
| SIM012 | 0 | -78358 | -64205 |
| SIM013 | 0 | -133592 | -79971 |
| SIM014 | 0 | -76738 | -73585 |
| SIM015 | 0 | -77281 | -74304 |
| Continuous Migration | 0 | -41582 | -41233 |
| Continuous Migration 2 | 0 | -10263 | -12825 |

Table S3: Model fits for null models including structure, and departures from panmixia in the MH ancestor.

| **Model** | **Admixture frac** | $\Delta\mathcal{LL}$ **(N)** | $\Delta\mathcal{LL}$ **(D)** |
|---|---|---|---|
| Model A.1 | 0 | -76485 | -73336 |
| Model A.1 | 0.01 | -74330 | -71022 |
| Model A.1 | 0.05 | -69637 | -66718 |
| Model A.1 | 0.1 | -69422 | -66628 |
| Model A.1 | 0.2 | -68072 | -65192 |
| Model A.1 | 0.3 | -73853 | -71014 |
| Model A.1 | 0.4 | -75826 | -72684 |
| Model A.1 | 0.5 | -77667 | -74332 |
| Model A.2 | 0 | -76051 | -72478 |
| Model A.2 | 0.01 | -61468 | -59062 |
| Model A.2 | 0.05 | -68762 | -65950 |
| Model A.2 | 0.1 | -70774 | -67646 |
| Model A.2 | 0.2 | -76970 | -73358 |
| Model A.2 | 0.3 | -78398 | -74864 |
| Model A.2 | 0.4 | -75826 | -72684 |
| Model A.2 | 0.5 | -77667 | -74332 |
| Model B.1 | 0 | -76145 | -73595 |
| Model B.1 | 0.01 | -76329 | -74226 |
| Model B.1 | 0.05 | -80157 | -80135 |
| Model B.1 | 0.1 | -89212 | -89070 |
| Model B.1 | 0.2 | -97318 | -95547 |
| Model B.1 | 0.3 | -95664 | -92503 |
| Model B.1 | 0.4 | -97898 | -93205 |
| Model B.1 | 0.5 | -100880 | -97505 |
| Model B.2 | 0 | -74598 | -71684 |
| Model B.2 | 0.01 | -58683 | -63534 |
| Model B.2 | 0.05 | -119042 | -131895 |
| Model B.2 | 0.1 | -142358 | -153703 |
| Model B.2 | 0.2 | -153758 | -153893 |
| Model B.2 | 0.3 | -187791 | -180909 |
| Model B.2 | 0.4 | -220041 | -211750 |
| Model B.2 | 0.5 | -237342 | -237537 |
| Model C.1 | 0 | -76983 | -74461 |
| Model C.1 | 0.01 | -37989 | -35735 |
| Model C.1 | 0.05 | -5293 | -4283 |
| Model C.1 | 0.1 | -7722 | -6535 |
| Model C.1 | 0.2 | -39705 | -36102 |
| Model C.1 | 0.3 | -82461 | -77298 |
| Model C.1 | 0.4 | -119884 | -114146 |
| Model C.1 | 0.5 | -135067 | -129395 |
| Model C.2 | 0 | -74166 | -71302 |
| Model C.2 | 0.01 | -27535 | -25768 |
| Model C.2 | 0.05 | 0 | 0 |
| Model C.2 | 0.1 | -6513 | -5465 |
| Model C.2 | 0.2 | -40939 | -37021 |
| Model C.2 | 0.3 | -90789 | -84908 |
| Model C.2 | 0.4 | -128417 | -121805 |
| Model C.2 | 0.5 | -143948 | -137907 |

Table S4: Model fits for alternate models including admixture from other lineages.

| Model | Admixture frac | $\Delta\mathcal{LL}$ (N) | $\Delta\mathcal{LL}$ (D) |
|---|---|---|---|
| Model sA.1 | 0 | -77663 | -74836 |
| Model sA.1 | 0.01 | -77073 | -73515 |
| Model sA.1 | 0.05 | -68729 | -65993 |
| Model sA.1 | 0.1 | -69547 | -66381 |
| Model sA.1 | 0.2 | -69401 | -66540 |
| Model sA.1 | 0.3 | -73397 | -70053 |
| Model sA.1 | 0.4 | -73311 | -70225 |
| Model sA.1 | 0.5 | -80641 | -77307 |
| Model sA.2 | 0 | -75448 | -71842 |
| Model sA.2 | 0.01 | -73554 | -70830 |
| Model sA.2 | 0.05 | -66347 | -63575 |
| Model sA.2 | 0.1 | -63292 | -61303 |
| Model sA.2 | 0.2 | -66388 | -64169 |
| Model sA.2 | 0.3 | -71185 | -68382 |
| Model sA.2 | 0.4 | -75456 | -72415 |
| Model sA.2 | 0.5 | -77676 | -74768 |
| Model sB.1 | 0 | -76720 | -73376 |
| Model sB.1 | 0.01 | -76002 | -73286 |
| Model sB.1 | 0.05 | -78327 | -77557 |
| Model sB.1 | 0.1 | -91956 | -91172 |
| Model sB.1 | 0.2 | -100780 | -98156 |
| Model sB.1 | 0.3 | -98971 | -95341 |
| Model sB.1 | 0.4 | -100061 | -94648 |
| Model sB.1 | 0.5 | -106902 | -99152 |
| Model sB.2 | 0 | -75851 | -72753 |
| Model sB.2 | 0.01 | -59875 | -63472 |
| Model sB.2 | 0.05 | -131901 | -137205 |
| Model sB.2 | 0.1 | -156244 | -157522 |
| Model sB.2 | 0.2 | -161724 | -157291 |
| Model sB.2 | 0.3 | -190116 | -180659 |
| Model sB.2 | 0.4 | -228504 | -216002 |
| Model sB.2 | 0.5 | -257411 | -242870 |
| Model sC.1 | 0 | -77344 | -73884 |
| Model sC.1 | 0.01 | -36504 | -34910 |
| Model sC.1 | 0.05 | -5128 | -4854 |
| Model sC.1 | 0.1 | -8594 | -7840 |
| Model sC.1 | 0.2 | -40173 | -37336 |
| Model sC.1 | 0.3 | -83519 | -79010 |
| Model sC.1 | 0.4 | -118962 | -113399 |
| Model sC.1 | 0.5 | -134253 | -128664 |
| Model sC.2 | 0 | -77341 | -73731 |
| Model sC.2 | 0.01 | -27255 | -25989 |
| Model sC.2 | 0.05 | 468 | 15 |
| Model sC.2 | 0.1 | -5550 | -5185 |
| Model sC.2 | 0.2 | -38750 | -35986 |
| Model sC.2 | 0.3 | -87765 | -83089 |
| Model sC.2 | 0.4 | -126740 | -121280 |
| Model sC.2 | 0.5 | -141187 | -135455 |

Table S5: Model fits for alternate models using a simplified demography. See Section S4 for more information.

| **Model** | **Admixture frac** | $\Delta\mathcal{LL}$ **(N)** | $\Delta\mathcal{LL}$ **(D)** |
|---|---|---|---|
| Model A.2 contraction2 | 0.01 | -75254 | -72244 |
| Model A.2 contraction2 | 0.05 | -70771 | -67645 |
| Model A.2 contraction2 | 0.1 | -81146 | -77265 |
| Model A.2 contraction2 | 0.2 | -86625 | -83723 |
| Model A.2 contraction2 | 0.3 | -72311 | -69370 |
| Model A.2 contraction2 | 0.4 | -48472 | -47207 |
| Model A.2 contraction2 | 0.5 | -26220 | -26562 |
| Model A.2 contraction2 | 0 | -76424 | -73021 |
| Model A.2 contraction2 CEU-YRI migration | 0.01 | -69128 | -58417 |
| Model A.2 contraction2 CEU-YRI migration | 0.05 | -64879 | -56175 |
| Model A.2 contraction2 CEU-YRI migration | 0.1 | -68617 | -60630 |
| Model A.2 contraction2 CEU-YRI migration | 0.2 | -73427 | -65167 |
| Model A.2 contraction2 CEU-YRI migration | 0.3 | -67430 | -57793 |
| Model A.2 contraction2 CEU-YRI migration | 0.4 | -49946 | -39325 |
| Model A.2 contraction2 CEU-YRI migration | 0.5 | -41164 | -27819 |
| Model A.2 contraction2 CEU-YRI migration | 0 | -67856 | -58139 |
| Model A.2 contraction3 | 0.01 | -73682 | -70561 |
| Model A.2 contraction3 | 0.05 | -70574 | -68219 |
| Model A.2 contraction3 | 0.1 | -78825 | -75131 |
| Model A.2 contraction3 | 0.2 | -93451 | -88494 |
| Model A.2 contraction3 | 0.3 | -81440 | -78479 |
| Model A.2 contraction3 | 0.4 | -55167 | -53834 |
| Model A.2 contraction3 | 0.5 | -28170 | -28008 |
| Model A.2 contraction3 | 0 | -75734 | -73148 |
| Model A.2 contraction | 0.01 | -74718 | -71715 |
| Model A.2 contraction | 0.05 | -73414 | -70446 |
| Model A.2 contraction | 0.1 | -79103 | -75184 |
| Model A.2 contraction | 0.2 | -89882 | -85375 |
| Model A.2 contraction | 0.3 | -73552 | -71187 |
| Model A.2 contraction | 0.4 | -47006 | -46154 |
| Model A.2 contraction | 0.5 | -27072 | -26996 |
| Model A.2 contraction | 0 | -77686 | -73900 |
| Model A.2 trifurcation | 0.01 | -40988 | -40070 |
| Model A.2 trifurcation | 0.05 | -9819 | -12247 |
| Model A.2 trifurcation | 0.1 | -11416 | -13852 |
| Model A.2 trifurcation | 0.2 | -72025 | -68212 |
| Model A.2 trifurcation | 0.3 | -101322 | -97129 |
| Model A.2 trifurcation | 0.4 | -113989 | -109130 |
| Model A.2 trifurcation | 0.5 | -42019 | -41136 |
| Model A.2 trifurcation | 0 | -15355 | -16825 |

Table S6: Model fits for variations of model A.

| **Parameter** | **Value** | **95% HPD** | **Pop** |
|---|---|---|---|
| Branch split time ($t_s$) | $62.5 \times 10^4$ | $36.5 \times 10^4 - 102 \times 10^4$ | ESN |
| Admixture fraction ($\alpha$) | 0.09 | 0.02-0.19 | ESN |
| Admixture time ($t_a$) | $3.8 \times 10^4$ | $0 - 10.8 \times 10^4$ | ESN |
| Eff. population size ($N_e$) | $2.3 \times 10^4$ | $2.2 \times 10^4 - 2.5 \times 10^4$ | ESN |
| Branch split time ($t_s$) | $66 \times 10^4$ | $41.5 \times 10^4 - 99.9 \times 10^4$ | GWD |
| Admixture fraction ($\alpha$) | 0.12 | 0.06-0.19 | GWD |
| Admixture time ($t_a$) | $5.9 \times 10^4$ | $3.2 \times 10^4 - 10.4 \times 10^4$ | GWD |
| Eff. population size ($N_e$) | $2.4 \times 10^4$ | $2.3 \times 10^4 - 2.5 \times 10^4$ | GWD |
| Branch split time ($t_s$) | $62.4 \times 10^4$ | $29.1 \times 10^4 - 113.7 \times 10^4$ | LWK |
| Admixture fraction ($\alpha$) | 0.18 | 0.09-0.31 | LWK |
| Admixture time ($t_a$) | $5 \times 10^4$ | $1.1 \times 10^4 - 13.3 \times 10^4$ | LWK |
| Eff. population size ($N_e$) | $2.4 \times 10^4$ | $2.2 \times 10^4 - 2.6 \times 10^4$ | LWK |
| Branch split time ($t_s$) | $61.1 \times 10^4$ | $39 \times 10^4 - 88.4 \times 10^4$ | MSL |
| Admixture fraction ($\alpha$) | 0.09 | 0.05-0.16 | MSL |
| Admixture time ($t_a$) | $3.7 \times 10^4$ | $0.82 \times 10^4 - 9.8 \times 10^4$ | MSL |
| Eff. population size ($N_e$) | $2.6 \times 10^4$ | $2.4 \times 10^4 - 2.8 \times 10^4$ | MSL |
| Branch split time ($t_s$) | $62.5 \times 10^4$ | $36 \times 10^4 - 97.5 \times 10^4$ | YRI |
| Admixture fraction ($\alpha$) | 0.11 | 0.045-0.19 | YRI |
| Admixture time ($t_a$) | $4.3 \times 10^4$ | $0.6 \times 10^4 - 12.4 \times 10^4$ | YRI |
| Eff. population size ($N_e$) | $2.5 \times 10^4$ | $2.3 \times 10^4 - 2.7 \times 10^4$ | YRI |

Table S7: Best fitting parameter values for all populations using approximate Bayesian computation. Times are expressed in years. See Section S6.3 for details on the conversion from generations to years.

| Model | $p$-value |
|---|---|
| A.1 | $6.7 \times 10^{-16}$ |
| A.2 | $3.2 \times 10^{-9}$ |
| B.1 | $1.4 \times 10^{-11}$ |
| B.2 | $6.7 \times 10^{-16}$ |
| C.1 | 0.003 |
| C.2 | $9.5 \times 10^{-6}$ |

Table S8: P-values of a test of goodness of fit for the best fitting parameters for each class of demographic models. We computed the residuals comparing the empirical CSFS to the CSFS simulated under the best-fitting parameters for each model. We assessed goodness-of-fit of the model by testing if the residuals are approximately normally distributed using the approach outlined in Section S1.5. We computed goodness-of-fit restricting to derived allele counts ranging between 6 and 95. While all the compared are formally rejected (likely due to additional unmodeled complexities in the true underlying demographic model), we find that model C has substantially higher p-values than the other demographic models consistent with the observations in the main text.
